# Design of 8-mer Peptides that Block *Clostridioides difficile* Toxin A in Intestinal Cells

**DOI:** 10.1101/2023.01.10.523493

**Authors:** Sudeep Sarma, Carly M. Catella, Ellyce T. San Pedro, Xingqing Xiao, Deniz Durmusoglu, Stefano Menegatti, Nathan Crook, Scott T. Magness, Carol K. Hall

## Abstract

*Clostridioides difficile* (*C. diff*.) is a bacterium that causes severe diarrhea and inflammation of the colon. The pathogenicity of *C. diff*. infection is derived from two major toxins, toxins A (TcdA) and B (TcdB). Peptide inhibitors that can be delivered to the gut to inactivate these toxins are an attractive therapeutic strategy. In this work, we present a new approach that combines a *pep*tide *b*inding *d*esign algorithm (PepBD), molecular-level simulations, rapid screening of candidate peptides for toxin binding, a primary human cell-based assay, and surface plasmon resonance (SPR) measurements to develop peptide inhibitors that block the glucosyltransferase activity of TcdA by targeting its glucosyltransferase domain (GTD). Using PepBD and explicit-solvent molecular dynamics simulations, we identified seven candidate peptides, SA1-SA7. These peptides were selected for specific TcdA GTD binding through a custom solid-phase peptide screening system, which eliminated the weaker inhibitors SA5-SA7. The efficacies of SA1-SA4 were then tested using a trans-epithelial electrical resistance (TEER) assay on monolayers of the human gut epithelial culture model. One peptide, SA1, was found to block TcdA toxicity in primary-derived human jejunum (small intestinal) and colon (large intestinal) epithelial cells. SA1 bound TcdA with a K_D_ of 56.1 ± 29.8 nM as measured by surface plasmon resonance (SPR).

**Significance Statement:** Infections by *Clostridioides difficile*, a bacterium that targets the large intestine (colon), impact a significant number of people worldwide. Bacterial colonization is mediated by two exotoxins: toxins A and B. Short peptides that can inhibit the biocatalytic activity of these toxins represent a promising strategy to prevent and treat *C. diff*. infection. We describe an approach that combines a *Peptide B*inding *D*esign (PepBD) algorithm, molecular-level simulations, a rapid screening assay to evaluate peptide:toxin binding, a primary human cell-based assay, and surface plasmon resonance (SPR) measurements to develop peptide inhibitors that block Toxin A in small intestinal and colon epithelial cells. Importantly, our designed peptide, SA1, bound toxin A with nanomolar affinity and blocked toxicity in colon cells.

## 1. Introduction

*Clostridioides difficile* (*C. diff*.) is a Gram-Positive, spore-forming bacterium that infects the intestinal tract of humans and animals. In the last decade, *C. diff*. infection has been the leading cause of diarrhea and inflammation of the colon in North America and in Europe^1^. In many cases, *C. difficile* infection is the consequence of a microbial imbalance caused by overtreatment with antibiotics such as penicillin, carbapenem, and fluoroquinolone^2,3^. These disrupt the gut microbiome, allowing the germination of *C. diff*. spores and leading to the proliferation of bacteria and the subsequent release of virulent toxins. In 2017, more than 200K people were infected with *C. diff*. resulting in 12,800 deaths in the United States alone^4,5^. Most of the infections are associated with in-patient care, and more than 80% of the deaths occur in people above 65 years in age^6^. The colonic epithelium is the primary site of infection as the epithelial cells that line the gut wall are highly sensitive to the effects of *C. diff* toxins and *C. diff* preferentially colonizes the colon^7,8^.

The pathogenicity of *C. diff*. derives primarily from two major toxins: Toxin A (TcdA) and Toxin B (TcdB)^9,10^. *C. diff*. adheres to the gut wall using its surface layer proteins and produces two large Rho-glucosylating toxins, TcdA and B, that share ~63% sequence homology^11,12^. These toxins comprise four domains: glucosyltransferase domain (GTD), autoprotease domain (APD), delivery domain and the combined repetitive oligopeptides domain (CROP) (Figure 1A). The *C. diff*. toxins act via a four-step intracellular mechanism: (1) The CROP domain, which is at the C-terminus of the toxins, binds to carbohydrate molecules and proteins on the surface of colonic epithelial cells^13–15^; (2) the delivery domain helps translocate the toxin into the cytosol of the target cells; (3) the APD cleaves the GTD from the rest of the toxin; and (4) the GTD (Figure 1B) utilizes uridine diphosphate glucose (UDP-glucose) to glucosylate Rho-family GTPases that are present in intestinal epithelial cells. The glucosylation of these Rho-family GTPases disrupts transcription, cell cycle progression, apoptosis, and cytoskeleton regulation, leading to cytopathic and cytotoxic effects^16–19^.

**Figure 1.**
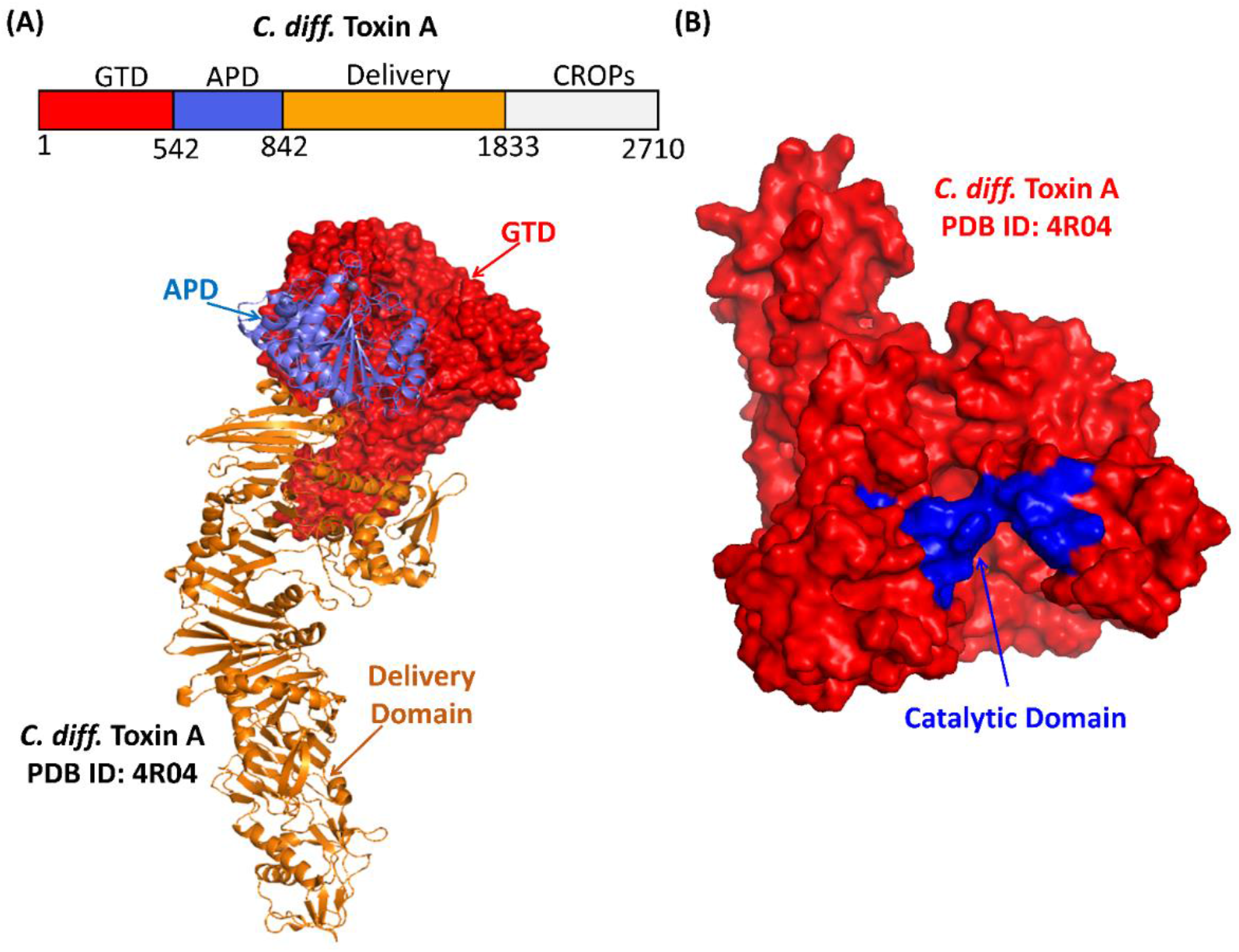
(A) The crystal structure of Toxin A with the glucosyltransferase domain (GTD, red), autoprotease domain (APD, blue) and delivery domain (orange) (PDB ID: 4R04) is illustrated. (B) The catalytic site of TcdA (shown in blue) in the GTD (red) plays an important role in inducing *C. diff*. infection.

Multiple therapeutic approaches have been developed to treat *C. diff*. infection. The standard practice is treatment with antibiotics (metronidazole and vancomycin), but in 20% of cases infection reoccurs^20^. Exposure to these antibiotics alters the microbial community in the gut, facilitating colonization by *C. diff*^21^. Merck introduced a monoclonal antibody, Bezlotoxumab, (marketed as Zinplava) that targets *C. diff* toxin B. While the rate of recurrent infection among patients receiving Bezlotoxumab was substantially lower than for antibiotic-treated cohorts, the high cost of a single dose (~ $4K) and its intravenous infusion are burdensome^22,23^. Another *C. diff*. treatment is Fecal Microbiota Transplant (FMT), an investigational treatment not yet approved by the FDA^24^. The methods of FMT administration and optimal dosing strategies still vary from case to case. Additionally, FMT carries the risk of transmitting infectious diseases and antibiotic-resistant bacteria^25^.

Short peptides are promising candidates for the prevention and treatment of *C. diff*. infection as they are cost effective and can be specific in action. Hence, the goal of this study is to identify peptide inhibitors that bind the catalytic domain of *C. diff*. Toxin A GTD by combining computational design, molecular-level simulations, and experimental refinement. To do this we employ PepBD, a computational *pep*tide *b*inding *d*esign (PepBD) algorithm developed in our group, which performs high-throughput screening of peptide binders to biomolecular targets^26–29^, e.g., proteins and RNA. The PepBD algorithm has been used successfully in the past to design 15-mer transfer RNA^Lys3^-binding peptides^30^, peptides that recognize cardiac troponin I^31^ and neuropeptide Y^32^, peptide ligands that bind to the Fc and Fab domains of immunoglobulin G^33,34^ and peptides that bind to the Receptor Binding Domain of the SARS-CoV-2 spike-protein^35^. In an effort to rank and appraise the computationally suggested peptides, a microfluidic bead-based platform was used to rapidly identify peptides that exhibit the desired binding characteristics. This system uses fluorescence imaging and automated image analysis to measure the propensity for both on-target and off-target binding and has previously been applied to identify peptides that bind specifically to Cas9, VCAM-1, and IgG Fab fragments^36–38^. The efficacy of the peptides is tested via a trans-epithelial electrical resistance (TEER) assay on monolayers of the human gut epithelial culture model and via surface plasmon resonance measurements.

The starting point for the work described in this paper is our previously reported computationally identified 10-mer peptide, “NPA”, that binds to TcdA GTD^39^. This peptide neutralizes TcdA in differentiated small intestinal absorptive cells (SI) but has no effect on differentiated colon absorptive cells. While the mechanisms for this observation are unknown, a possible explanation is that proteases present on the brush border of SI cells cleave the 10-mer peptides into shorter, more-active forms that neutralize the toxins in the SI cells. Since the colon epithelial cells do not appreciably express proteases^40^, the 10-mer peptides are less likely to be cleaved in colon cells than in SI cells, and hence cannot neutralize the toxins in the colon cells.

In this work, we computationally design engineered variants of the NPA peptide shorter than 10 amino acids with the goal of identifying effective inhibitors of *C. diff*. TcdA. We begin by performing molecular dynamics simulations of fragments of NPA to see which of them can bind to the catalytic site of TcdA GTD. The simulation results predict that 8-mer peptide candidates are optimum. Hence, we apply the PepBD algorithm to design 8-mer peptide sequences with 8-mer NPA as the “reference peptide”. Explicit solvent atomistic MD simulations and binding free energy calculations are carried out to evaluate the binding of the *in-silico-suggested* peptides to the TcdA GTD in solution. The peptides are rapidly screened for TcdA binding and TcdA GTD binding through an in-house bead-based peptide display system to eliminate weak peptide inhibitors. The efficacy of the peptides that make it through the bead-based peptide display assay are tested using a trans-epithelial electrical resistance (TEER) assay on monolayers of the human gut epithelial culture model. While conventional cellular toxicity assays for *C. diff* toxins use non-physiologically relevant colon cancer cells or transformed kidney epithelial cells, here we use primary human gut epithelial stem cells from the small intestine (jejunum) and the large intestine (descending colon) that are differentiated in the main lineage (absorptive) of the gut lining. The experimental binding affinity of the top performing peptide to TcdA is characterized using surface plasmon resonance (SPR).

Highlights of our results are as follows. Seven candidate peptide inhibitors (SA1-SA7) were identified using our PepBD algorithm coupled with molecular level simulations. Based on the bead-based peptide display screen for TcdA GTD binding, four peptides (SA1-SA4) were selected for further *in vitro* assessment. SA1 was the only peptide that demonstrated neutralization properties of TcdA in both the small intestine (SI) and colon. The dissociation constant, K_D_, of SA1 to TcdA measured by SPR is 56.1 ± 29.8 nM. These findings suggest that peptide SA1 might be an effective therapeutic drug to treat *C. diff*. infection.

## 2. Results

### 2.1 Determining the Optimal Sequence Length and Initial Peptide Sequence for Designing Peptide Inhibitors of TcdA GTD Catalytic Domain

We began by determining which fragment of NPA plays the most significant role in binding to the TcdA GTD catalytic domain, as this fragment can then serve as the reference peptide in our design process. NPA was identified in our previous study^39^, which reported computationally designed 10-mer peptide sequences (*SI Appendix, Table S1*) that were experimentally tested using a functional cell culture assay for their ability to neutralize TcdA in the small intestinal (jejunum) and large intestinal (colon) cells. The reference peptide used to initialize the computational design in that study was RP: *EGWHAHTGGG* (Figure 2 (A) left), discovered by Feig’s team at Wayne State University^41^ using phage display and verified experimentally by them to inhibit the glucosyltransferase activity of TcdA. Using our PepBD algorithm^26–29^ and molecular dynamics simulations, we identified peptide NPA: *DYWFQRHGHR* (Figure 2 (B) left) that binds to TcdA GTD. Peptides RP and NPA showed toxin-neutralizing activity in jejunum cells but showed no effect in the colon cells. Since cells of the small intestine express proteases on the brush boarder of the cell to break down dietary proteins^40^, we speculated that when NPA was applied to jejunum cells, it was getting cleaved into smaller fragments with neutralizing activity. We performed LC-MS/MS on cell culture supernatants from colonic monolayers after peptide NPA and confirmed the presence of shorter peptides derived from full-length NPA. This is not surprising since expression of common proteases in the colon is nearly absent^42^. Thus, based on the assumption that peptides smaller than 10-mers might act as stronger toxin inhibitors, we modified our computational design approach.

**Figure 2.**
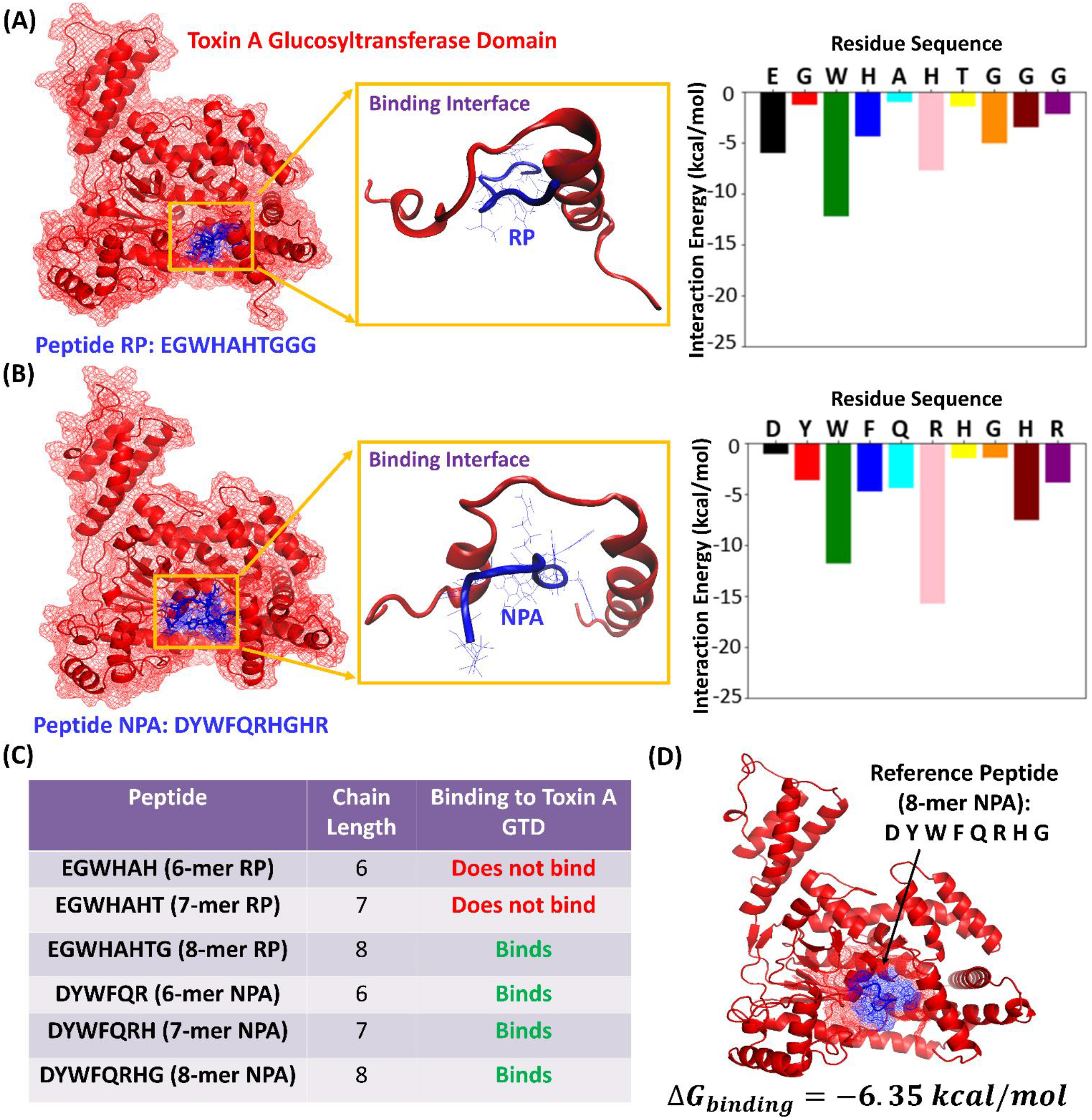
**(A)** Peptide RP (EGWHAHTGGG) and **(B)** NPA (DYWFQRHGHR) at the catalytic site of the Toxin A Glucosyltransferase Domain (left side). The TcdA GTD is shown in red, and the peptide is colored in blue. The residue wise decomposition of the interaction energy plot of peptides RP and NPA is shown on the right. **(C)** Table showing predictions from molecular dynamics simulation of whether or not the 6-mer, 7-mer, and 8-mer fragments on RP and NPA bind to TcdA GTD **(D)** Starting structure for PepBD: Reference peptide NPA (8-mer) bound to the TcdA GTD at the catalytic site with a Δ*G_binding_* of −6.35 kcal/mol.

Based on the experimental observations described above, we proceeded to test via molecular level simulations if peptides shorter than 10-mers can bind to the TcdA GTD. First, we performed atomistic molecular dynamics simulations of RP and NPA bound to the catalytic domain of TcdA GTD. Plots of the interaction energy (van der Waals + electrostatic + polar solvation energy terms) of each of the ten residues along the peptide sequence with the catalytic site of TcdA GTD were generated. The results indicate that residues on the N-terminus of the peptides have a higher contribution to the interaction energy with the catalytic site of TcdA GTD than the flanking residues on the C-terminus of the peptides (Figure 2A and 2B right). Second, we simulated 6-, 7-, and 8-mer fragments of peptides RP and NPA bound to the catalytic domain of TcdA GTD. The simulations reveal that the 6-mer and 7-mer fragments of RP, *EGWHAH* and *EGWHAHT* are not stable (*i.e.,ΔG_binding_* > 0) near the binding site of TcdA GTD, whereas the 8-mer fragment *EGWHAHTG* binds to TcdA GTD with a binding free energy of 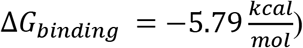. In contrast, all of the fragments of NPA, namely 6-mer *DYWFQR*, 7-mer *DYWFQRH* and 8-mer *DYWFQRHG* 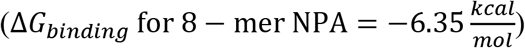 show good binding affinity for the TcdA GTD (Figure 2C). Accordingly, we resolved to design 8-mer peptide variants using *DYWFQRHG* as reference ligand for the PepBD algorithm.

### 2.2 *In-silico* Screening of TcdA GTD Binding Peptides and Evaluation of Binding Free Energies

PepBD is a Monte Carlo-based *pep*tide *b*inding *d*esign algorithm that uses an iterative procedure to optimize the binding affinity and selectivity of peptides to a biomolecular target. The algorithm utilizes the complex formed by a reference peptide sequence and the biomolecular target as input structure and selects peptide variants by implementing sequence and conformation change moves. The desired hydration properties of the designed peptides can be customized based on 6 residue types (hydrophobic, hydrophilic, positive, negative, other, and glycine). The classification of the 20 natural amino acids into the six residue types can be found in *SI Appendix, Table S2*. A score function, Γ*_score_*, which considers (i) the binding energy of the peptide to the receptor and (ii) the conformational stability of the peptide when bound to the receptor, is used to evaluate the acceptance of new peptide candidates. Details of the algorithm are provided in *Methods* and *SI Appendix*.

We implemented the PepBD algorithm to identify an improved set of peptide inhibitors using the 8-mer NPA:TcdA GTD complex (Figure 2D) as the input structure. We investigate three cases with different sets of hydration properties for the peptide chain. For all three cases we ensure diversity in the amino acid composition of the peptide chain by allowing a balance among the various contributions to the binding energy, namely electrostatic, hydrophobic, hydrogen bonding, π-π, etc. which contributes to the peptide’s affinity and selectivity. The three cases are as follows. Case One: *N*_hydrophobic_ = 3, *N*_hydrophilic_ = 2, *N*_positive_ = 2, *N*_negative_ = 1, *N*_other_ = 0 and *N*_glycine_ = 0, Case Two: *N*_hydrophobic_ = 3, *N*_hydrophilic_ = 2, *N*_positive_ = 1, *N*_negative_ = 1, *N*_other_ = 0 and *N*_glycine_ = 1 and Case Three: *N*_hydrophobic_= 1, *N*_negative_ = 3, *N*_positive_ = 2, *N*_negative_ = 1, *N*_other_ = 0 and *N*_glycine_ = 1. For each case we perform the PepBD search with three different initial random seed numbers to randomize the initial peptide sequence. This enables our designs to proceed along different search pathways and sample peptides from a large pool of peptide sequences and conformations. During the *in-silico* evolution, new sequences and conformers are generated by mutating and exchanging amino acids on the peptide chain, which results in a fluctuation of the score. A lower Γ_score_ means stronger binding affinity of a peptide to the bound target. The root-mean-squared deviation, RMSD, of the new peptide conformers compared to the conformation of the initial peptide chain’s conformation reflects the changes in the backbone scaffold of the peptide as the design process progresses. Figure 3A shows the Score (Γ_score_) and the RMSD profile vs the number of sequence and conformation change moves performed with a distinct initial random seed for Case 1. Figure 3B shows the structure of one of the top performing peptides, SA1: *EFWWRRHN*, complexed with the TcdA GTD binding interface from Case 1. Peptide SA1 has a Γ_score_ = −44.59 which was obtained at the 459^th^ step of the sequence evolution. From Case 2 the top performing peptide is SA3 *QEWMGRHW* (*SI Appendix, Fig S1A and S1C*), and from Case 3 the top performing peptide is SA6 *EGWQHRHR* (*SI Appendix, Fig S1B and S1D*); more information on SA3 and SA6, and Cases 2 and 3 is provided in *SI Appendix*.

**Figure 3.**
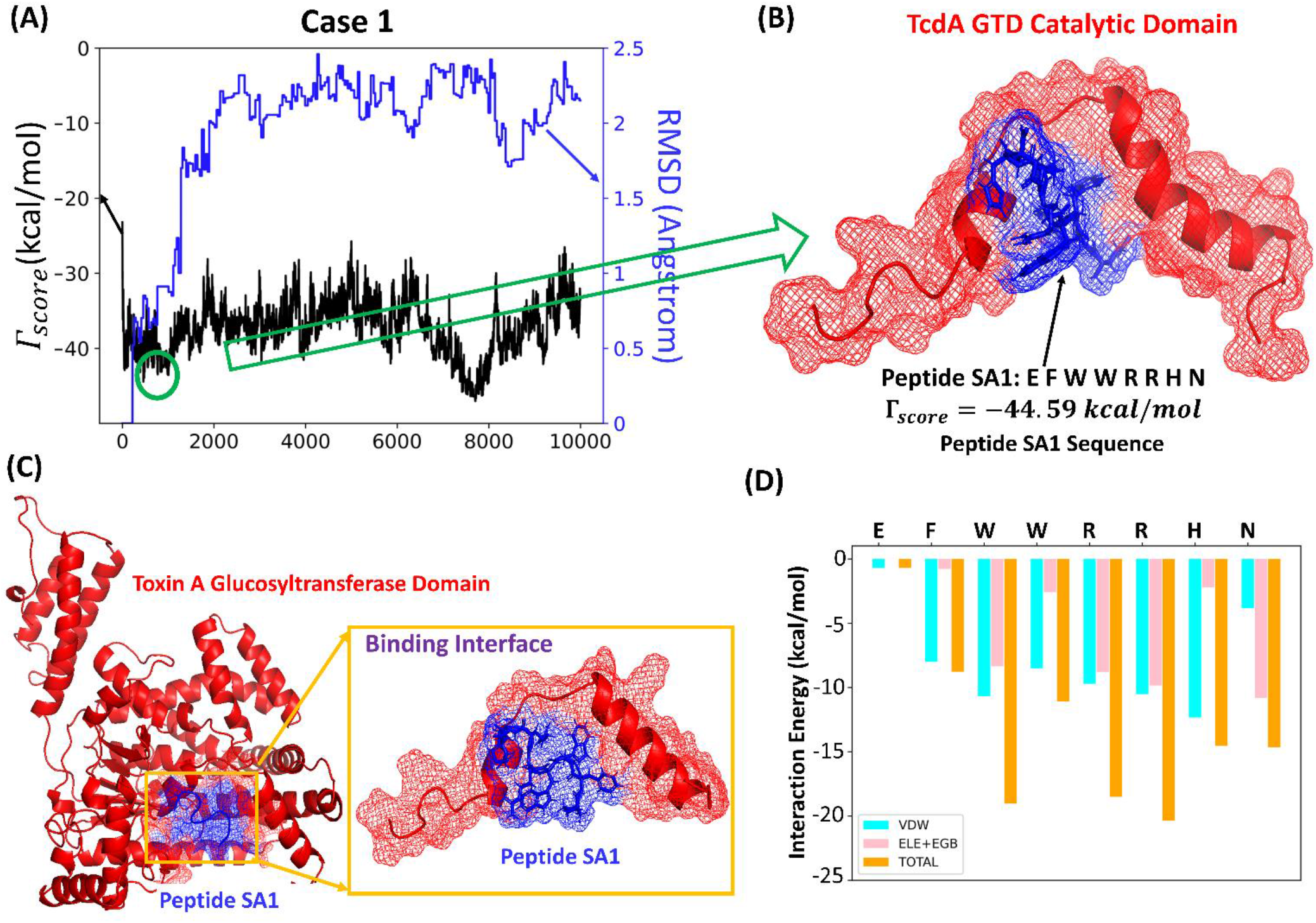
**(A)** The score/RMSD vs the number of sequence and conformation steps for Case 1 with a distinct initial random seed results in **(B)** peptide SA1: EFWWRRHN **(C)** Snapshot of peptide SA1 bound to TcdA GTD obtained from molecular dynamics simulation. The conformation of the peptide at the binding interface is shown. **(D)** Plot showing the residue-wise decomposition of the interaction energy (van der Waals + Electrostatic + Polar solvation energy contribution) in the SA1:TcdA GTD complex.

Once the *in-silico* evolution terminates, we perform explicit-solvent atomistic molecular dynamics simulations of the complexes formed between the lowest scoring peptides and TcdA GTD to predict their binding affinity. Three independent simulations are carried out for each peptide:TcdA GTD complex for 100 ns to ensure that the system reaches an equilibrated state. The simulations are performed at 298K using the AMBER ff14SB forcefield and the AMBER18 package. We calculate the Δ*G_binding_* of the peptide:receptor complex following the MD simulations using the MMGBSA protocol and variable dielectric constant method. Details of our atomistic MD simulation and Δ*G_binding_* calculation procedure are provided in *Methods* and *SI Appendix*^26–29,43^. Table 1 reports the top peptides that we obtain from our computational procedure and their corresponding scores and Δ*G_binding_* values. SA1 is the most promising peptide with a Δ*G_binding_* value of −15.94 kcal/mol. (Note: the lower the value of Δ*G_binding_*, the higher the binding affinity). The SA1:TcdA GTD complex obtained by performing a hierarchical clustering analysis on the last 5 ns of a 100 ns MD simulation is shown in Figure 3C. Additionally, the plot of the residue-wise decomposition of the interaction energy between SA1 and the catalytic site of TcdA GTD is shown in Figure 3D. The plot reveals that the critical SA1 residues involved in TcdA GTD binding are Trp3, Trp4, Arg5, Arg6, His7 and Asn8. Thus, tryptophan, arginine, histidine, and asparagine on SA1 are the four essential amino acids for TcdA GTD binding. A detailed discussion of key amino acid interactions of SA1 with TcdA GTD is provided in the *SI Appendix*. Amino acid sequence signatures in *C. diff* TcdA GTD binding peptides were derived from the residue composition of the top 1% of the lowest scoring peptides identified by PepBD for Cases 1 and 2 and are reported in *SI Appendix, Table S3*.

**Table 1.**
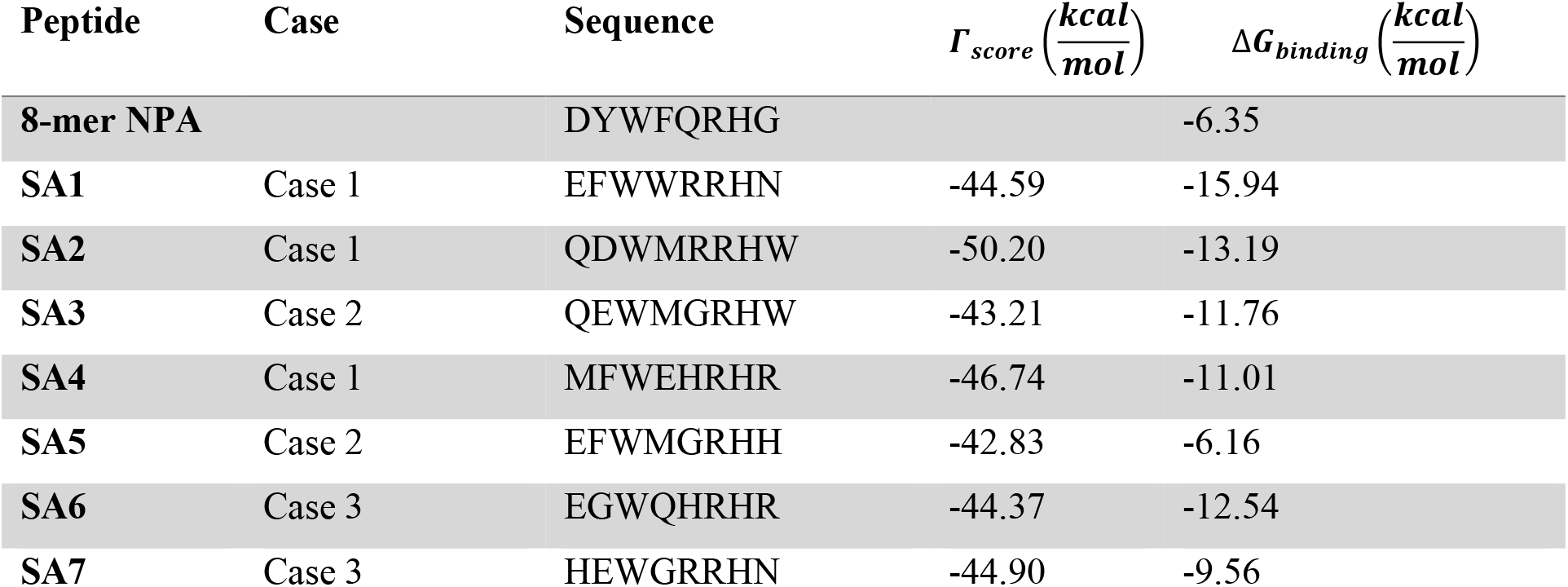
The initial peptide sequence (8-mer NPA) and the list of peptide sequences identified by PepBD screening with their corresponding ΔΓ*_score_* and Δ*G_binding_* values.

### 2.3 Bead-Based Pre-Screening for TcdA Binding

The candidate peptide inhibitors suggested by PepBD were pre-screened *in vitro* for potent and selective TcdA GTD binding to eliminate weak inhibitors prior to the cell-based assays. Robust inhibition of TcdA glucosyltransferase activity relies on the ability of the peptides to outcompete the TcdA GTD’s substrate, UDP-Glucose. Given TcdA’s relatively low Michaelis constant (K_M_ ~ 4.5 μM) compared to the cellular concentration of its substrate UDP-Glucose (92 μM), it is especially important that the peptides exhibit high binding strength and selectivity to TcdA GTD (see *SI Appendix* for detailed analysis)^11,44–47^.

Using a microfluidic screening system developed by our team in prior work^36,38^, we implemented a dual-fluorescence assay to evaluate the TcdA GTD inhibitory activity of the peptide candidates SA1-SA7, 8-mer NPA and 8-mer RP. Each peptide sequence was immobilized on a translucent ChemMatrix Aminomethyl bead that was then contacted with red fluorescently labeled TcdA (TcdA-AF594) and with a green-fluorescent analog of UDP-Glucose (UDP-Glucose-Fluorescein). Beads displaying peptides that bind to TcdA exhibit a red fluorescence signal; beads exhibit green fluorescence when UDP-Glucose-Fluorescein binds to the catalytic site of the TcdA GTD. Thus, peptides that can selectively bind to the TcdA GTD and displace the UDP-Glucose-Fluorescein from the catalytic site will be exhibit red, but not green, fluorescence. Beads are analyzed by a high-throughput assay (350 beads per hr) that correlates the fluorescence intensity of the beads to the binding strength and selectivity of the peptides displayed. Images of each bead are analyzed via a custom algorithm that ensures consistent and objective bead characterization within a screened ensemble.

Beads displaying TcdA-binding peptides accumulate TcdA-AF594 on their surface, leading to a characteristic red halo fluorescence. This is because TcdA is quite large (MW=308 kDa, 2710 residues) and hence, poorly diffuses into the narrow pores of the beads. The intensity of the halo correlates with the amount of TcdA-AF594 bound^38^ (Figure 4B). Beads displaying selective TcdA GTD-binding peptides displace UDP-Glucose-Fluorescein from the GTD, and thus peptide:TcdA GTD binding, can be observed as a loss of green fluorescence^44^ from the TcdA binding region (Figure 4C).

**Figure 4.**
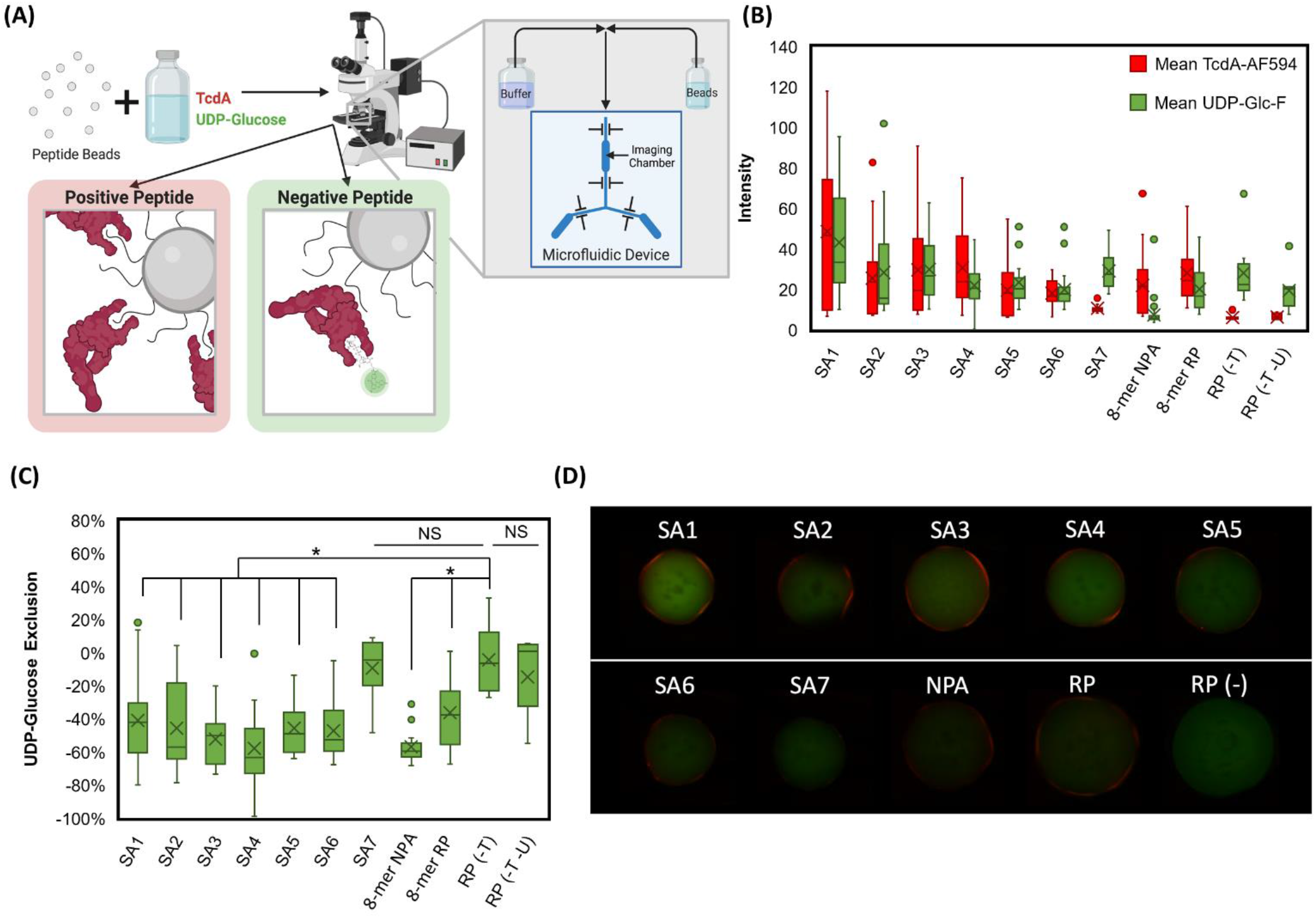
Bead based screening for selective TcdA GTD binding for each peptide sequence using TcdA-AF594 (in red) and UDP-Glucose-Fluorescein (in green). **(A)** Schematic for bead capture and imaging on custom microfluidic device and microscope. **(B)** Mean red and green intensity (0-255 pixel value) of 90^th^ percentile red area (TcdA halo) of beads displaying each peptide (number of beads (n) = 23, 20, 25, 19, 23, 21, 15, 36, 14, 8, 7 respectively). RP (-T) and RP (-T -U) are no TcdA-AF594 and no TcdA-AF594/UDP-Glucose-Fluorescein controls, respectively. **(C)** Percent change in green fluorescence from 90^th^ percentile red halo region to 50^th^ percentile green; more negative values indicate exclusion of UDP-Glucose from peptide:TcdA interface, and thus selective peptide binding to the TcdA GTD (* p<0.005). **(D)** Representative red and green composite images for each peptide.

The performance of the peptides (SA1-SA7, 8-mer NPA and 8-mer RP) was inferred based on the intensity of the red halo fluorescence, and on the loss of green fluorescence of the beads, respectively (Figure 4D). SA1 was the most promising TcdA binding peptide with the highest mean red halo fluorescence. Despite having high green fluorescence, there was a significant (p = 0.0011) reduction in the green fluorescence from the center of the beads to the peptide:TcdA binding interface compared to the no-TcdA-AF594 no-UDP-Glucose-Fluorescein control (RP(-T -U)) (Figure 4C), meaning that SA1 bound selectively to TcdA GTD. SA1, SA2, SA3 and SA4 had higher mean red halo fluorescence, and thus higher TcdA binding, than 8-mer NPA. SA1, SA3 and SA4 had higher mean red halo fluorescence, and thus higher TcdA binding, than 8-mer RP. Notably, all of the peptides that were screened showed binding to TcdA, however the peptides from Cases 1 (SA1, SA2, SA4) and 2 (SA3, SA5) performed significantly better than those from Case 3 (SA6, SA7). All peptides except SA7 showed a significant (p < 0.005) decrease in green fluorescence from the center to the halo (TcdA binding) region, indicating relatively universal exclusion of UDP-Glucose from the peptide:TcdA interface (Figure 4C). Despite differences in background green fluorescence between peptides, the uniform decrease in green fluorescence at the TcdA halo provided little information to differentiate between peptides. The selective binding of most PepBD designed peptides to the TcdA GTD demonstrates the robustness of the peptide design algorithm to reliably identify peptide binders to particular pockets. Peptides SA1-SA4, which had higher red fluorescence than NPA and thus promising TcdA binding, were selected for further evaluation.

### 2.4 Functional testing of SA1 on human colonic epithelium

Peptides SA1-SA4 were screened on a human colon epithelial culture system to preliminarily evaluate neutralizing capabilities. In this assay system, colonic epithelial stem cells are applied to transwell inserts and cultured to confluence on the permeable membrane of the insert. Once the cell barrier is achieved after about 4 days of ISC (intestinal stem cell) expansion, the media is changed to promote differentiation into the primary absorptive lineage of the colon and the cell type most exposed to C. difficile toxins^48–50^. Tight junction proteins are upregulated thereby increasing the barrier function of the cellular monolayer and decreasing the ion flux, which is measured as Trans-Epithelial Electrical Resistance (TEER)^51^.

Toxicity is measured over time as a drop in TEER indicating TcdA-dependent changes in cytoskeleton, which cause leaky tight junctions. For the quick screen, peptides were pre-incubated with differentiated colon epithelial monolayers for 2 hours, then TcdA was added at a concentration of 30 pM, which is the concentration observed in stool of *C. diff* patients^51^. SA1 demonstrated some neutralizing capabilities in the primary screen as observed by preservation of TEER compared to conditions with TcdA without SA1 (data not shown). Because SA1 showed promising neutralizing activity, it was more extensively tested for efficacy using the same TcdA toxicity assay. As expected, colon monolayers responded with a near complete loss of TEER at 12 hours post-TcdA exposure, however, pre-exposure of monolayers with SA1 produced ~79% protection from TcdA toxicity (Figure 5). These data demonstrate the ability of SA1 to protect barrier function in the presence of clinically relevant concentrations of TcdA.

**Figure 5.**
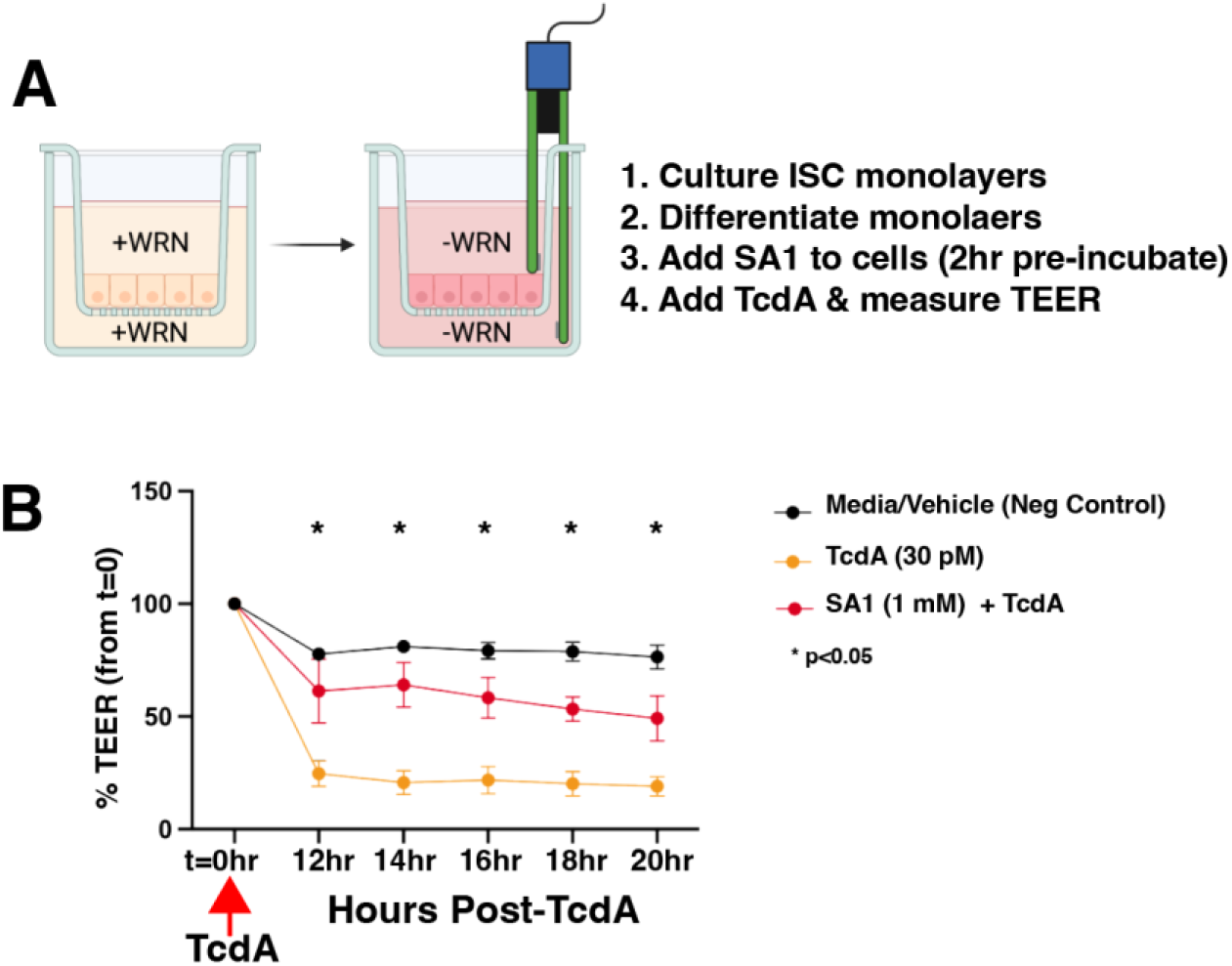
Peptide SA1 has functional neutralizing effects on TcdA in a human colonic epithelial culture model. (A) Summary of experimental design and schematic of human ISC expansion in defined media (EM) of human colonic epithelial monolayers on transwells (*left*). Forced differentiation with defined media (DM) (*right*). **(B)** TEER measurements on differentiated Colon (Descending) monolayers treated with DM (media with vehicle), TcdA, and TcdA with SA1. Values are expressed as a % of TEER at t=0 when TcdA was added to cultures. One way ANOVA p<0.005, multiple comparisons tests 12hr-20hr p<0.05.

### 2.5 Measuring SA1 Kinetic Parameters by Surface Plasmon Resonance

Surface plasmon resonance was used to measure key kinetic parameters for SA1:TcdA binding. SA1 was covalently bound to a mixed bio-resistant thiol self-assembled monolayer (SAM) on the gold sensor surface. Characterization of the surface thickness and composition can be found in *SI Appendix*.

Various concentrations of the analyte, TcdA, were injected over the surface and the net angular response in degrees was recorded. The amount of analyte, TcdA, bound to the surface was calculated using a device-specific conversion factor previously described^52^. The net equilibrium response was fit to a Langmuir isotherm (Equation 1) as shown in Figure 6A, resulting in an equilibrium dissociation constant, K_D_, of 56.1 ± 29.8 nM and a maximum binding capacity, Q_max_, of 12.0 ± 2.2 nmol/m^2^.

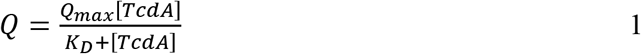

SA1:TcdA binding is a moderate-to-high affinity interaction based on its mid-nanomolar K_D_. A fuller picture of the binding emerges by considering the kinetics in addition to the equilibrium state. Rapid recognition, which is characterized by a high adsorption rate constant (k_a_), is important in the context of competitive inhibitors. Additionally, complex stability, as reflected by a low desorption rate constant (k_d_), is desired. To succeed as a potent competitive TcdA inhibitor, SA1 needs to outcompete UDP-Glucose for the active site and slowly dissociate from it. Dynamic response measurements over TcdA injections allowed for the calculation of k_a_ and k_d_ using Equation 2, which are summarized in Table 2, and shown in Figure 6B.

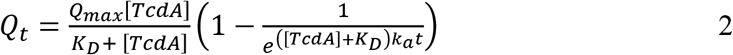

The average k_a_ was found to be 7.0 ± 2.5 x10^-5^ nmol^-1^ s^-1^, which is high for a peptide inhibitor, given its relative backbone flexibility. Review of the literature indicates that peptide inhibitor adsorption rate constants typically fall between 10^-7^ and 10^-6^ nmol^-1^ s^-1^ ^53,54^ The average k_d_ was found to be 4.0 ± 1.4 x10^-3^ s^-1^, which is consistent with that of other peptide ligands^53–57^. Compared to antibody:antigen binding interactions, which are known to be high affinity (typical K_D_ are between 10^-8^ and 10^-11^ M), SA1:TcdA (K_D_ ~ 10^-8^ M) exhibits remarkably high binding affinity for non-antibody:antigen biorecognition^58^.

**Figure 6.**
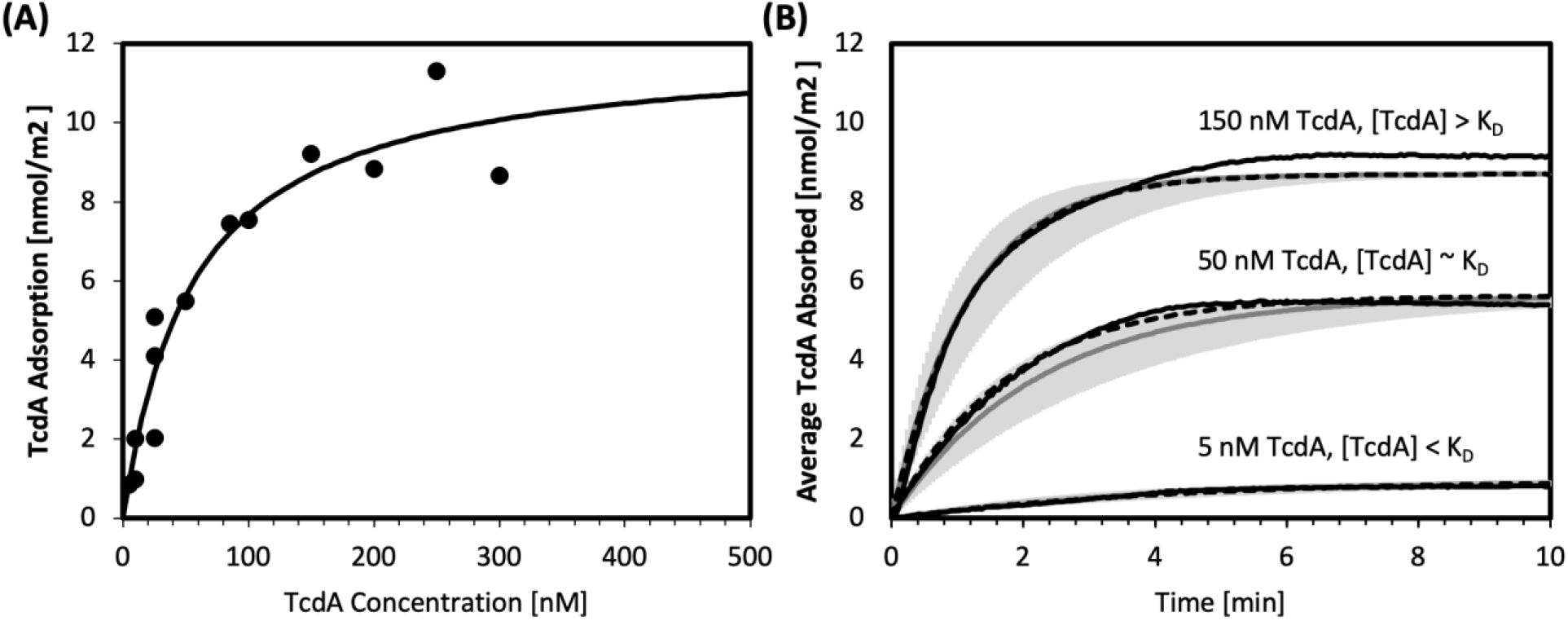
Surface plasmon resonance adsorption isotherms and dynamic responses for SA1:TcdA binding. (A) Maximum TcdA adsorption and TcdA concentration fit to a Langmuir isotherm (Equation 1) yielding K_D_ and Q_max_. (B) Experimental dynamic response (black solid lines), individual theoretical fit (black dashed lines), and average theoretical fit (solid grey lines, 95% confidence interval in light grey) for TcdA concentrations above, below, and approximately at K_D_.

**Table 2.**
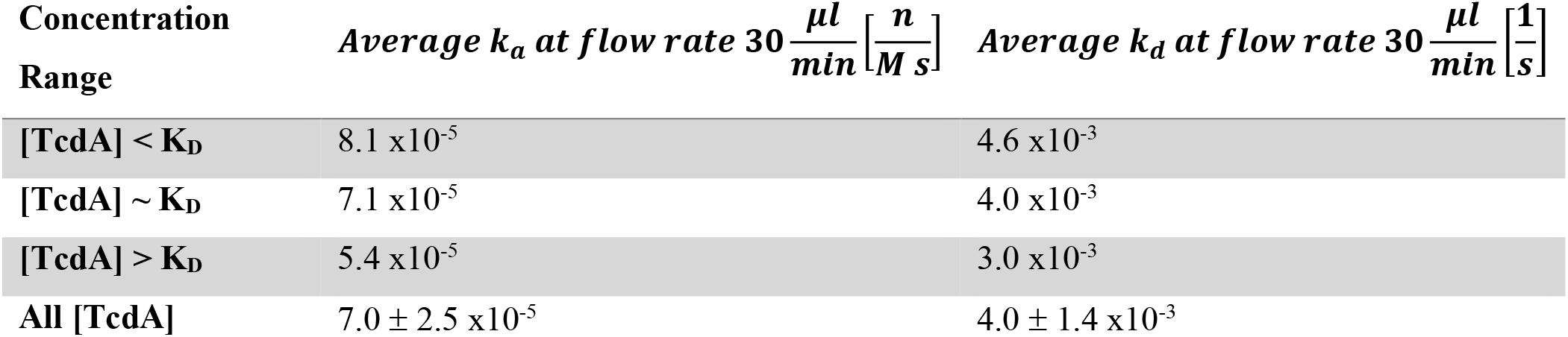
Association and dissociation constants for various TcdA concentration ranges.

| Concentration Range | Average $k_a$ at flow rate $30 \frac{\mu l}{min} \left[ \frac{n}{M s} \right]$ | Average $k_d$ at flow rate $30 \frac{\mu l}{min} \left[ \frac{1}{s} \right]$ |
| --- | --- | --- |
| [TcdA] < $K_D$ | $8.1 \times 10^{-5}$ | $4.6 \times 10^{-3}$ |
| [TcdA] ~ $K_D$ | $7.1 \times 10^{-5}$ | $4.0 \times 10^{-3}$ |
| [TcdA] > $K_D$ | $5.4 \times 10^{-5}$ | $3.0 \times 10^{-3}$ |
| All [TcdA] | $7.0 \pm 2.5 \times 10^{-5}$ | $4.0 \pm 1.4 \times 10^{-3}$ |

## 3. Materials and Methods

The materials and methods are described in detail in *SI Appendix*. The *Peptide B*inding *D*esign (PepBD) algorithm was used to design peptide inhibitors that target the TcdA GTD. Explicit-solvent atomistic molecular dynamics simulation using the AMBER 18 package was employed to investigate the dynamics of the binding process between the peptide sequences and TcdA GTD. The PepBD algorithm and the molecular dynamics simulation procedure are explained in detail in *SI Appendix*. The *in-silico* identified peptide sequences were screened for binding to TcdA on an in-house microfluidic bead imaging system originally designed for sorting solid phase peptide libraries. Exclusion of a fluorescent UDP-Glucose co-factor analog (UDP-Glucose-Fluorescein) from the UDP-Glucose binding pocket of TcdA was used as a proxy for visualizing peptide binding in the desired position. The bead-based screening assay is described in detail in *SI Appendix*. Peptides SA1-SA4 for functional testing on human gut epithelium were synthesized on Wang resin pre-loaded with the first amino acid on an Initiator+ Alstra (Biotage, Uppsala, Sweden) following the Fmoc/tBu protecting strategy. Details are provided in *SI Appendix* A new screening assay was used to evaluate the TcdA/B toxicities in the presence of peptides. Briefly, human colonic epithelial stem cells were isolated from healthy adult donor organs. The stem cells were expanded as described and applied to transwell inserts, expanded in defined expansion media (EM) to create a barrier, and differentiated to mature absorptive colonocytes in defined differentiated cell media (DM)^49,50^. Candidate peptides were preincubated with colonic monolayers and then TcdA was added to the apical reservoir of the transwell. TEER was measured over time to evaluate barrier function. A Drop in TEER was considered to reflect the extent of toxicity due to impaired tight junctions caused by the toxins. Peptide efficacy was measured as reduced loss of TEER compared to TcdA alone. Details are provided in *SI Appendix*. The equilibrium dissociation constant, K_D_, of SA1:TcdA binding was measured using Surface Plasmon Resonance (SPR). Details are provided in *SI Appendix*.

## 4. Conclusion

The goal of this work was to identify lead peptide candidates that bind to the catalytic site of TcdA glucosyltransferase domain and hence can inhibit the glucosyltransferase activity of toxin TcdA. In our previous work, we reported 10-mer peptides that were able to neutralize TcdA in differentiated small intestinal absorptive cells (SI) but showed no effect on differentiated colon absorptive cells. A likely explanation for this observation is that proteases present on the brush border of SI cells cleaved the 10-mer peptides into shorter, more-active forms that neutralize the toxins in the SI cells, but such an effect is not possible in the colon cells as they do not express proteases significantly. We performed atomistic molecular dynamics simulations to test if shorter peptides might be more effective inhibitors of *C. diff*. TcdA than the 10-mers. The simulations predicted that 8-mer peptides are at an optimum peptide length. We applied our PepBD algorithm and combined it with molecular dynamics simulations and binding free energy calculations to find seven 8-mer peptides (SA1-SA7). These peptides were rapidly screened using an in-house bead-based microfluidic screening technique to check for selective TcdA-GTD binding, and the weak peptide inhibitors (SA5-SA7) were eliminated. The efficacies of peptides SA1-SA4 were tested using a trans-epithelial electrical resistance (TEER) assay on monolayers of the human gut epithelial stem cells from the small intestine (jejunum) and the large intestine (descending colon). Peptide SA1 blocked TcdA toxicity in both jejunum and colon epithelial cells. The binding affinity of this peptide to TcdA was characterized using surface plasmon resonance (SPR) and the dissociation constant, K_D_, was found to be 56.1 ± 29.8 nM.

Anti-toxin drugs (such as SA1 developed here) are therapeutically relevant because they eliminate the causative agent of disease (e.g., the toxin). Anti-toxin drugs are also appealing because they are highly specific (limiting off-target effects on the host or microbiota) and impose low, if any, fitness cost on the toxin-producing pathogen (thereby reducing the pressure for resistance to develop). In this way, anti-toxin drugs are complementary to (and potentially synergistic with) standard-of-care antibiotics. By binding to the toxin’s catalytic site, SA1 competitively inhibits the key disease-causing biochemical reaction employed by *C. diff*. Further, because the toxin active site is the most highly conserved region of the toxin, we hope that SA1 will be active on a variety of *C. diff* strains and maintain robust activity as new strains arise. Looking forward, the ease with which peptides can be manufactured at scale promises to improve the equitability of access to anti-toxin therapies, and also allows them to be manufactured at the site of disease via engineered gut microbes. Taken together, this work illustrates a structure-guided, rational approach to designing anti-toxin peptides that is readily generalizable to other toxins secreted by *C. diff* and other antibiotic-resistant pathogens.

## Acknowledgements

This research was supported by funds from the National Science Foundation (CBET-1934284), from the National Institutes of Health (R01DK109559), from the North Carolina Biotechnology Center (2019-FLG-3841) and from the UNC Center for Gastrointestinal Biology and Disease, P30DK034987. This work used the Extreme Science and Engineering Discovery Environment (XSEDE), supported by National Science Foundation (ACI-1548562). S. Sarma, X. Xiao and C.K. Hall thank the San Diego Supercomputer Center (SDSC) for computing time.

## References

1. E. G. McDonald, T.C. Lee, Clostridium difficile Infection. N. Engl. J. Med. 373, 286–288 (2015).

2. J. Czepiel, et al., Clostridium difficile infection: review. Eur. J. of Clin. Microbiol. Infect. Dis. 38, 1211–1221 (2019).

3. Centers for Disease Control and Prevention, Nearly half a million Americans suffered from Clostridium difficile infections in a single year. Page last reviewed: March 22, 2017 (archived document); Available from: https://www.cdc.gov/media/releases/2015/p0225-clostridium-difficile.html (accessed 2 November 2022).

4. Balsells E, Shi T, Leese C, et al. Global burden of Clostridium difficile infections: A systematic review and meta-analysis. J. Glob. Health. 9, 010407 (2019).

5. Centers for Disease Control and Prevention, Clostridioides difficile Infection. Page last reviewed: November 13, 2019; Available from: https://www.cdc.gov/hai/organisms/cdiff/cdiff_infect.html (accessed 2 November 2022).

6. F. C. Lessa, et al., Burden of Clostridium difficile Infection in the United States. N. Engl. J. Med. 372, 825–834 (2015).

7. L. M. Napolitano, C. E. Edminstion, Clostridium difficile disease: Diagnosis, pathogenesis, and treatment update. Surgery. 162, 325–348 (2017).

8. M. Rupnik, M. H. Wilcox, D. N. Gerding, Clostridium difficile infection: new developments in epidemiology and pathogenesis. Nat. Rev. Microbiol. 7, 526–536 (2009).

9. K. Aktories, C. Schwan, T. Jank, Clostridium difficile Toxin Biology. Annu. Rev. Microbiol. 71, 281–307 (2017).

10. M. C. Abt, P. T. McKenney, E. G. Pamer, Clostridium difficile colitis: Pathogenesis and host defence. Nat. Rev. Microbiol. 14, 609–620 (2016).

11. W. J. Bradshaw, A. K. Roberts, C. C. Shone, K. R. Acharya, The structure of the S-layer of Clostridium difficile. J. Cell. Commun. Signal. 12, 319–331 (2018).

12. R. P. Fagan, et al., Structural insights into the molecular organization of the S-layer from Clostridium difficile. Mol. Microbiol. 71, 1308–1322 (2009).

13. J. G. S. Ho, A. Greco, M. Rupnik, K. K. S. Ng, Crystal structure of receptor-binding C-terminal repeats from Clostridium difficile toxin A. Proc. Natl. Acad. Sci. U.S.A. 102, 18373–18378 (2005).

14. T. Murase, et al., Structural basis for antibody recognition in the receptorbinding domains of toxins a and B from clostridium difficile. J. of Biol. Chem. 289, 2331–2343 (2014).

15. L. Tao, et al., Sulfated glycosaminoglycans and low-density lipoprotein receptor contribute to *Clostridium difficile* toxin A entry into cells. Nat. Microbiol. 4, 1760–1769 (2019).

16. M. O. P. Ziegler, T. Jank, K. Aktories, G. E. Schulz, Conformational Changes and Reaction of Clostridial Glycosylating Toxins. J. Mol. Biol. 377, 1346–1356 (2008).

17. P. Chen, et al., Structure of the full-length Clostridium difficile toxin B. Nat. Struct. Mol. Biol. 26, 712–719 (2019).

18. D. Lyras, et al., Toxin B is essential for virulence of Clostridium difficile. Nature. 458, 1176–1179 (2009).

19. S. A. Kuehne, et al., The role of toxin A and toxin B in Clostridium difficile infection. Nature. 467, 711–713 (2010).

20. C. Slimings, T. V. Riley, Antibiotics and hospital-acquired Clostridium difficile infection: update of systematic review and meta-analysis. J. of Antimicrob. Chemother. 69, 881–891 (2014).

21. B. B. Lewis, et al., Loss of Microbiota-Mediated Colonization Resistance to Clostridium difficile Infection with Oral Vancomycin Compared with Metronidazole. J. Infect. Dis. 212, 1656–1665 (2015).

22. Y. Lee, et al., Bezlotoxumab (Zinplava) for Clostridium Difficile Infection: The First Monoclonal Antibody Approved to Prevent the Recurrence of a Bacterial Infection. P. T. 42, 735–738 (2017).

23. K. Kinoshita, Preclinical and clinical properties of bezlotoxumab (ZINPLAVA® 25 mg/ml concentrate for solution for infusion), novel therapeutic agent for clostridium difficile infection. Folia Pharmacol. Jpn. 152, 39–50 (2018).

24. B. J. Kelly, P. Tebas, Clinical Practice and Infrastructure Review of Fecal Microbiota Transplantation for Clostridium difficile Infection. Chest. 153, 266–277 (2018).

25. Z. DeFillip, et al., Drug-Resistant *E. coli* Bacteremia Transmitted by Fecal Microbiota Transplant. N. Engl. J. Med. 381, 2043–2050 (2019)

26. X. Xiao, C. K. Hall, P. F. Agris, The design of a peptide sequence to inhibit HIV replication: A search algorithm combining Monte Carlo and self-consistent mean field techniques. J. Biomol. Struct. and Dyn. 32, 1523–1536 (2014).

27. X. Xiao, M. E. Hung, J. N. Leonard, C. K. Hall, Adding energy minimization strategy to peptide-design algorithm enables better search for RNA-binding peptides: Redesigned λ N peptide binds boxB RNA. J. Comp. Chem. 37, 2423–2435 (2016).

28. X. Xiao, P. F. Agris, C. K. Hall. Introducing folding stability into the score function for computational design of RNA-binding peptides boosts the probability of success. Proteins: Struct. Funct. Genet. 84, 700–711 (2016).

29. X. Xiao, Y. Wang, J. N. Leonard, C. K. Hall. Extended Concerted Rotation Technique Enhances the Sampling Efficiency of the Computational Peptide-Design Algorithm. J. Chem. Theory Comput. 13, 5709–5720 (2017).

30. J. L. Spears, X. Xiao, C. K. Hall, P. F. Agris. Amino acid signature enables proteins to recognize modified tRNA. Biochemistry. 53, 1125–1133 (2014).

31. X. Xiao, et al., Advancing Peptide-Based Biorecognition Elements for Biosensors Using in-Silico Evolution. ACS Sensors. 3, 1024–1031 (2018).

32. X. Xiao, et al., In Silico Discovery and Validation of Neuropeptide-Y-Binding Peptides for Sensors. J. Phys. Chem. B. 124, 61–68 (2020).

33. X. Xiao, et al., Novel peptide ligands for antibody purification provide superior clearance of host cell protein impurities. J. Chromatogr. A. 1625, 461237 (2020).

34. X. Xiao, et al. De novo discovery of peptide-based affinity ligands for the fab fragment of human immunoglobulin G. J. Chromatogr. A. 1669, 462941 (2022).

35. S. Sarma, S. M. Herrera, X. Xiao, G. A. Hudalla, C. K. Hall, Computational Design and Experimental Validation of ACE2-Derived Peptides as SARS-CoV-2 Receptor Binding Domain Inhibitors. J. Phys. Chem. B. 126, 8129–8139 (2022)

36. K. Day, et al., Discovery and Evaluation of Peptide Ligands for Selective Adsorption and Release of Cas9 Nuclease on Solid Substrates. Bioconjugate Chem. 30, 3057–3068 (2019).

37. R. Prodromou, et al., Engineering Next Generation Cyclized Peptide Ligands for Light-Controlled Capture and Release of Therapeutic Proteins. Adv. Funct. Mater. 31, 2101410 (2021).

38. S Saberi-Bosari, et al., Affordable Microfluidic Bead-Sorting Platform for Automated Selection of Porous Particles Functionalized with Bioactive Compounds. Scientific Reports. 9, 7210 (2019).

39. X. Xiao, et al. In Silico Identification and Experimental Validation of Peptide-Based Inhibitors Targeting Clostridium difficile Toxin A. ACS Chem. Biol. 17, 118–128 (2022).

40. N. C. Chumbler, et al., Crystal structure of Clostridium diffiicile toxin A. Nat. Microbiol. 15002, 1–6 (2016).

41. S. J. Abdeen, R. J. Swett, A. L. Feig, Peptide inhibitors targeting clostridium difficile toxins A and B. ACS Chem. Biol. 5, 1097–1103 (2010)

42. J. Burclaff, et al., A Proximal-to-Distal Survey of Healthy Adult Human Small Intestine and Colon Epithelium by Single-Cell Transcriptomics. Cell Mol. Gastroenterol. Hepatol. 13, 1544–1589 (2022).

43. X. Xiao, B. Zhao, P. F. Agris, C. K. Hall, Simulation study of the ability of a computationally-designed peptide to recognize target tRNALys3 and other decoy tRNAs. Protein Sci. 25, 2243–2255 (2016).

44. J. W Loughney, et al., Development of a non-radiolabeled glucosyltransferase activity assay for C. difficile toxin A and B using ultra performance liquid chromatography. J. Chromatogr. A. 1498, 169–175 (2017).

45. W. P. Ciesla, D. A. Bobak. Clostridium difficile toxins A and B are cation-dependent UDP-glucose hydrolases with differing catalytic activities. J. Biol. Chem. 273, 16021–16026 (1998).

46. M. R. Laughlin, W. A. Petit, J. M. Dizon, R. G. Shulman, E. J. Barrett. NMR measurements of in vivo myocardial glycogen metabolism. J. Biol. Chem. 263, 2285–2291 (1988).

47. S. Bhattacharyya, A. Kerzmann, A. L. Feig, Fluorescent analogs of UDP-glucose and their use in characterizing substrate binding by toxin A from Clostridium difficile. Eur. J. Biochem. 269, 3425–3432 (2002).

48. D. B. Gunasekara, et al., A Monolayer of Primary Colonic Epithelium Generated on a Scaffold with a Gradient of Stiffness for Drug Transport Studies. Anal. Chem. 90, 13331–13340 (2018).

49. Y. Wang, et al., Self-renewing Monolayer of Primary Colonic or Rectal Epithelial Cells. Cell. Mol. Gastroenterol. Hepatol. 4, 165–182 (2017).

50. I. Gomez-Martinez, et al., A Planar Culture Model of Human Absorptive Enterocytes Reveals Metformin Increases Fatty Acid Oxidation and Export. Cell. Mol. Gastroenterol. Hepatol. 14, 409–434 (2022).

51. L. Song, et al., Development and Validation of Digital Enzyme-Linked Immunosorbent Assays for Ultrasensitive Detection and Quantification of Clostridium difficile Toxins in Stool. J. Clin. Microbiol. 53, 3204–3212 (2015).

52. N. Islam, F. Shen, P. V. Gurgel, O. J. Rojas, R. G. Carbonell, Dynamic and equilibrium performance of sensors based on short peptide ligands for affinity adsorption of human IgG using surface plasmon resonance. Biosens. Bioelectron. 58, 380–387 (2014).

53. Y. Wang, et al., PD-1-targeted discovery of peptide inhibitors by virtual screening, molecular dynamics simulation, and surface plasmon resonance. Molecules. 24, 3784 (2019).

54. P. V. Ershov, et al., Kinetic and thermodynamic analysis of dimerization inhibitors binding to HIV protease monomers by surface plasmon resonance. Biochem. (Mosc.) Suppl. B: Biomed. Chem. 6, 94–97 (2012).

55. M. C. Weiger, et al., Quantification of the binding affinity of a specific hydroxyapatite binding peptide. Biomaterials. 31, 2955–2963 (2010).

56. E. Weiss, et al., Determination of Kinetic Constants for the Interaction between a Monoclonal Antibody and Peptides Using Surface Plasmon Resonance. Biochemistry. 31, 6298–6304 (1992).

57. M. M. Morelock, R. H. Ingraham, R. Betageri, S. Jakes, Determination of Receptor-Ligand Kinetic and Equilibrium Binding Constants using Surface Plasmon Resonance: Application to the lck SH2 Domain and Phosphotyrosyl Peptides. Journal of Medicinal Chemistry. 38, 1309–1318 (1995).

58. S. Lin, A. Shih-Yuan Lee, C. C. Lin, C. K. Lee, Determination of Binding Constant and Stoichiometry for Antibody-Antigen Interaction with Surface Plasmon Resonance. Curr. Proteom. 3, 271–282 (2007).

