## Supporting Information for "Design of 8-mer Peptides that Block *Clostridioides difficile* Toxin A in Intestinal Cells"

**Materials and Methods**

1. **Materials**

N, N-dimethylformamide (DMF), dichloromethane (DCM), Alexa Fluor 594 NHS Ester, N-hydroxysuccinimide (NHS), 1-ethyl-3-(3-dimethylaminopropyl) carbodiimide hydrochloride (EDC), sodium chloride, HPLC-grade acetonitrile, HPLC-grade formic acid, LCMS-grade acetonitrile, LCMS-grade formic acid, Pierce™ Dye Removal Columns, glacial acetic acid, and hydrochloric acid were purchased from ThermoFisher Scientific (Waltham, MA). Triisopropylsilane-silane (TIS), sodium hydroxide (NaOH), phosphate-buffered saline (PBS) pH 7.4, 1,2-ethanedithiol (EDT), Tween 20, thioanisole, phenol, 3 kDa MWCO Amicon Ultra centrifugal filters, and Glucose-UDP-Fluorescein conjugate were purchased from Millipore-Sigma (Burlington, MA). 30% Hydrogen peroxide, Kaiser test kits, and Magnesium chloride were purchased from Sigma Aldrich (St. Louis, MO). 96% sulfuric acid was purchased from Macron Fine Chemicals (Randor, PA). Petroleum ether, and ethyl ether were purchased from EMD Millipore Corporation (Darmstadt, Germany). Tris HCl was purchased from IBI Scientific (Peosta, IA). Piperidine, Diisopropylethylamine (DIPEA), trifluoroacetic acid (TFA), 1-methyl-2-pyrrolidinone (NMP), 2-(7-aza-1Hbenzotriazol-1- yl) −1,1,3,3-tetramethyluronium hexafluorophosphate (HATU), Fmoc-L-Arg(Pbf)-Wang resin, Fmoc-L-Trp(Boc)-Wang resin, Fmoc-L-Thr(tBu)-Wang resin, Fmoc-L-Pro-Wang resin, Fmoc-L-Asn(Trt)-Wang resin, Rink-Amide resin, and all Fmoc protected amino acids were purchased from ChemImpex, Inc. (Wood Dale, IL). ChemMatrix Aminomethyl resin (0.7 mmol/g functional density, 100-200 mesh) was purchased from PCAS Biomatrix, Inc. (Saint-Jean-sur-Richelieu, Quebec, Canada). Toxin A from Clostridioides difficile was purchased from List Biological Labs, Inc. (Campbell, CA). Bioresistant alkanetiols hydroxyl terminated (HSC_11_(EG)_3_OH, 2-{2-[2-(1-mercaptoundec-11-yloxy)-ethoxy]-ethoxy}-ethanol) and carboxyl terminated (HSC_11_(EG)_6_OCH_2_COOH, (2-(2-(2-(2-(2-(2-(11-mercaptoundecyloxy)-ethoxy)-ethoxy)-ethoxy)-ethoxy)-ethoxy-acetic acid)) were obtained from Prochimia Surfaces (Poland). Gold sensor slides were obtained from BioNavis Ltd. (Tampere, Finland). Ethanol (200 proof) was obtained from Decon Labs, Inc (King of Prussia, PA). Milli-Q water (MQ water, resistivity > 18 MΩ cm) was obtained by using a Millipore water purification system (Billerica, MA). Nitrogen gas and liquid nitrogen were obtained from Airgas National Welders (Raleigh, NC).

2. **Computational Peptide Design**

The PepBD algorithm uses an iterative procedure that optimizes peptide sequences to bind with higher affinity and specificity to a biomolecular target than a known reference ligand. The design process is summarized below.

**(a)** *Generate input peptide:TcdA GTD structure:* The input structure for the PepBD algorithm was an 8-mer fragment of an *in-silico* peptide, NPA, identified earlier and experimentally-verified to neutralize TcdA in jejunum cells complexed with the TcdA GTD.

**(b)** *Compute initial score of random peptide:TcdA structure:* A random peptide sequence is generated and draped on the backbone scaffold of the initial peptide (NPA 8-mer) bound to the TcdA GTD and its $\Gamma_{score}$ is calculated.

**(c)** *Iteration of peptide sequence-change and conformation-change moves:* The design algorithm performs 10,000 evolution steps and generates variants of the original peptide that bind to the target protein by two kinds of moves: sequence change (mutation) and conformation change.

**(d)** *Evaluate score* $\Gamma_{score}$ *of new peptide sequence/conformer:* The score of the newly generated peptide sequence or conformer in complex with the TcdA GTD is evaluated. The score function, $\Gamma_{score}$, that we use to evaluate newly generated peptide candidates is given by:

$$\begin{aligned} \Gamma_{score}=\Delta E_{binding}+\lambda\left( E_{peptide-VDW}^{bound}+E_{peptide-ELE}^{bound}+E_{peptide-EGB}^{bound} \right)\#\left( S1 \right) \end{aligned}$$

The first term of Eq. (1), $\Delta E_{binding},$accounts for the difference in the energy of the complex and the energies of the peptide and target biomolecule prior to binding. The second term is the peptide stability term and accounts for the energy of the free peptide in the bound-state configuration. Lower scores mean better binders. The force field parameters are taken from the Amber 14SB force field.

**(e)** *Monte Carlo Metropolis Algorithm:* The Monte Carlo Metropolis algorithm is used to accept or reject new trial peptides.

More details regarding the PepBD algorithm and $\Gamma_{score}$ can be found in our previous work^1–6^.

3. **Atomistic Molecular Dynamics Simulation**

Explicit-solvent atomistic MD simulations are carried out in the canonical (NVT) ensemble using the AMBER 18 package to investigate the dynamics of the binding process between the peptide sequences and the TcdA GTD. The starting configurations of the peptide:TcdA GTD complexes in each MD simulation are the output from the searches in the PepBD algorithm. We carry out three independent simulations for each peptide:TcdA GTD complex for 100 ns to ensure that the system reaches an equilibrated state. Each peptide-receptor complex is solvated in a periodically-truncated octahedral box containing a 12 Å buffer of TIP3P water ($\sim$ 36,000 water molecules) surrounding the complex in each direction. The implicit-solvent molecular mechanics/generalized Born surface area (MM/GBSA) approach with the variable internal dielectric constant model is used to post-analyze the last 5 ns simulation trajectories of the peptide:TcdA GTD complexes to calculate the binding free energies. Details of the computational procedures and post-analysis of the atomistic MD simulations can be found in our previous work^1–6^.

**4. Bead-Based Screening Assay**

**4.1 Solid Phase Peptide Synthesis**

SA1-SA7, NPA (8-mer), and RP (8-mer) were synthesized on ChemMatrix Aminomethyl resin following a GSG linker on a Biotage Syro I peptide synthesizer (Biotage, Uppsala, Sweden) following the Fmoc/tBu protecting strategy. The GSG linker aids in displaying the peptide on the surface of the resin. The resin was swelled in DMF for 30 min. Amino acids were coupled by incubating the resin with 3 equivalents (relative to functional density of resin) protected amino acid, 3 eq. HATU, and 6 eq. DIPEA in dry DMF for 15 minutes at 45 °C. The coupling of each amino acid coupling was monitored by Kaiser test. Fmoc removal of the resin and after each amino acid conjugation was performed using 5 ml 20% v/v piperidine in DMF at room temperature for 3 min and then again for 10 min. Protected resin was stored dried under nitrogen at 4 °C. The completed peptide was deprotected immediately before use through acidolysis by incubating the resin in the deprotection cocktail, Reagent K, 82.5% v/v TFA, 5% v/v phenol, 5% v/v water, 5% v/v thioanisole, 2.5% v/v EDT for 3 hr at room temperature and under mixing. The deprotected resin was rinsed with DCM, DMF, then DCM and dried under nitrogen.

**4.2 Screening**

Peptide sequences were screened for binding to TcdA on an in-house microfluidic bead imaging system originally designed for sorting solid phase peptide libraries^7,8^. Exclusion of a fluorescent UDP-Glucose co-factor analog (UDP-Glucose-Fluorescein) from the UDP-Glucose binding pocket of TcdA was used as a proxy for visualizing peptide binding in the desired position.

Tris-buffer was removed from the TcdA solution with 3 kDa MWCO Amicon Ultra centrifugal filters, through 5 rounds of 10-fold concentration and dilution following the manufacturer’s recommended protocol. TcdA was labeled with Alexa Fluor 594 through NHS chemistry. 1 μl 10 mg/ml NHS-Alexa Fluor 594 was added to 100 μl 1 mg/ml TcdA. After 1 h incubation, unbound dye was removed with Pierce Dye Removal Columns per the manufacturer’s instructions. UDP-Glucose-Fluorescein was dissolved at 1 mg/ml in PBS, 10 mM Magnesium Chloride (Binding Buffer, BB).

Resin was incubated with 0.2 μM fluorescently labeled TcdA (TcdA-AF594) overnight. Immediately prior to screening, UDP-Glucose-Fluorescein was spiked into the resin solution at a final concentration of 0.5 μM for 15 min. The resin was gently washed 4 times with BB + 0.2% Tween 20 (Screening Buffer, SB). Resin beads were imaged in the red and green channels on the microfluidic system previously developed^7,8^. Beads were visualized on an Olympus IX81 Motorized Trinocular Inverted Fluorescence Phase Contrast Microscope fitted with FITC and RFA8 Chroma filter cubes and were imaged with a Hamamatsu C13440 camera. Each resin solution was diluted in excess SB prior to loading on the microfluidic device. Resin beads were flown through the imaging chamber one at a time and imaged in the red (TcdA-AF594) and green (UDP-Glucose-Fluorescein) channels. Approximately 30 individual beads from each peptide sequence were imaged.

Image processing and analysis was performed on the red and green channel images for each bead. Because the red channel displayed prominent halo fluorescence, the 90^th^ percentile of pixels in the bead were used for the analysis. This reduced bias from variance in the center of the bead. Additionally, the concentration of TcdA used for incubation was previously tuned to allow for a range of intensities in the halo, which allowed for differentiation between moderate and strong binders. The mean green fluorescence of the 90^th^ percentile red area and the mean 50^th^ percentile green fluorescence were calculated to determine UDP-Glucose exclusion from TcdA. The average and standard deviation of the 90^th^ percentile of the red channel and the green channel were computed.

**5. Free Peptide Synthesis and Purification for Functional Testing of SA1 on Human Gut Epithelium**

SA1-SA4 were synthesized on Wang resin pre-loaded with the first amino acid on an Initiator+ Alstra (Biotage, Uppsala, Sweden) following the Fmoc/tBu protecting strategy. The pre-loaded Wang resin was end capped with 1 ml 5M acetic anhydride in 2.5 ml 2M DIPEA for 30 minutes at room temperature. Amino acids were coupled by incubating the resin with 5 equivalents (relative to functional density of resin) protected amino acid, 5 eq. HATU, and 10 eq. DIPEA in dry DMF for 5 minutes at 75 °C. The coupling of each amino acid coupling was monitored by Kaiser test. Fmoc removal after each amino acid conjugation was performed using 5 ml 20% v/v piperidine in DMF at room temperature for 3 min and then again for 10 min. The completed peptide was cleaved and side chain protecting groups were removed through acidolysis by incubating the resin in the deprotection cocktail, Reagent K, 82.5% v/v TFA, 5% v/v phenol, 5% v/v water, 5% v/v thioanisole, 2.5% v/v EDT for 3 h at room temperature and under mixing. The deprotected peptide dissolved in the deprotection cocktail was precipitated in ice cold 50% v/v ethyl ether and 50% v/v petroleum ether (ether solution). The solution was cooled at -80 °C for 30 min, pelleted, and washed three times with ice cold ether solution. The crude peptide pellet was dried under nitrogen, redissolved in 50% v/v acetonitrile and 50% v/v water, and dried on the Biotage V-10 Touch (Biotage, Uppsala, Sweden) prior to purification and analysis.

SA1-SA4 were purified via flash chromatography on an Isolera Prime (Biotage, Uppsala, Sweden) with a Biotage Sfär Bio C18 column. Crude peptide was dissolved in 10% v/v acetonitrile, 90% v/v water, and 0.1% v/v formic acid and applied to a column samplet. Reverse phase chromatography was performed with a gradient from 5% to 70% acetonitrile in water. 0.1% formic acid was used as a modifier. Fractions were collected above a threshold 220 nm and 280 nm absorbance. Fractions were analyzed by LC-MS to identify and quantify the desired peptide. Purified peptide fractions were lyophilized. For final polishing, the peptides were dissolved in 50% v/v water and 50% v/v acetonitrile, filter sterilized, aliquoted, and lyophilized.

**6. Cell-based Assay**

**6. 1 Culture of primary colonic stem cells**. Donor Selection. Human transplant-grade donor intestines were obtained from HonorBridge (Durham, NC) and exempted from human subject’s research by the UNC Office of Human Research Ethics. Donor acceptance criteria were as follows: age 65 years or younger, brain-dead only, negative for human immunodeficiency virus, hepatitis, syphilis, tuberculosis, or COVID-19, as well as no prior history of severe abdominal injury, bowel surgery, cancer, or chemotherapy. Colonic tissue from a 34-year-old Hispanic male was used for all studies. Colonic ISCs from this donor were isolated from primary tissues and expanded as described^9^. To generate differentiated enterocytes for the toxicity assays, 2 wells of a 6-well colonic ISC expansion plate, where cells were ~90% confluent, were dissociated as described^9^ and plated on 12-well transwell inserts coated with 1% Matrigel. Colonic ISCs were expanded in colon expansion media (EM, see **reference 9** for all media formulations) until confluent (~4-days). After colonic ISCs were confluent, the media was changed to differentiation media (DM) and transepithelial electrical resistance (TEER) was measured every 24-hours using the EVOM2 (World Precision Instruments, FL). At 3-4 days of differentiation when TEER was >1000 ohms/cm^2^, the toxicity assays were initiated. All cells were incubated at 37°C in a humidified environment containing 5% CO_2._

**6.2 TcdA and SA1 exposure to differentiated human colonic epithelial cells.**

Peptides were diluted at a concentration of 1.0 mM in 500 µl of DM and added to the apical reservoir. The peptides were allowed to preincubate with the colonic monolayers for 2 hours prior to the addition of TcdA (SML1154; Sigma-Aldrich, St. Louis, MO). TcdA at 30 pMol final concentration was added directly to the apical reservoir media containing the peptide and incubated at 37°C in a humidified environment containing 5% CO_2_. TEER was measured before and after the peptides were applied. TEER was then measured at the timepoints indicated.

**7. Surface Plasmon Resonance (SPR)**

**7.1 Synthesis and purification of modified SA1 for SPR experiments**

Modified peptide SA1-K-amide was synthesized to facilitate grafting to gold sensors through the C-terminal lysine residue. Amide functionalization of the C-terminus prevented electrostatic repulsion between the C-terminus of the peptide and carboxylic acid groups on the SAM. SA1-K-amide was synthesized on Rink Amide resin on an Initiator+ Alstra (Biotage, Uppsala, Sweden) following the Fmoc/tBu protecting strategy as described above. A modified deprotection cocktail, 91.5% w/w TFA, 2.5% w/w water, 2.5% w/w TIS, 2.5% w/w DTT, and 1% w/w indole, was used. SA1-K-amide was collected, purified, and analyzed through the protocol described above.

**7.2 Cleaning gold sensors**

Gold sensor slides (12 x 20 x 0.5 mm^3^), 50 nm gold adhered on glass sensors with 2 nm chromium, were cleaned prior to use by soaking in Piranha solution (98% H_2_SO_4_ and 30% H_2_O_2_ at 3:1 v/v) for 15-20 min, followed by profuse rinsing in MQ water, rinsing in 200 proof ethanol, and drying under a stream of nitrogen. The gold sensor slides were briefly immersed in 200 proof ethanol prior to modification. (*Warning: Piranha solution reacts violently with organic materials and should be handled with extreme caution.)*

**7.3 Self-assembled monolayer (SAM) on gold surfaces followed by peptide grafting**

A mixed thiol SAM was produced by dissolving HSC_11_(EG)_3_OH and HSC_11_(EG)_6_OCH_2_COOH at a 3:1 molar ratio, 1 mM total concentration, in 200 proof ethanol. The gold sensors were immersed in the thiol solution under nitrogen and protected from light for 24 h. The surfaces were vigorously rinsed with 200 proof ethanol to disrupt multilayers, and then dried under nitrogen for subsequent characterization, peptide grafting, or SPR experiments. Modified peptide SA1-K-amide was grafted to carboxylic acid groups on the SAM through NHS/EDC chemistry. 200 mM EDC and 50 mM NHS dissolved in MQ water was applied to the surfaces of the gold sensor for 1 h. The activated gold sensors were rinsed in MQ water and dried under nitrogen. 1 mg/ml peptide SA1-K-amide was dissolved in MQ water and conjugated to the activated gold sensors for 1 h. The peptide grafted gold sensors were rinsed in MQ water and dried under nitrogen. Remaining activated carboxylic acid sites were blocked by incubating 3 M ethanolamine in MQ water on the surface of the gold sensors for 30 min. The gold sensors were rinsed in MQ water and dried under nitrogen for subsequent characterization and SPR experiments.

**7.4 Surface density of SAMs and grafted peptide**

The surface thickness of the mixed thiol SAM and SAM with grafted peptide SA1 was measured using an M-2000 DI Spectroscopic Ellipsometer (J.A. Woollam, Lincoln, NE) at three angles of incidence Φ = 55°, 65°, 75°. From the bottom up, the samples were modeled as 3 mm Cauchy substrate (glass), 2 nm chromium, and 50 nm gold per the manufacturer’s specifications. The initial value of the Cauchy substrate (polymer layer) on the gold surface was estimated to be 25 nm and the surface thickness was measured after the model was generated. Absorbance interference was mitigated by condensing the wavelength range from 600-1000 nm.

**7.5 Time-of-flight secondary ion mass spectrometry (ToF-SIMS) analysis of SPR Sensors**

Time-of-flight secondary ion mass spectrometry (ToF-SIMS) characterization of gold sensor slides with self-assembled monolayers (SAMs) and SA1 covalently grafted to SAM. Experiments were performed on an TOF SIMS V instrument (ION TOF, Inc., Chestnut Ridge, NY). Positive and negative spectra were recorded for the mixed-thiol SAM and SAM-SA1. The positive secondary ion mass spectra were calibrated using H^+^, C^+^, C_2_H_3_^+^, C_3_H_5_^+^, and C_4_H_7_^+^. The negative secondary ion mass spectra were calibrated using C^-^, O^-^, OH^-^, and C_n_^-^, respectively.

**7.6 Surface Plasmon Resonance (SPR) experiments**

A KSV SPR 200 instrument (BioNavis Instruments, Helsinki, Finland) was used to detect changes in the refractive index at the sensor interface. Sensors were equilibrated with 30 μl/min 50 mM Tris, 50 mM NaCl, 10 mM MgCl_2_ (Running Buffer, RB) for more than 5 minutes. After a stable baseline was established, 250 μl TcdA in RB at various concentrations was injected at 30 μl/min followed by RB. New sensors were used for each measurement due to the difficulty disrupting the peptide:TcdA binding interaction.

**7.7 Equilibrium and kinetic parameters**

The equilibrium dissociation constant, K_D_, was found by measuring the equilibrium net change in SPR signal (degrees) after each injection of TcdA at concentrations from approximately 0.1K_D_ to 10K_D_. The net change in degrees was converted to mass of protein adsorbed per unit area through a previously measured conversion factor, 1 degree response = 7.31 mg/m^2^. The data was fit to a Langmuir isotherm using the adsorbed mass per unit area, Q (nmol/m^2^), and the solution TcdA concentration, [TcdA] (nM),

$$\begin{aligned} Q=\frac{Q_{max}\left[ TcdA \right]}{K_{D}+\left[ TcdA \right]}\#\left( S2 \right) \end{aligned}$$

where K_D_ is the equilibrium dissociation constant (nM) and Q_max_ is the maximum binding capacity (nmol/m^2^). A Langmuir model is appropriate when there is a 1:1 interaction between the ligand and analyte, there are no mass transfer limitations, and binding events are independent. Assuming reversible binding,

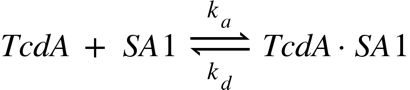
(S3)

where k_a_ is the second order association (adsorption) constant and k_d_ is the first order dissociation (desorption) constant. Assuming no mass transfer limitations, the concentration of TcdA in the bulk and at the surface is equal. The SPR response and amount adsorbed, Q, are proportional to the concentration of TcdA bound to the peptide, [TcdA•SA1]. The maximum SPR response and maximum binding capacity, Q_max_, are proportional to the maximum bound ligand [TcdA]_tot_. Substituting these into the rate equation yields,

$$\begin{aligned} \frac{dQ}{dt}=k_{a}\left[ TcdA \right]\left( Q_{max} -Q_{t} \right)-k_{d}Q_{t}\#\left( S4 \right) \end{aligned}$$

Integrating Equation S4 and substituting for k_d_,

$$\begin{aligned} Q_{t}=\frac{Q_{max}\left[ TcdA \right]}{K_{D}+ \left[ TcdA \right]}\left( 1-\frac{1}{e^{\left( \left[ TcdA \right]+K_{D} \right)k_{a}t}} \right)\#\left( S5 \right) \end{aligned}$$

The first term in Equation S5 determines the equilibrium level, and the second term determines the time to reach equilibrium. The dissociation constant can be determined from K_D_ and k_a_.

**Supplementary Text**

**Table S1.** Computationally designed 10-mer peptides reported in our previous work^10^ that neutralized toxin A in jejunum cells and reference peptide RP^11^ with their corresponding ${\Delta\Gamma}_{score}$ and ${\Delta G}_{binding}$ values.

| **Peptides** | **Sequences** | $\boldsymbol{\Gamma}_{\boldsymbol{score}}$ | $\boldsymbol{\Delta G}_{\boldsymbol{binding}}\left( \frac{\boldsymbol{kcal}}{\boldsymbol{mol}} \right)$ |
| --- | --- | --- | --- |
| RP | EGWHAHTGGG | -37.97 | -8.23 |
| NPA | DYWFQRHGHR | -41.54 | -12.81 |
| NPB | GMFWQHRRHD | -40.60 | -11.35 |
| NPC | DGWIQHYKHR | -39.36 | -6.15 |

**Table S2.** Classification of the 20 natural amino acids into six residue types according to their hydrophobicity, polarity, size and charge.

| **Residue type** | **Amino Acid** |
| --- | --- |
| Hydrophobic | Leu, Val, Ile, Met, Phe, Tyr, Trp |
| Negatively charged | Glu, Asp |
| Positively charged | Arg, Lys |
| Hydrophilic | Ser, Thr, Asn, Gln, His |
| Other | Ala, Cys, Pro |
| Glycine | Gly |

1. ***In-Silico* Screening of TcdA GTD Binding Peptides (Case 2 and Case 3)**

The lowest scoring peptides obtained from Cases 2 and 3 are SA3 and SA6. Figure S1A and S1B show examples of the plot of the Score (Γ_score_) and the RMSD profile v/s the number of sequence and conformation change moves performed in PepBD for Cases 2 and 3. The SA3:TcdA GTD catalytic domain structure obtained from the PepBD algorithm is shown in Figure S1C. Peptide SA3 has a Γ_score_ = -43.21, obtained at the 9401^th^ evolution step. Likewise, SA6 has a Γ_score_ = -44.37 which is obtained at the 7415^th^ step of the evolution process. The structure of SA6 complexed with the TcA GTD catalytic domain is shown in Figure S1D.

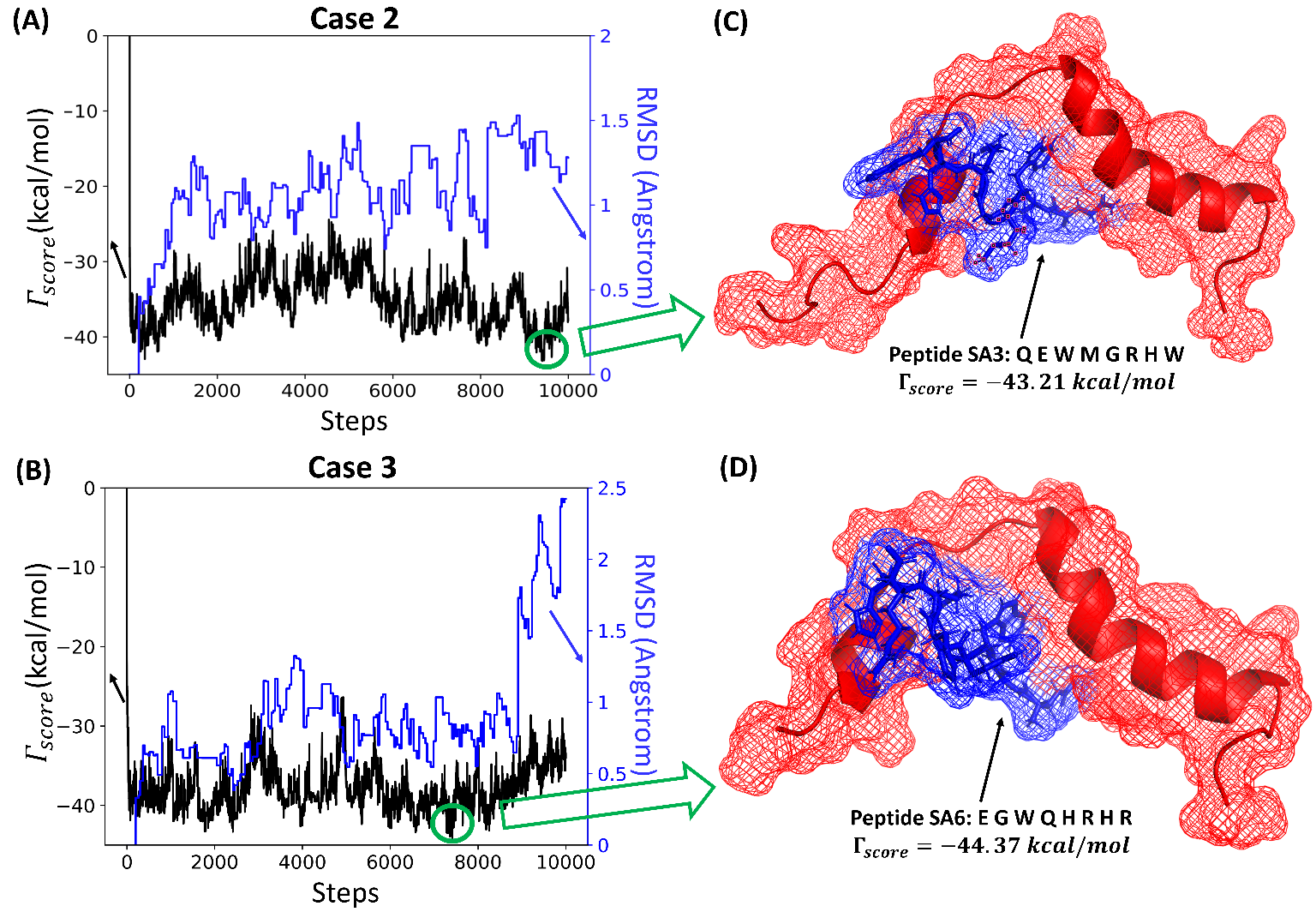

**Figure S1.** The score/RMSD vs the number of sequence and conformation steps for **(A)** Case 1 and **(B)** Case 2 with distinct initial random seeds results and their corresponding top peptides **(C)** SA3: QEWMGRHW and **(D)** SA6: EGWQHRHR obtained from PepBD.

2. **Key amino acid interactions of SA1 with TcdA GTD**

Investigation of the biorecognition mechanism of specific residues on the Toxin A neutralizing peptides will help us understand what the key amino acid interactions are on the peptide:TcdA GTD complexes and also support future efforts to design peptides targeting TcdA. When evaluated experimentally, SA1 neutralized toxin A in both primary-derived human jejunum small intestinal and colon epithelial cells and exhibited a K_D_ of 56.1 ± 29.8 nM measured by surface plasmon resonance (SPR). Hence, to draw further insights as to which SA1 amino acids play key roles in recognizing specific residues on TcdA GTD, we examine the residue-wise decomposition of the interaction energy between SA1 and the catalytic site of TcdA GTD. In Figure 3C of the *Main Text* we plot the residue-wise decomposition of the interaction energy between SA1 and the catalytic site of TcdA GTD. In Figure S2A we construct an energy panel detailing the pair wise interactions of SA1:TcdA GTD complex. Figure S2B shows the number of contacts that each residue on peptide SA1 forms with the TcdA GTD at the binding site. A contact is defined to occur when the distance between a residue on the peptide and a residue on the receptor is ≤ 4.5 Å.

Figures 3C, S2A and S2B reveal that the critical residues on SA1 involved in TcdA GTD binding are Trp3, Trp4, Arg5, Arg6, His7 and Asn8. Thus, the four amino acids: tryptophan, arginine, histidine, and asparagine on SA1 play a key role in binding to the TcdA GTD. Phe2 also contributes significantly to the interaction energy but has a slightly lower contribution than the residues mentioned above. It is also worth noting that peptide SA1 contains three aromatic residues (Phe2, Trp3, Trp4), which reflects the importance of aromatic side chains for peptide:TcdA GTD binding. Trp2 and Trp3 both form strong polar- π interaction with Asn521 while cationic Arg5 forms an ionic bond with anionic Glu517. Arg6 forms a cation-π interaction with Trp524 and interacts via coulombic forces with Asn521. His7 and Asn8 form strong polar bonds with Glu517 and Asn338 respectively. The “Number of Contacts” plot also reveals that Trp3, Arg5 and Arg6 form the most contacts with the TcdA GTD at the catalytic domain, further highlighting the importance of tryptophan and arginine in TcdA GTD binding. Glu1 on SA1 does not contribute significantly to the interaction energy but might be necessary to maintain the conformational stability of the peptide, as it can form strong anion-π interactions with neighboring Phe2, Trp3 and Trp4 on the peptide chain. It appears that the 3 three aromatic residues (Phe2, Trp3 and Trp4) also interact with each other on the peptide via π-π interactions and maintain the conformational stability of the peptide.
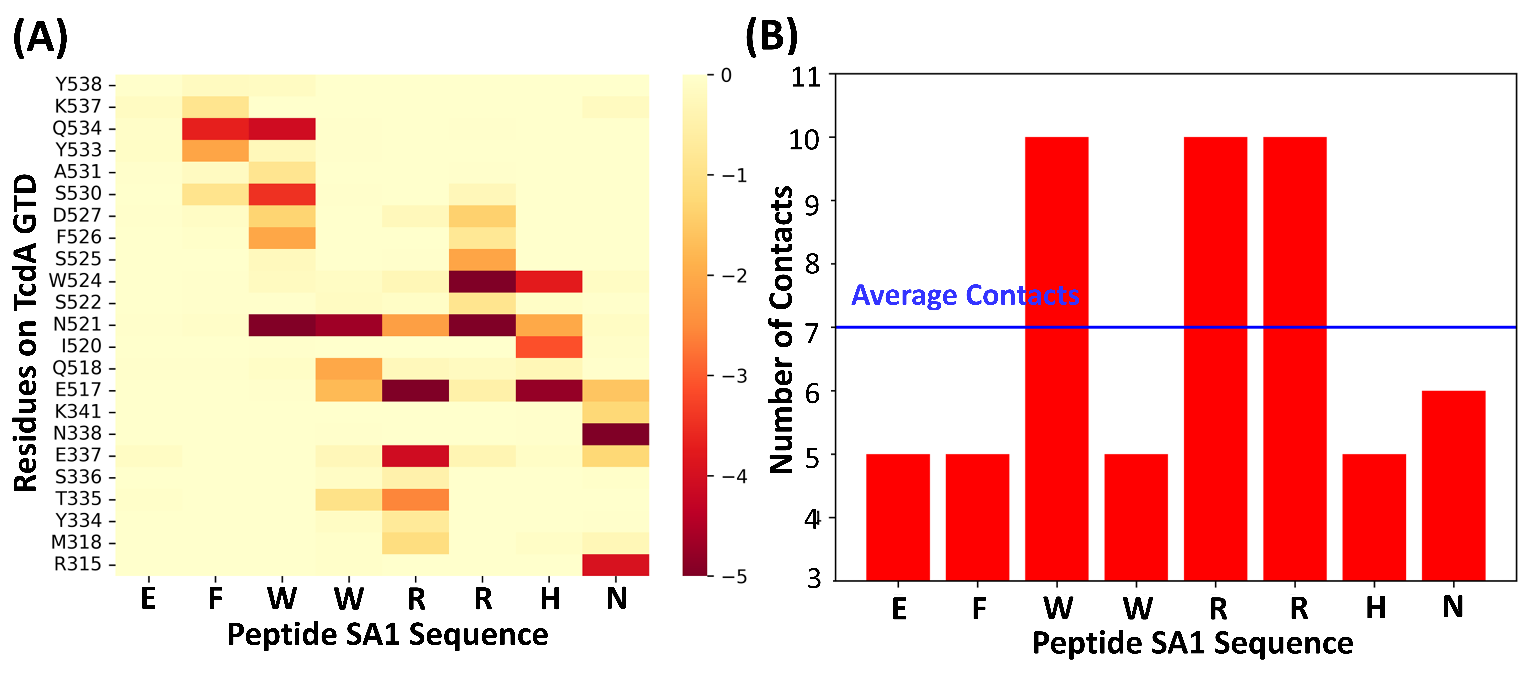
**Figure S2. (A)** Energy panel detailing the pair-wise interactions in the SA1:TcdA GTD complex. **(B)** Plot showing the Number of Contacts that each residue forms at the binding interface of the TcdA GTD.

3. **Amino acid sequence signatures in *C. diff.* Tcd A GTD binding *in-silico* peptides obtained from PepBD.**

Identifying patterns in the peptide sequences suggested via PepBD gives us an opportunity to obtain signature amino acid sequences that can uniquely bind TcdA GTD with high affinity and specificity. Such patterns offer clues as to why specific amino acids and amino acid types are preferred at certain positions along the peptide chain. PepBD generated ~15000 distinct sequences for Case 1 and ~17000 distinct sequences for Case 2. In Figure S3A, S3B and S3C, we show the sequence homology constructed using Weblogo of the top 1% of the lowest scoring distinct peptide sequences obtained from Case 1, Case 2, and Cases 1 and 2 combined, respectively. In Figure S3D, S3E and S3F, we plot the occupation % by residue type (hydrophobic, polar, positive, negative, glycine) for each site on the peptide chain for Case 1, Case 2, and Cases 1 and 2 combined, respectively. Occupation % by residue type for a site is defined as: $\frac{Number of residues of a residue type}{Total number of residues of all residue types in Top 1\% of the lowest scoring peptides}\times100$. We also plot the occupation % for the 3 most preferred residues at each site on the peptide chain for Case 1 (Figure S3G), Case 2 (Figure S3H) and Cases 1 and 2 combined (S3I). Similary, Occuptation % of a residue for a site is defined as: $\frac{Number of times a residue occurs}{Total number of residues in Top 1\% of the lowest scoring peptides}\times100$. From Figure S3 it is clear that site 3 on the peptide chain prefers Trp (W), an aromatic hydrophobic amino acid that has an occupation percentage of ~90% for Case 1 and ~100% for Case 2. Trp (W) at site 3 recruits the amino acids on TcdA GTD via both van der Waals and electrostatic interactions. The other two clear favorites are Arg (R) at site 6 and His (H) at site 7. In Case 1, where we specify that the peptide must contain two cationic residues, Arg (R) is preferred over Lys (K) for the 2^nd^ cationic residue on the peptide chain; it occupies either site 5 or 8. In both cases, where we specify that the peptide contain one anionic amino acid, Glu (E) is preferred over Asp (D). Anionic Glu (E) prefers sites 1, 2 or 4 in Case 1 and sites 1, 2 or 8 in Case 2. In Case 2, glycine prefers sites 2, 4, 5 or 8. From the *in-silico* peptide sequence data available, computational binding studies and experimental testing, we can predict amino acid sequence signatures for 8-mer peptides that can bind to *C. diff.* TcdA GTD. The consensus amino acid signature that emerges from Case 1 and Case 2 is given in Table S3.

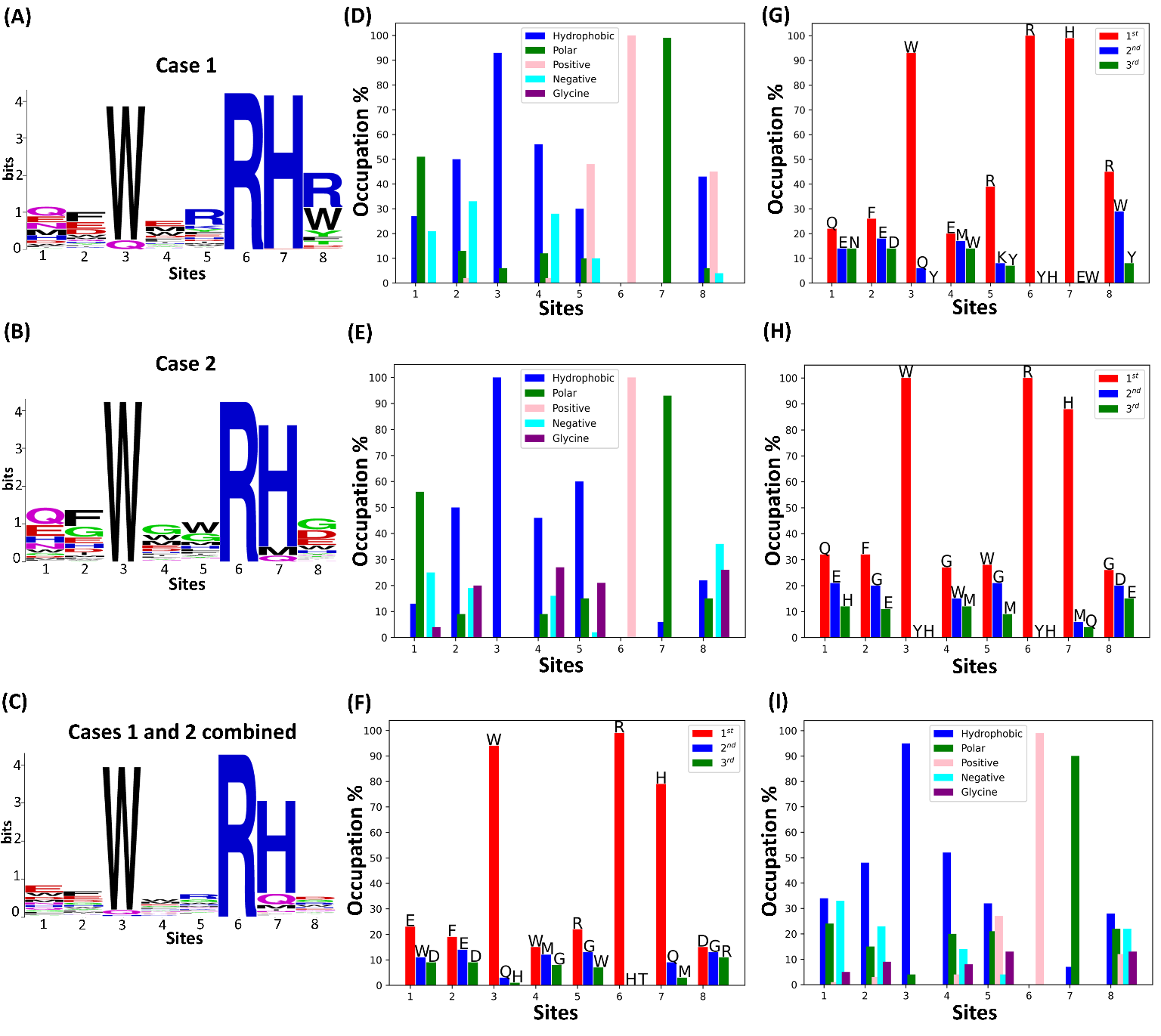

**Figure S3.** Identification of amino acid sequence pattern and signature for the top 1% of the lowest scoring distinct peptides obtained from the PepBD algorithm for each case. (A), (B) and (C): Sequence homology of the identified peptides constructed with Weblogo for Cases 1, 2, and 1 and 2 combined, respectively. (D), (E) and (F): Occupation % by residue type at each site along the peptide chain for Cases 1, 2, and 1 and 2 combined, respectively. (G), (H) and (I): Top three amino acids at each site along the peptide chain for Cases 1, 2, and 1 and 2 combined, respectively.

**Table S3.** Amino acid signature sequences derived from residue composition of top 1% of the lowest scoring peptides identified by Case 1 and Case 2.

|  | **Sites** | | | | | | | |
| --- | --- | --- | --- | --- | --- | --- | --- | --- |
| **Case** | **1** | **2** | **3** | **4** | **5** | **6** | **7** | **8** |
| 1 | Neg/Polar/Pho | Pho/Neg | W | Pho/Neg | Pos/Pho | R | H | Pho/R |
| 2 | Neg/Polar | Pho/Neg/Gly | W | Pho/Gly | Pho/Gly | R | H | X^*^ |

**^*^**Here, X can be an amino acid of any residue type

Here, Neg: Negative, Pho: Hydrophobic, Gly: Glycine, Pos: Positive

4. **Surface Characterization of SPR Sensors via Ellipsometry and Time-of-Flight Secondary Ion Mass Spectrometry**

The surface thickness of the self-assembled monolayer (SAM) and SAM with peptide SA1 were measured using ellipsometry. The model was fit over the wavelength range from 600-1000 nm to minimize the mean squared error. The mixed thiol SAM had a thickness of 3.01 ± 0.148 nm and the SAM with SA1 had a thickness of 3.76 ± 0.148 nm. The surface thickness of the mixed thiol SAM is consistent with prior work using similar alkane-oligo (ethylene glycol) thiols^12–14^, indicating tight packing of a single layer over the surface. The increase in thickness of the SAM with SA1, 7.5 Å, is consistent with the maximum diameter of SA1 estimated from the coordinate file of SA1:TcdA simulated complex, 14.6 Å, and the thickness of other short peptides layers grafted on SAMs^12–14^.

Time-of-flight secondary ion mass spectrometry (ToF-SIMS) was used to characterize the surface of the Surface Plasmon Resonance sensors. Complete coverage by the mixed-thiol self-assembled monolayer (SAM) is necessary to prevent non-specific binding of analyte protein to the gold surface. Analysis of the positive ion spectra shows the complete formation of an alkane-oligo (ethylene glycol) (OEG) SAM as expected. Figure S4A shows characteristic OEG peaks on the SAM sensor, CH_3_O^+^ (*m/z* 29), C_2_H_3_O^+^ (*m/z* 43), and C_2_H_5_O^+^ (*m/z* 45)^15^. On the SAM-SA1 sensor there is a depletion of OEG peaks and an enrichment in N-containing fragments, including CH_4_N^+^ (*m/z* 30), C_2_H_6_N^+^ (*m/z* 44), and C_4_H_8_N^+^ (*m/z* 70), indicating engraftment of SA1 onto the SAM (Figures S4A and S4B). Additionally, on the SAM-SA1 sensor there is the appearance of amino acid fragments, including glutamic acid (*m/z* 102), arginine (*m/z* 110 and 112), histidine (*m/z* 110 and 121), and tryptophan (*m/z* 130) (Figures S4C and S4D)^12,16^. The presence of these amino acid signatures further confirms the engraftment of SA1 onto the SAM.

**Figure S4.** Positive ToF-SIMS spectra within the range of m/z 0-100 of (A) mixed-thiol self-assembled monolayer (SAM) on gold and (B) SAM-SA1 (SA1 covalently grafted to SAM), and m/z 100-200 of (C) SAM, and (D) SAM-SA1. Key molecular species are labeled.

m/z

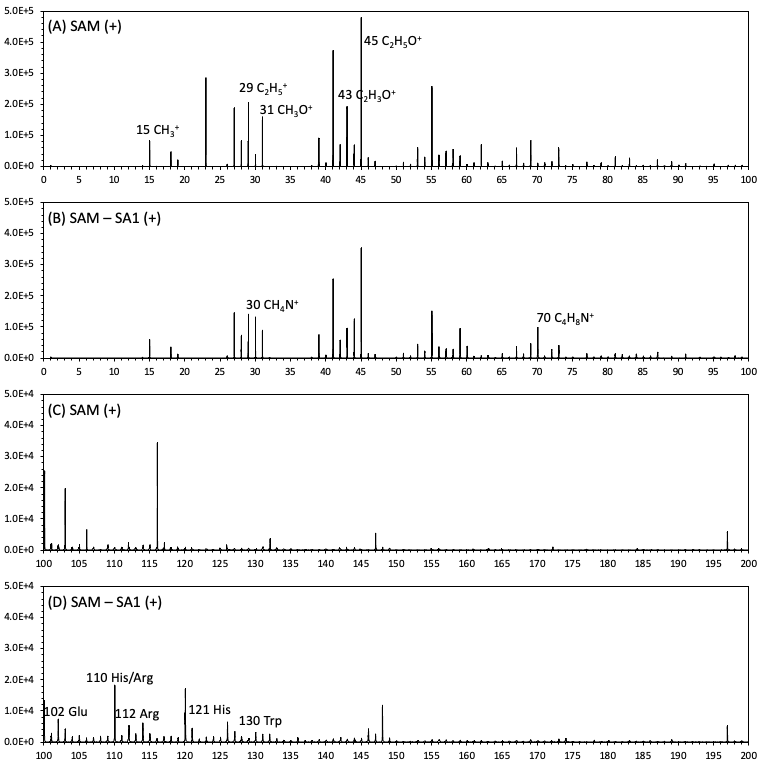

Analysis of the negative ion spectra further confirms the complete coverage by the SAM and engraftment of SA1. Figure S5A shows the characteristic peaks for SAMs constructed with OEG thiols, CHO_2_^-^ (*m/z* 45), C_2_H_3_O_2_^-^ (*m/z* 59), SO_3_^-^ (*m/z* 80), and SO_4_^-^ (*m/z* 96)^15^. Similar to the positive ion spectra, the SAM-SA1 sensor showed a depletion of the OEG peaks and an enrichment of N-containing fragments, including, CN^-^ (*m/z* 26) and CNO^-^ (*m/z* 42) (Figure S5)^15^.

**Figure S5.** Negative ToF-SIMS spectra within the range of m/z 0-100 of (A) mixed-thiol self-assembled monolayer (SAM) on gold and (B) SAM-SA1 (SA1 covalently grafted to SAM). Key molecular species are labeled.

m/z

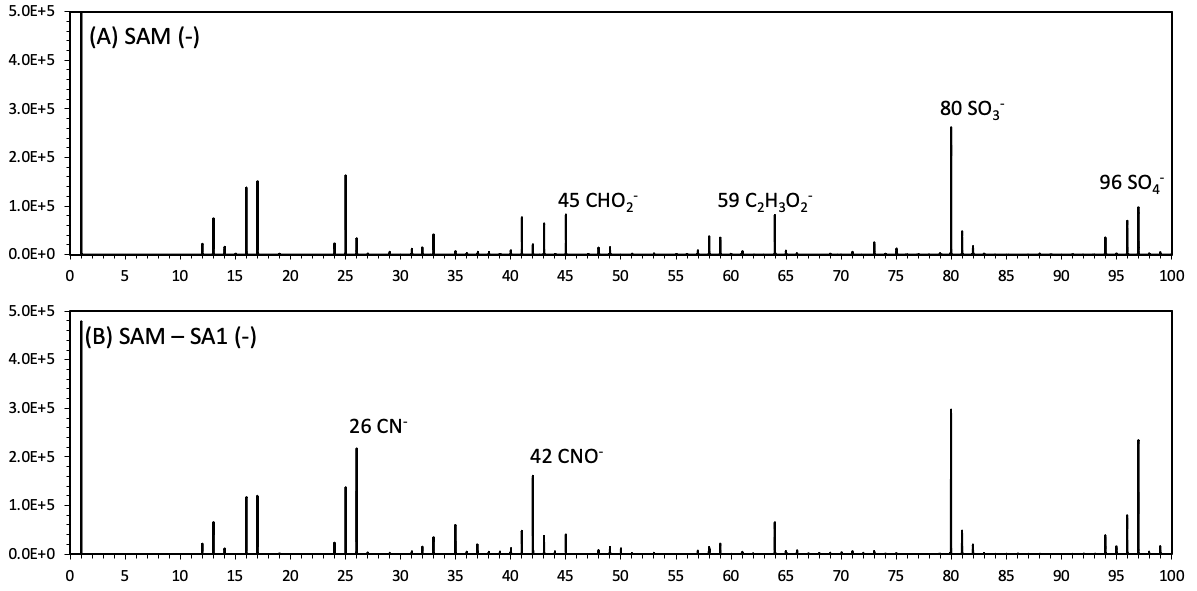

Normalized Response [degree]

| Time [min] | 300 nM |
| --- | --- |
| 0 | -0.00052781 |
| 0.050983 | -0.00059681 |
| 0.101967 | -0.00083281 |
| 0.152683 | 0.00015819 |
| 0.203684 | -0.00017681 |
| 0.254667 | -0.00101581 |
| 0.305384 | -0.00158081 |
| 0.356367 | -0.00038281 |
| 0.4071 | 0.00076919 |
| 0.4581 | -0.00048181 |
| 0.509084 | -0.00019981 |
| 0.560067 | 2.91905e-05 |
| 0.6108 | -0.00017681 |
| 0.661783 | -0.00129881 |
| 0.712517 | -0.00030681 |
| 0.762684 | 0.00010519 |
| 0.8142 | -3.98095e-05 |
| 0.865167 | -0.00023781 |
| 0.9159 | -0.00106981 |
| 0.966884 | -0.00197781 |
| 1.01787 | -0.00219081 |
| 1.06887 | -0.00103881 |
| 1.1196 | -0.00019181 |
| 1.1698 | -0.00012381 |
| 1.22107 | -0.00097081 |
| 1.27177 | 0.00131119 |
| 1.32275 | 0.0132512 |
| 1.37372 | 0.0314242 |
| 1.42447 | 0.0515202 |
| 1.47522 | 0.0701892 |
| 1.52567 | 0.0861722 |
| 1.57693 | 0.100821 |
| 1.6279 | 0.116293 |
| 1.67887 | 0.130797 |
| 1.7296 | 0.145064 |
| 1.78088 | 0.157324 |
| 1.83188 | 0.169623 |
| 1.8829 | 0.180067 |
| 1.93365 | 0.192106 |
| 1.98467 | 0.20178 |
| 2.0354 | 0.210303 |
| 2.08637 | 0.218626 |
| 2.13708 | 0.226095 |
| 2.18807 | 0.233092 |
| 2.23905 | 0.238638 |
| 2.29007 | 0.244856 |
| 2.34052 | 0.250776 |
| 2.39208 | 0.255697 |
| 2.44307 | 0.260153 |
| 2.4938 | 0.264654 |
| 2.5448 | 0.268004 |
| 2.5958 | 0.272139 |
| 2.64678 | 0.275397 |
| 2.69777 | 0.277517 |
| 2.74875 | 0.282766 |
| 2.79972 | 0.28385 |
| 2.8507 | 0.288634 |
| 2.90147 | 0.291144 |
| 2.95218 | 0.293867 |
| 3.00317 | 0.294607 |
| 3.05395 | 0.298987 |
| 3.1047 | 0.300947 |
| 3.15542 | 0.304533 |
| 3.20613 | 0.30654 |
| 3.25712 | 0.305922 |
| 3.30783 | 0.306685 |
| 3.35883 | 0.30966 |
| 3.40983 | 0.311751 |
| 3.46082 | 0.313986 |
| 3.51178 | 0.317618 |
| 3.56278 | 0.316954 |
| 3.6135 | 0.317366 |
| 3.6645 | 0.319884 |
| 3.71548 | 0.32263 |
| 3.76647 | 0.322081 |
| 3.81745 | 0.324156 |
| 3.86817 | 0.326849 |
| 3.91915 | 0.326605 |
| 3.96988 | 0.329458 |
| 4.02085 | 0.334471 |
| 4.07157 | 0.330107 |
| 4.12255 | 0.332831 |
| 4.17353 | 0.334784 |
| 4.22452 | 0.334677 |
| 4.27552 | 0.337881 |
| 4.32625 | 0.339987 |
| 4.37697 | 0.33853 |
| 4.42795 | 0.340773 |
| 4.47893 | 0.342085 |
| 4.52965 | 0.340841 |
| 4.58037 | 0.34397 |
| 4.63082 | 0.344542 |
| 4.68237 | 0.344809 |
| 4.73307 | 0.346831 |
| 4.78432 | 0.347449 |
| 4.83528 | 0.348898 |
| 4.88632 | 0.349623 |
| 4.9373 | 0.35034 |
| 4.98828 | 0.350355 |
| 5.03928 | 0.353193 |
| 5.09028 | 0.353209 |
| 5.14098 | 0.353079 |
| 5.19198 | 0.353377 |
| 5.24242 | 0.355429 |
| 5.29367 | 0.356924 |
| 5.34438 | 0.357176 |
| 5.39512 | 0.357184 |
| 5.44582 | 0.357969 |
| 5.49682 | 0.359305 |
| 5.54753 | 0.358732 |
| 5.59883 | 0.360869 |
| 5.64955 | 0.359854 |
| 5.7003 | 0.360365 |
| 5.75102 | 0.360472 |
| 5.80175 | 0.362158 |
| 5.8522 | 0.362616 |
| 5.90347 | 0.363127 |
| 5.95445 | 0.36244 |
| 6.00533 | 0.361906 |
| 6.05632 | 0.361571 |
| 6.10732 | 0.363318 |
| 6.1583 | 0.36228 |
| 6.20927 | 0.36363 |
| 6.26002 | 0.363989 |
| 6.31128 | 0.362997 |
| 6.36227 | 0.363646 |
| 6.41328 | 0.363722 |
| 6.46427 | 0.362715 |
| 6.51525 | 0.363318 |
| 6.56623 | 0.364149 |
| 6.61725 | 0.36434 |
| 6.66822 | 0.364058 |
| 6.71922 | 0.362188 |
| 6.76993 | 0.363814 |
| 6.82065 | 0.362463 |
| 6.87135 | 0.361235 |
| 6.92262 | 0.364065 |
| 6.97333 | 0.363295 |
| 7.02407 | 0.363196 |
| 7.07503 | 0.36373 |
| 7.12603 | 0.362188 |
| 7.17727 | 0.363066 |
| 7.22828 | 0.363196 |
| 7.27925 | 0.36289 |
| 7.33 | 0.361479 |
| 7.38098 | 0.360113 |
| 7.43172 | 0.362745 |
| 7.48323 | 0.361464 |
| 7.53422 | 0.364851 |
| 7.5847 | 0.360617 |
| 7.63648 | 0.361868 |
| 7.68722 | 0.360373 |
| 7.73822 | 0.360655 |
| 7.78922 | 0.360068 |
| 7.83993 | 0.359663 |
| 7.8909 | 0.360823 |
| 7.94163 | 0.360869 |
| 7.99237 | 0.359305 |
| 8.04308 | 0.359396 |
| 8.09332 | 0.359228 |
| 8.14483 | 0.358709 |
| 8.19562 | 0.358298 |
| 8.2466 | 0.359656 |
| 8.29757 | 0.358305 |
| 8.34853 | 0.358069 |
| 8.39927 | 0.358366 |
| 8.45025 | 0.357565 |
| 8.50122 | 0.356947 |
| 8.5522 | 0.356894 |
| 8.60318 | 0.357069 |
| 8.65418 | 0.356115 |
| 8.70488 | 0.356169 |
| 8.75562 | 0.356123 |
| 8.8066 | 0.355444 |
| 8.85758 | 0.354002 |
| 8.90842 | 0.353308 |
| 8.95865 | 0.352987 |
| 9.01023 | 0.354025 |
| 9.0613 | 0.351866 |
| 9.11202 | 0.355147 |
| 9.16302 | 0.352659 |
| 9.21373 | 0.35356 |
| 9.26473 | 0.351034 |
| 9.31552 | 0.350363 |
| 9.36625 | 0.350256 |
| 9.41725 | 0.350104 |
| 9.46798 | 0.350943 |
| 9.51895 | 0.349714 |
| 9.56995 | 0.350332 |
| 9.62092 | 0.350432 |
| 9.67192 | 0.349135 |
| 9.72288 | 0.348814 |
| 9.77362 | 0.34825 |
| 9.82435 | 0.349325 |
| 9.87508 | 0.347586 |
| 9.92555 | 0.347166 |
| 9.97657 | 0.348066 |
| 10.0273 | 0.347143 |
| 10.0785 | 0.347304 |
| 10.1295 | 0.346075 |

| Time [min] | 250 nM |
| --- | --- |
| 0 | -0.00026444 |
| 0.050717 | -0.00077544 |
| 0.101717 | -0.00073044 |
| 0.1527 | -0.00095144 |
| 0.203433 | -0.00142444 |
| 0.254433 | -0.00223344 |
| 0.305417 | -0.00105844 |
| 0.3564 | -0.00285844 |
| 0.4074 | -0.00285844 |
| 0.458367 | -0.00288944 |
| 0.508583 | -0.00311844 |
| 0.560617 | -0.00190544 |
| 0.6108 | -0.00270644 |
| 0.66205 | -0.00270644 |
| 0.712833 | -0.00472044 |
| 0.763567 | -0.00288944 |
| 0.814267 | -0.00410244 |
| 0.86525 | -0.00557444 |
| 0.916233 | -0.00429344 |
| 0.96695 | -0.00507844 |
| 1.01718 | -0.00377444 |
| 1.06868 | -0.00550644 |
| 1.11913 | -0.00603244 |
| 1.17038 | -0.00671144 |
| 1.22138 | -0.00391944 |
| 1.27237 | -0.00322444 |
| 1.32308 | 0.00761656 |
| 1.37408 | 0.0273686 |
| 1.42507 | 0.0529806 |
| 1.47578 | 0.0779446 |
| 1.5265 | 0.10094 |
| 1.57722 | 0.121058 |
| 1.62822 | 0.137408 |
| 1.67895 | 0.155253 |
| 1.72997 | 0.172931 |
| 1.78067 | 0.190005 |
| 1.83168 | 0.204951 |
| 1.8824 | 0.217082 |
| 1.93287 | 0.230983 |
| 1.98412 | 0.242488 |
| 2.03537 | 0.256907 |
| 2.08612 | 0.265635 |
| 2.13708 | 0.277408 |
| 2.18782 | 0.281207 |
| 2.23855 | 0.291568 |
| 2.28927 | 0.297412 |
| 2.33997 | 0.30514 |
| 2.39072 | 0.312976 |
| 2.44143 | 0.317309 |
| 2.49242 | 0.32149 |
| 2.54317 | 0.327517 |
| 2.59388 | 0.332438 |
| 2.64487 | 0.337497 |
| 2.69585 | 0.342051 |
| 2.74657 | 0.343821 |
| 2.79755 | 0.349353 |
| 2.84828 | 0.354335 |
| 2.899 | 0.35644 |
| 2.94998 | 0.363154 |
| 3.00097 | 0.364917 |
| 3.05168 | 0.36822 |
| 3.10242 | 0.372218 |
| 3.1534 | 0.372882 |
| 3.20438 | 0.375247 |
| 3.25538 | 0.37955 |
| 3.30637 | 0.378375 |
| 3.35735 | 0.382037 |
| 3.40833 | 0.388499 |
| 3.45933 | 0.387858 |
| 3.51033 | 0.390803 |
| 3.56107 | 0.391734 |
| 3.6123 | 0.394961 |
| 3.6631 | 0.397281 |
| 3.71408 | 0.398051 |
| 3.7648 | 0.403041 |
| 3.81552 | 0.406283 |
| 3.86623 | 0.405284 |
| 3.91722 | 0.406863 |
| 3.96822 | 0.407367 |
| 4.01893 | 0.410525 |
| 4.06993 | 0.410029 |
| 4.12092 | 0.41788 |
| 4.1719 | 0.417415 |
| 4.22262 | 0.416842 |
| 4.2736 | 0.418544 |
| 4.32458 | 0.421054 |
| 4.37528 | 0.422984 |
| 4.42627 | 0.428225 |
| 4.477 | 0.423556 |
| 4.52772 | 0.426104 |
| 4.57818 | 0.427958 |
| 4.6297 | 0.430766 |
| 4.68072 | 0.431605 |
| 4.73168 | 0.434512 |
| 4.78268 | 0.434848 |
| 4.83367 | 0.435023 |
| 4.88465 | 0.436114 |
| 4.93537 | 0.438334 |
| 4.98635 | 0.438922 |
| 5.03707 | 0.441729 |
| 5.08805 | 0.44205 |
| 5.13878 | 0.44308 |
| 5.1895 | 0.44411 |
| 5.24048 | 0.443049 |
| 5.29148 | 0.447314 |
| 5.34245 | 0.447261 |
| 5.39345 | 0.448382 |
| 5.44442 | 0.450992 |
| 5.49518 | 0.451686 |
| 5.54595 | 0.450435 |
| 5.59667 | 0.454356 |
| 5.64765 | 0.455142 |
| 5.69865 | 0.453921 |
| 5.7491 | 0.454539 |
| 5.8001 | 0.455241 |
| 5.85107 | 0.458293 |
| 5.90212 | 0.461337 |
| 5.95238 | 0.458804 |
| 6.00395 | 0.461741 |
| 6.05493 | 0.458468 |
| 6.10593 | 0.460406 |
| 6.1569 | 0.462321 |
| 6.20763 | 0.466289 |
| 6.25833 | 0.464175 |
| 6.30938 | 0.461032 |
| 6.35985 | 0.46345 |
| 6.41085 | 0.463183 |
| 6.46182 | 0.464137 |
| 6.51208 | 0.467296 |
| 6.5636 | 0.464961 |
| 6.61458 | 0.465686 |
| 6.66558 | 0.466647 |
| 6.71657 | 0.466731 |
| 6.76757 | 0.468219 |
| 6.81827 | 0.466792 |
| 6.8695 | 0.467334 |
| 6.91997 | 0.469462 |
| 6.97147 | 0.469836 |
| 7.0222 | 0.469516 |
| 7.0732 | 0.471782 |
| 7.12432 | 0.469661 |
| 7.1745 | 0.47111 |
| 7.22522 | 0.469829 |
| 7.27648 | 0.471782 |
| 7.32747 | 0.470691 |
| 7.37845 | 0.471484 |
| 7.42945 | 0.472934 |
| 7.47988 | 0.470645 |
| 7.53063 | 0.471881 |
| 7.58162 | 0.469447 |
| 7.63263 | 0.472285 |
| 7.68333 | 0.472789 |
| 7.73405 | 0.471049 |
| 7.78503 | 0.475512 |
| 7.83577 | 0.472712 |
| 7.88675 | 0.471148 |
| 7.93722 | 0.471385 |
| 7.98847 | 0.476481 |
| 8.0392 | 0.469088 |
| 8.09017 | 0.471797 |
| 8.14115 | 0.473208 |
| 8.19187 | 0.473575 |
| 8.24287 | 0.471171 |
| 8.29358 | 0.476481 |
| 8.34433 | 0.473567 |
| 8.39532 | 0.473643 |
| 8.44628 | 0.47301 |
| 8.49732 | 0.475764 |
| 8.54805 | 0.472873 |
| 8.59908 | 0.476352 |
| 8.6498 | 0.472514 |
| 8.70052 | 0.474025 |
| 8.75127 | 0.473514 |
| 8.80223 | 0.47256 |
| 8.85325 | 0.471553 |
| 8.90453 | 0.470889 |
| 8.95523 | 0.471324 |
| 9.00625 | 0.471293 |
| 9.05698 | 0.472979 |
| 9.10825 | 0.469714 |
| 9.15925 | 0.474223 |
| 9.2105 | 0.469455 |
| 9.2615 | 0.473742 |
| 9.31247 | 0.474261 |
| 9.36347 | 0.471362 |
| 9.41418 | 0.470553 |
| 9.46518 | 0.475467 |
| 9.51615 | 0.470424 |
| 9.56713 | 0.46934 |
| 9.61817 | 0.469417 |
| 9.66917 | 0.473231 |
| 9.72013 | 0.470492 |
| 9.77088 | 0.470164 |
| 9.82187 | 0.467654 |
| 9.87287 | 0.466327 |
| 9.9236 | 0.469676 |
| 9.9746 | 0.464045 |
| 10.0256 | 0.468898 |
| 10.0766 | 0.467105 |
| 10.1273 | 0.467868 |

| Time [min] | 200 nM |
| --- | --- |
| 0 | -0.000131444 |
| 0.050716 | 0.00189056 |
| 0.101699 | -0.000124444 |
| 0.152683 | -0.000276444 |
| 0.203683 | 0.00179856 |
| 0.254383 | 0.00108856 |
| 0.305366 | -0.00190944 |
| 0.355817 | -0.00148944 |
| 0.407083 | -0.000627444 |
| 0.458067 | -0.000230444 |
| 0.50905 | 0.000364556 |
| 0.560033 | 0.00114256 |
| 0.61075 | 0.000264556 |
| 0.661501 | -0.000299444 |
| 0.712217 | -0.00103944 |
| 0.763467 | 0.000333556 |
| 0.814199 | -0.000673444 |
| 0.865182 | 3.65556e-05 |
| 0.915916 | 0.000493556 |
| 0.966884 | -0.000177444 |
| 1.01788 | -0.000543444 |
| 1.06862 | -4.74444e-05 |
| 1.11933 | -0.000108444 |
| 1.17033 | 0.000318556 |
| 1.22142 | 0.000287556 |
| 1.2724 | 0.00275956 |
| 1.32313 | 0.0110526 |
| 1.37412 | 0.0270896 |
| 1.42512 | 0.0422876 |
| 1.47608 | 0.0567226 |
| 1.52655 | 0.0707756 |
| 1.57778 | 0.0848216 |
| 1.62823 | 0.0999736 |
| 1.67922 | 0.113173 |
| 1.72997 | 0.127432 |
| 1.78122 | 0.142287 |
| 1.83222 | 0.154982 |
| 1.8832 | 0.165648 |
| 1.93418 | 0.178183 |
| 1.98517 | 0.186995 |
| 2.03588 | 0.196677 |
| 2.08687 | 0.206732 |
| 2.13785 | 0.213782 |
| 2.18805 | 0.221533 |
| 2.23903 | 0.228789 |
| 2.29055 | 0.235007 |
| 2.34153 | 0.241354 |
| 2.39252 | 0.245451 |
| 2.44377 | 0.252333 |
| 2.49475 | 0.256941 |
| 2.54573 | 0.261549 |
| 2.5967 | 0.266463 |
| 2.64743 | 0.269614 |
| 2.69815 | 0.271948 |
| 2.74887 | 0.277228 |
| 2.79987 | 0.279356 |
| 2.85087 | 0.282469 |
| 2.90132 | 0.285948 |
| 2.95283 | 0.288527 |
| 3.00382 | 0.291296 |
| 3.05453 | 0.297766 |
| 3.10525 | 0.296805 |
| 3.15623 | 0.29891 |
| 3.20724 | 0.301634 |
| 3.25797 | 0.303404 |
| 3.3095 | 0.305922 |
| 3.3605 | 0.310202 |
| 3.41148 | 0.313109 |
| 3.46275 | 0.313849 |
| 3.5132 | 0.314719 |
| 3.5642 | 0.318747 |
| 3.61493 | 0.320883 |
| 3.66567 | 0.321173 |
| 3.71638 | 0.323637 |
| 3.76739 | 0.325323 |
| 3.8181 | 0.325613 |
| 3.86909 | 0.328352 |
| 3.92005 | 0.327444 |
| 3.97103 | 0.33119 |
| 4.02202 | 0.333357 |
| 4.073 | 0.334944 |
| 4.124 | 0.333647 |
| 4.17498 | 0.337576 |
| 4.226 | 0.339491 |
| 4.2767 | 0.341292 |
| 4.32768 | 0.34191 |
| 4.37865 | 0.339873 |
| 4.42965 | 0.343344 |
| 4.48062 | 0.344496 |
| 4.5316 | 0.345946 |
| 4.5826 | 0.346815 |
| 4.63359 | 0.348959 |
| 4.68433 | 0.348898 |
| 4.73505 | 0.349852 |
| 4.78523 | 0.348555 |
| 4.83678 | 0.35195 |
| 4.88779 | 0.353598 |
| 4.9385 | 0.355978 |
| 4.9895 | 0.357855 |
| 5.03973 | 0.355307 |
| 5.09125 | 0.354384 |
| 5.14223 | 0.359618 |
| 5.19325 | 0.360113 |
| 5.24422 | 0.359816 |
| 5.29497 | 0.358206 |
| 5.34567 | 0.361464 |
| 5.3964 | 0.360793 |
| 5.44737 | 0.36286 |
| 5.4981 | 0.363737 |
| 5.54909 | 0.361586 |
| 5.60007 | 0.366316 |
| 5.65105 | 0.365271 |
| 5.70205 | 0.367285 |
| 5.75302 | 0.367529 |
| 5.804 | 0.366209 |
| 5.85498 | 0.363814 |
| 5.90596 | 0.367079 |
| 5.95724 | 0.368201 |
| 6.0082 | 0.367636 |
| 6.05892 | 0.367369 |
| 6.1099 | 0.368582 |
| 6.16088 | 0.367087 |
| 6.21188 | 0.368109 |
| 6.26287 | 0.368513 |
| 6.31358 | 0.366759 |
| 6.3643 | 0.367415 |
| 6.41503 | 0.366499 |
| 6.46602 | 0.370123 |
| 6.51673 | 0.36878 |
| 6.56747 | 0.370077 |
| 6.61819 | 0.369841 |
| 6.66918 | 0.369261 |
| 6.72017 | 0.372321 |
| 6.77115 | 0.370139 |
| 6.82213 | 0.367873 |
| 6.87312 | 0.36785 |
| 6.92383 | 0.369871 |
| 6.97482 | 0.369032 |
| 7.02582 | 0.367491 |
| 7.0768 | 0.369086 |
| 7.12753 | 0.371916 |
| 7.17825 | 0.36949 |
| 7.22925 | 0.367728 |
| 7.28023 | 0.369536 |
| 7.33122 | 0.368864 |
| 7.3822 | 0.369032 |
| 7.43293 | 0.369452 |
| 7.48365 | 0.369124 |
| 7.53465 | 0.368323 |
| 7.58563 | 0.368117 |
| 7.63662 | 0.366782 |
| 7.6876 | 0.367064 |
| 7.7386 | 0.366568 |
| 7.78958 | 0.366408 |
| 7.84057 | 0.366225 |
| 7.89155 | 0.368689 |
| 7.94227 | 0.366865 |
| 7.99325 | 0.366011 |
| 8.04425 | 0.367934 |
| 8.09522 | 0.365332 |
| 8.1462 | 0.366804 |
| 8.19692 | 0.36772 |
| 8.24818 | 0.367606 |
| 8.29892 | 0.366263 |
| 8.34988 | 0.366042 |
| 8.40086 | 0.364493 |
| 8.4516 | 0.364584 |
| 8.50258 | 0.366804 |
| 8.55358 | 0.364775 |
| 8.60457 | 0.365271 |
| 8.65555 | 0.363783 |
| 8.70628 | 0.366049 |
| 8.75698 | 0.364767 |
| 8.80772 | 0.36283 |
| 8.85846 | 0.362402 |
| 8.90919 | 0.362936 |
| 8.96023 | 0.363753 |
| 9.01097 | 0.362273 |
| 9.06197 | 0.362051 |
| 9.11267 | 0.36196 |
| 9.16365 | 0.361952 |
| 9.21438 | 0.361159 |
| 9.26537 | 0.361288 |
| 9.31635 | 0.36106 |
| 9.36737 | 0.362311 |
| 9.41807 | 0.361128 |
| 9.46887 | 0.36006 |
| 9.51958 | 0.36273 |
| 9.57035 | 0.35993 |
| 9.62139 | 0.358122 |
| 9.6721 | 0.359381 |
| 9.72282 | 0.359351 |
| 9.77382 | 0.360228 |
| 9.8248 | 0.359389 |
| 9.87552 | 0.360266 |
| 9.92622 | 0.359892 |
| 9.9775 | 0.358626 |
| 10.0285 | 0.358443 |
| 10.0792 | 0.359038 |
| 10.1302 | 0.358786 |

| Time [min] | 150 nM |
| --- | --- |
| 0 | -2.952e-05 |
| 0.050717 | -0.00083852 |
| 0.101717 | 0.00055748 |
| 0.1527 | -0.00041052 |
| 0.203433 | 8.48e-06 |
| 0.254433 | 0.00016148 |
| 0.305417 | -0.00051752 |
| 0.3564 | -0.00025052 |
| 0.4074 | 0.00045848 |
| 0.458367 | 6.248e-05 |
| 0.508583 | -0.00034252 |
| 0.560617 | -0.00075452 |
| 0.6108 | 0.00038248 |
| 0.66205 | 1.648e-05 |
| 0.712833 | 0.00035948 |
| 0.763567 | -0.00038852 |
| 0.814267 | -0.00070852 |
| 0.86525 | 0.00048948 |
| 0.916233 | -0.00029652 |
| 0.96695 | -0.00074652 |
| 1.01718 | 0.00035148 |
| 1.06868 | 9.248e-05 |
| 1.11913 | -0.00031952 |
| 1.17038 | -0.00033452 |
| 1.22138 | 6.948e-05 |
| 1.27237 | 0.00070248 |
| 1.32308 | 0.0101865 |
| 1.37408 | 0.0247965 |
| 1.42507 | 0.0389875 |
| 1.47578 | 0.0532085 |
| 1.5265 | 0.0671245 |
| 1.57722 | 0.0815595 |
| 1.62822 | 0.0929425 |
| 1.67895 | 0.10669 |
| 1.72997 | 0.119057 |
| 1.78067 | 0.131104 |
| 1.83168 | 0.139428 |
| 1.8824 | 0.153107 |
| 1.93287 | 0.161164 |
| 1.98412 | 0.171143 |
| 2.03537 | 0.181397 |
| 2.08612 | 0.190163 |
| 2.13708 | 0.198365 |
| 2.18782 | 0.205987 |
| 2.23855 | 0.2128 |
| 2.28927 | 0.219887 |
| 2.33997 | 0.226632 |
| 2.39072 | 0.232735 |
| 2.44143 | 0.238152 |
| 2.49242 | 0.243905 |
| 2.54317 | 0.249116 |
| 2.59388 | 0.253624 |
| 2.64487 | 0.25824 |
| 2.69585 | 0.262101 |
| 2.74657 | 0.26664 |
| 2.79755 | 0.269959 |
| 2.84828 | 0.27327 |
| 2.899 | 0.27784 |
| 2.94998 | 0.281174 |
| 3.00097 | 0.283928 |
| 3.05168 | 0.287095 |
| 3.10242 | 0.289765 |
| 3.1534 | 0.293541 |
| 3.20438 | 0.295823 |
| 3.25538 | 0.298371 |
| 3.30637 | 0.301102 |
| 3.35735 | 0.303101 |
| 3.40833 | 0.306 |
| 3.45933 | 0.308564 |
| 3.51033 | 0.310387 |
| 3.56107 | 0.312378 |
| 3.6123 | 0.315194 |
| 3.6631 | 0.316895 |
| 3.71408 | 0.319474 |
| 3.7648 | 0.320107 |
| 3.81552 | 0.32264 |
| 3.86623 | 0.325028 |
| 3.91722 | 0.326409 |
| 3.96822 | 0.328209 |
| 4.01893 | 0.330514 |
| 4.06993 | 0.331261 |
| 4.12092 | 0.333054 |
| 4.1719 | 0.334763 |
| 4.22262 | 0.336655 |
| 4.2736 | 0.338219 |
| 4.32458 | 0.339348 |
| 4.37528 | 0.341279 |
| 4.42627 | 0.342042 |
| 4.477 | 0.343751 |
| 4.52772 | 0.345437 |
| 4.57818 | 0.34668 |
| 4.6297 | 0.348168 |
| 4.68072 | 0.349412 |
| 4.73168 | 0.350464 |
| 4.78268 | 0.351769 |
| 4.83367 | 0.35273 |
| 4.88465 | 0.354706 |
| 4.93537 | 0.355065 |
| 4.98635 | 0.356568 |
| 5.03707 | 0.357743 |
| 5.08805 | 0.358742 |
| 5.13878 | 0.360032 |
| 5.1895 | 0.361336 |
| 5.24048 | 0.362641 |
| 5.29148 | 0.363419 |
| 5.34245 | 0.364174 |
| 5.39345 | 0.36541 |
| 5.44442 | 0.366356 |
| 5.49518 | 0.367325 |
| 5.54595 | 0.367981 |
| 5.59667 | 0.368088 |
| 5.64765 | 0.369225 |
| 5.69865 | 0.369889 |
| 5.7491 | 0.371148 |
| 5.8001 | 0.371995 |
| 5.85107 | 0.372246 |
| 5.90212 | 0.372536 |
| 5.95238 | 0.372696 |
| 6.00395 | 0.374093 |
| 6.05493 | 0.375062 |
| 6.10593 | 0.375939 |
| 6.1569 | 0.376206 |
| 6.20763 | 0.377221 |
| 6.25833 | 0.377831 |
| 6.30938 | 0.378373 |
| 6.35985 | 0.37954 |
| 6.41085 | 0.379433 |
| 6.46182 | 0.379189 |
| 6.51208 | 0.379815 |
| 6.5636 | 0.380761 |
| 6.61458 | 0.381379 |
| 6.66558 | 0.381432 |
| 6.71657 | 0.381592 |
| 6.76757 | 0.382584 |
| 6.81827 | 0.383027 |
| 6.8695 | 0.382599 |
| 6.91997 | 0.383111 |
| 6.97147 | 0.383637 |
| 7.0222 | 0.384125 |
| 7.0732 | 0.383408 |
| 7.12432 | 0.384148 |
| 7.1745 | 0.383973 |
| 7.22522 | 0.384621 |
| 7.27648 | 0.383713 |
| 7.32747 | 0.385407 |
| 7.37845 | 0.385911 |
| 7.42945 | 0.385598 |
| 7.47988 | 0.386223 |
| 7.53063 | 0.38617 |
| 7.58162 | 0.385544 |
| 7.63263 | 0.386345 |
| 7.68333 | 0.387368 |
| 7.73405 | 0.386658 |
| 7.78503 | 0.386841 |
| 7.83577 | 0.387101 |
| 7.88675 | 0.38794 |
| 7.93722 | 0.386902 |
| 7.98847 | 0.386674 |
| 8.0392 | 0.387238 |
| 8.09017 | 0.386231 |
| 8.14115 | 0.387818 |
| 8.19187 | 0.387375 |
| 8.24287 | 0.387162 |
| 8.29358 | 0.386971 |
| 8.34433 | 0.386414 |
| 8.39532 | 0.386757 |
| 8.44628 | 0.386704 |
| 8.49732 | 0.385903 |
| 8.54805 | 0.38659 |
| 8.59908 | 0.386117 |
| 8.6498 | 0.386544 |
| 8.70052 | 0.386056 |
| 8.75127 | 0.386231 |
| 8.80223 | 0.385888 |
| 8.85325 | 0.386239 |
| 8.90453 | 0.386208 |
| 8.95523 | 0.387528 |
| 9.00625 | 0.386109 |
| 9.05698 | 0.387505 |
| 9.10825 | 0.387406 |
| 9.15925 | 0.387291 |
| 9.2105 | 0.38649 |
| 9.2615 | 0.386101 |
| 9.31247 | 0.385979 |
| 9.36347 | 0.385834 |
| 9.41418 | 0.386155 |
| 9.46518 | 0.385964 |
| 9.51615 | 0.385499 |
| 9.56713 | 0.385994 |
| 9.61817 | 0.386147 |
| 9.66917 | 0.38559 |
| 9.72013 | 0.38588 |
| 9.77088 | 0.385209 |
| 9.82187 | 0.385178 |
| 9.87287 | 0.385094 |
| 9.9236 | 0.387246 |
| 9.9746 | 0.384804 |
| 10.0256 | 0.386445 |
| 10.0766 | 0.386513 |
| 10.1273 | 0.386475 |

| Time [min] | 100 nM |
| --- | --- |
| 0 | -0.00024105 |
| 0.050717 | 0.00027795 |
| 0.1017 | -0.00043905 |
| 0.15245 | 0.00039195 |
| 0.203433 | 4.895e-05 |
| 0.2544 | 0.00072795 |
| 0.30515 | -0.00024905 |
| 0.35585 | -1.205e-05 |
| 0.406317 | 0.00046095 |
| 0.4573 | 0.00060595 |
| 0.508283 | 0.00067395 |
| 0.55855 | -0.00030205 |
| 0.609533 | 0.00030795 |
| 0.660817 | 0.00019395 |
| 0.7118 | -8.105e-05 |
| 0.762767 | 7.195e-05 |
| 0.8135 | 5.695e-05 |
| 0.8642 | 0.00041495 |
| 0.914933 | -0.00014905 |
| 0.9659 | 0.00065195 |
| 1.01663 | 0.00052995 |
| 1.06765 | 0.00025495 |
| 1.11863 | 0.00011795 |
| 1.16935 | 0.00037695 |
| 1.22007 | -7.305e-05 |
| 1.2713 | 0.00333695 |
| 1.32205 | 0.0139109 |
| 1.373 | 0.0251339 |
| 1.42372 | 0.036754 |
| 1.4747 | 0.0467869 |
| 1.52595 | 0.057124 |
| 1.57692 | 0.066631 |
| 1.6279 | 0.07558 |
| 1.67863 | 0.084216 |
| 1.72962 | 0.093227 |
| 1.7806 | 0.101444 |
| 1.83135 | 0.110011 |
| 1.88207 | 0.117717 |
| 1.93278 | 0.124988 |
| 1.98352 | 0.132655 |
| 2.03448 | 0.139804 |
| 2.08547 | 0.146442 |
| 2.1362 | 0.153384 |
| 2.18692 | 0.159663 |
| 2.23765 | 0.16553 |
| 2.28838 | 0.171664 |
| 2.33908 | 0.177577 |
| 2.3898 | 0.182422 |
| 2.44105 | 0.187175 |
| 2.49205 | 0.19083 |
| 2.54278 | 0.196307 |
| 2.59377 | 0.200114 |
| 2.64397 | 0.205081 |
| 2.6952 | 0.208431 |
| 2.74673 | 0.208698 |
| 2.79772 | 0.2168 |
| 2.84843 | 0.219638 |
| 2.89942 | 0.223613 |
| 2.9504 | 0.221584 |
| 3.00113 | 0.224742 |
| 3.05183 | 0.227229 |
| 3.10253 | 0.230594 |
| 3.15352 | 0.232219 |
| 3.20423 | 0.239185 |
| 3.25523 | 0.241977 |
| 3.30622 | 0.239948 |
| 3.35693 | 0.246333 |
| 3.40793 | 0.244792 |
| 3.45867 | 0.248042 |
| 3.50937 | 0.253032 |
| 3.55985 | 0.250224 |
| 3.61083 | 0.252963 |
| 3.66182 | 0.257991 |
| 3.71257 | 0.255862 |
| 3.76355 | 0.258319 |
| 3.81427 | 0.259113 |
| 3.86502 | 0.261348 |
| 3.91572 | 0.262965 |
| 3.96672 | 0.264545 |
| 4.01768 | 0.266292 |
| 4.0684 | 0.268131 |
| 4.11912 | 0.269443 |
| 4.1701 | 0.270046 |
| 4.22108 | 0.271487 |
| 4.27208 | 0.273509 |
| 4.3228 | 0.27486 |
| 4.37352 | 0.275974 |
| 4.42452 | 0.277423 |
| 4.47552 | 0.278476 |
| 4.52622 | 0.280055 |
| 4.57722 | 0.28049 |
| 4.6282 | 0.282497 |
| 4.67893 | 0.283511 |
| 4.72963 | 0.285114 |
| 4.78062 | 0.285808 |
| 4.83133 | 0.287242 |
| 4.88205 | 0.288234 |
| 4.93303 | 0.289088 |
| 4.98375 | 0.290057 |
| 5.03473 | 0.291332 |
| 5.08548 | 0.291652 |
| 5.1362 | 0.293735 |
| 5.18693 | 0.294521 |
| 5.23793 | 0.295421 |
| 5.28918 | 0.296367 |
| 5.34015 | 0.296924 |
| 5.39088 | 0.297725 |
| 5.44187 | 0.298801 |
| 5.49285 | 0.300349 |
| 5.54355 | 0.299793 |
| 5.59427 | 0.300487 |
| 5.64525 | 0.30096 |
| 5.69622 | 0.301646 |
| 5.74642 | 0.301181 |
| 5.79768 | 0.30247 |
| 5.84842 | 0.302219 |
| 5.89945 | 0.303767 |
| 5.95043 | 0.303935 |
| 6.00115 | 0.304462 |
| 6.05213 | 0.304996 |
| 6.10312 | 0.304965 |
| 6.1541 | 0.305431 |
| 6.20482 | 0.305911 |
| 6.25582 | 0.306796 |
| 6.3068 | 0.306827 |
| 6.3578 | 0.307819 |
| 6.40878 | 0.307201 |
| 6.45923 | 0.307964 |
| 6.51077 | 0.308826 |
| 6.56175 | 0.309299 |
| 6.61273 | 0.308856 |
| 6.6637 | 0.309894 |
| 6.71468 | 0.309146 |
| 6.76567 | 0.311389 |
| 6.81613 | 0.311137 |
| 6.86763 | 0.311794 |
| 6.9186 | 0.311153 |
| 6.9696 | 0.312022 |
| 7.02058 | 0.311878 |
| 7.07157 | 0.311885 |
| 7.12232 | 0.312999 |
| 7.17328 | 0.312427 |
| 7.22428 | 0.312579 |
| 7.27503 | 0.313091 |
| 7.32573 | 0.313174 |
| 7.37643 | 0.313442 |
| 7.4269 | 0.313373 |
| 7.4784 | 0.313228 |
| 7.5294 | 0.314639 |
| 7.58037 | 0.312854 |
| 7.63135 | 0.313815 |
| 7.68232 | 0.31377 |
| 7.73307 | 0.31496 |
| 7.78403 | 0.314433 |
| 7.83502 | 0.314304 |
| 7.886 | 0.315334 |
| 7.937 | 0.314357 |
| 7.98797 | 0.315784 |
| 8.0387 | 0.315959 |
| 8.08945 | 0.316592 |
| 8.1407 | 0.315112 |
| 8.19143 | 0.31525 |
| 8.24243 | 0.315303 |
| 8.29315 | 0.316142 |
| 8.34387 | 0.315273 |
| 8.39487 | 0.316562 |
| 8.44585 | 0.315845 |
| 8.49657 | 0.315593 |
| 8.54757 | 0.316051 |
| 8.59828 | 0.315883 |
| 8.64928 | 0.315463 |
| 8.70027 | 0.315372 |
| 8.75098 | 0.316119 |
| 8.8017 | 0.316669 |
| 8.85268 | 0.316707 |
| 8.90367 | 0.315578 |
| 8.95467 | 0.315913 |
| 9.00563 | 0.31602 |
| 9.05663 | 0.317188 |
| 9.1076 | 0.31673 |
| 9.15833 | 0.315379 |
| 9.20907 | 0.315738 |
| 9.26005 | 0.316371 |
| 9.31102 | 0.315822 |
| 9.36177 | 0.317088 |
| 9.41248 | 0.317256 |
| 9.46322 | 0.315471 |
| 9.5142 | 0.31721 |
| 9.56517 | 0.315631 |
| 9.61617 | 0.315883 |
| 9.66715 | 0.317142 |
| 9.71787 | 0.317119 |
| 9.76858 | 0.315898 |
| 9.81957 | 0.316249 |
| 9.8703 | 0.31644 |
| 9.921 | 0.317416 |
| 9.972 | 0.316913 |
| 10.0227 | 0.317279 |
| 10.0737 | 0.317401 |
| 10.1244 | 0.316631 |

| Time [min] | 85 nM |
| --- | --- |
| 0 | 0.00072735 |
| 0.050716 | 0.00017835 |
| 0.101717 | 0.00046835 |
| 0.15245 | 7.135e-05 |
| 0.20345 | 0.00075835 |
| 0.254166 | -0.00137065 |
| 0.304883 | 0.00076635 |
| 0.355617 | 0.00165135 |
| 0.406066 | -0.00083665 |
| 0.4576 | 7.935e-05 |
| 0.5086 | 1.835e-05 |
| 0.559333 | -0.00044665 |
| 0.610316 | 0.00045335 |
| 0.661016 | -1.965e-05 |
| 0.712033 | 0.00052935 |
| 0.762766 | -0.00013465 |
| 0.813766 | 0.00076635 |
| 0.864516 | -0.00038565 |
| 0.915516 | -0.00062265 |
| 0.966233 | -0.00052365 |
| 1.01697 | -0.00046965 |
| 1.0677 | -0.00037865 |
| 1.11843 | -0.00024065 |
| 1.16918 | 1.835e-05 |
| 1.21992 | -0.00220965 |
| 1.27067 | -0.00194965 |
| 1.3214 | 0.00132335 |
| 1.3724 | 0.00563335 |
| 1.42312 | 0.0115003 |
| 1.47387 | 0.0181683 |
| 1.5246 | 0.0250803 |
| 1.57533 | 0.0338703 |
| 1.62608 | 0.0393023 |
| 1.67678 | 0.0470533 |
| 1.72725 | 0.0548733 |
| 1.7785 | 0.0590693 |
| 1.82948 | 0.0656693 |
| 1.88048 | 0.0703843 |
| 1.93147 | 0.0758853 |
| 1.98218 | 0.0792803 |
| 2.0329 | 0.0852313 |
| 2.0839 | 0.0893123 |
| 2.13488 | 0.0939663 |
| 2.18587 | 0.0994213 |
| 2.23687 | 0.103335 |
| 2.28785 | 0.10792 |
| 2.33885 | 0.112582 |
| 2.38982 | 0.115886 |
| 2.44082 | 0.119616 |
| 2.4918 | 0.125117 |
| 2.54283 | 0.130694 |
| 2.59383 | 0.134791 |
| 2.64483 | 0.139407 |
| 2.69582 | 0.142031 |
| 2.74685 | 0.148661 |
| 2.79753 | 0.153147 |
| 2.84797 | 0.157107 |
| 2.89925 | 0.159404 |
| 2.95002 | 0.163127 |
| 3.001 | 0.169207 |
| 3.05173 | 0.173983 |
| 3.10245 | 0.174945 |
| 3.15345 | 0.181315 |
| 3.20445 | 0.18513 |
| 3.25543 | 0.18912 |
| 3.30615 | 0.190379 |
| 3.3572 | 0.194217 |
| 3.40797 | 0.198062 |
| 3.45842 | 0.203288 |
| 3.5094 | 0.205279 |
| 3.5607 | 0.210727 |
| 3.61167 | 0.211665 |
| 3.66265 | 0.214831 |
| 3.71365 | 0.221728 |
| 3.76437 | 0.22127 |
| 3.81508 | 0.225428 |
| 3.86555 | 0.226756 |
| 3.91653 | 0.23202 |
| 3.9678 | 0.232287 |
| 4.0188 | 0.238032 |
| 4.06952 | 0.237735 |
| 4.1205 | 0.239581 |
| 4.17148 | 0.244868 |
| 4.22247 | 0.244433 |
| 4.27345 | 0.247042 |
| 4.32445 | 0.249461 |
| 4.37545 | 0.253558 |
| 4.42643 | 0.254535 |
| 4.47715 | 0.256213 |
| 4.52788 | 0.257907 |
| 4.57865 | 0.261897 |
| 4.62962 | 0.262934 |
| 4.68035 | 0.264224 |
| 4.73132 | 0.265612 |
| 4.78205 | 0.26797 |
| 4.83303 | 0.26871 |
| 4.88378 | 0.271975 |
| 4.93503 | 0.273776 |
| 4.98552 | 0.2746 |
| 5.0365 | 0.279513 |
| 5.08725 | 0.276805 |
| 5.13797 | 0.279269 |
| 5.18895 | 0.280574 |
| 5.23968 | 0.280619 |
| 5.2904 | 0.284373 |
| 5.34138 | 0.283473 |
| 5.39238 | 0.286212 |
| 5.4431 | 0.287799 |
| 5.49412 | 0.288623 |
| 5.54485 | 0.289653 |
| 5.59557 | 0.289805 |
| 5.64628 | 0.29198 |
| 5.69728 | 0.29169 |
| 5.74827 | 0.293765 |
| 5.79927 | 0.293261 |
| 5.85023 | 0.295451 |
| 5.90123 | 0.295779 |
| 5.95222 | 0.296618 |
| 6.00293 | 0.295794 |
| 6.05367 | 0.297618 |
| 6.10465 | 0.29774 |
| 6.15565 | 0.299754 |
| 6.20638 | 0.299144 |
| 6.25737 | 0.300257 |
| 6.3081 | 0.301707 |
| 6.35908 | 0.301165 |
| 6.40953 | 0.302188 |
| 6.46107 | 0.302295 |
| 6.51178 | 0.303775 |
| 6.56275 | 0.302226 |
| 6.61348 | 0.303866 |
| 6.66422 | 0.303035 |
| 6.71497 | 0.303981 |
| 6.76593 | 0.304126 |
| 6.81668 | 0.304423 |
| 6.8674 | 0.304934 |
| 6.91813 | 0.305285 |
| 6.9691 | 0.304942 |
| 7.02012 | 0.305224 |
| 7.07108 | 0.305651 |
| 7.12208 | 0.306254 |
| 7.17282 | 0.305514 |
| 7.22353 | 0.307727 |
| 7.27427 | 0.307185 |
| 7.32498 | 0.30717 |
| 7.3757 | 0.306269 |
| 7.42642 | 0.30749 |
| 7.4774 | 0.308047 |
| 7.52813 | 0.307887 |
| 7.57885 | 0.307475 |
| 7.62983 | 0.308078 |
| 7.68082 | 0.309146 |
| 7.73183 | 0.309115 |
| 7.78253 | 0.308932 |
| 7.8333 | 0.308451 |
| 7.88402 | 0.309253 |
| 7.93475 | 0.309008 |
| 7.98572 | 0.308337 |
| 8.03645 | 0.30878 |
| 8.08717 | 0.312075 |
| 8.13792 | 0.309291 |
| 8.18865 | 0.308986 |
| 8.23938 | 0.308596 |
| 8.2901 | 0.310244 |
| 8.34112 | 0.309634 |
| 8.39183 | 0.309802 |
| 8.44283 | 0.30897 |
| 8.4938 | 0.309497 |
| 8.5448 | 0.309672 |
| 8.59552 | 0.308093 |
| 8.64623 | 0.308749 |
| 8.69722 | 0.310725 |
| 8.74822 | 0.308558 |
| 8.79892 | 0.30942 |
| 8.84965 | 0.310847 |
| 8.90063 | 0.310252 |
| 8.95137 | 0.309726 |
| 9.00205 | 0.309924 |
| 9.05252 | 0.310672 |
| 9.10403 | 0.309596 |
| 9.15503 | 0.310382 |
| 9.20575 | 0.308726 |
| 9.25648 | 0.312724 |
| 9.30722 | 0.310122 |
| 9.35795 | 0.313472 |
| 9.40893 | 0.310412 |
| 9.45968 | 0.308642 |
| 9.51042 | 0.310633 |
| 9.56115 | 0.309809 |
| 9.6119 | 0.311396 |
| 9.66288 | 0.308795 |
| 9.7136 | 0.30881 |
| 9.76458 | 0.31016 |
| 9.81557 | 0.310427 |
| 9.86655 | 0.308078 |
| 9.9173 | 0.309886 |
| 9.96802 | 0.30865 |
| 10.019 | 0.311389 |
| 10.07 | 0.309665 |
| 10.121 | 0.309863 |

| Time [min] | 50 nM |
| --- | --- |
| 0 | -0.000208476 |
| 0.050683 | 0.000264524 |
| 0.10265 | 0.000157524 |
| 0.15255 | -0.000284476 |
| 0.203266 | -0.000910476 |
| 0.25425 | 0.000447524 |
| 0.304983 | -4.84762e-05 |
| 0.355966 | -0.000818476 |
| 0.4067 | -0.000170476 |
| 0.460599 | 0.000172524 |
| 0.51135 | -0.000650476 |
| 0.562333 | -0.000322476 |
| 0.613333 | 8.15238e-05 |
| 0.66405 | 0.000226524 |
| 0.714767 | -0.000239476 |
| 0.765749 | -0.000132476 |
| 0.816483 | -0.000368476 |
| 0.867199 | -0.000681476 |
| 0.917933 | 0.000241524 |
| 0.968916 | -0.000307476 |
| 1.0199 | -0.000696476 |
| 1.0709 | -0.000780476 |
| 1.12188 | -0.000902476 |
| 1.17257 | -0.000856476 |
| 1.22328 | -0.000612476 |
| 1.274 | 0.00252252 |
| 1.32475 | 0.00893152 |
| 1.37545 | 0.0137995 |
| 1.42645 | 0.0191325 |
| 1.47718 | 0.0249155 |
| 1.5279 | 0.0304845 |
| 1.57888 | 0.0354975 |
| 1.62987 | 0.0401665 |
| 1.68058 | 0.0450715 |
| 1.73103 | 0.0494665 |
| 1.782 | 0.0543725 |
| 1.83273 | 0.0590945 |
| 1.88372 | 0.0636955 |
| 1.93472 | 0.0678765 |
| 1.98542 | 0.0726675 |
| 2.0364 | 0.0770235 |
| 2.08737 | 0.0816855 |
| 2.13837 | 0.0849125 |
| 2.18913 | 0.0897265 |
| 2.2399 | 0.0941285 |
| 2.29088 | 0.0972645 |
| 2.34162 | 0.101011 |
| 2.39233 | 0.103537 |
| 2.44307 | 0.108961 |
| 2.49405 | 0.112585 |
| 2.54503 | 0.116743 |
| 2.59577 | 0.11759 |
| 2.64648 | 0.121076 |
| 2.69747 | 0.126745 |
| 2.74818 | 0.129873 |
| 2.79915 | 0.131613 |
| 2.84988 | 0.136755 |
| 2.9006 | 0.136991 |
| 2.95132 | 0.142416 |
| 3.00232 | 0.14282 |
| 3.05332 | 0.148306 |
| 3.10428 | 0.148672 |
| 3.15502 | 0.151098 |
| 3.206 | 0.15666 |
| 3.25673 | 0.156073 |
| 3.30745 | 0.158995 |
| 3.35815 | 0.161787 |
| 3.40887 | 0.163938 |
| 3.45958 | 0.165861 |
| 3.51058 | 0.171606 |
| 3.56133 | 0.17182 |
| 3.6123 | 0.175222 |
| 3.6633 | 0.174879 |
| 3.71428 | 0.176924 |
| 3.76528 | 0.178724 |
| 3.81598 | 0.181662 |
| 3.86673 | 0.182989 |
| 3.91743 | 0.185263 |
| 3.96813 | 0.185972 |
| 4.01857 | 0.188284 |
| 4.06982 | 0.190008 |
| 4.12053 | 0.191374 |
| 4.17072 | 0.193014 |
| 4.22222 | 0.194509 |
| 4.27322 | 0.196569 |
| 4.32418 | 0.198492 |
| 4.37518 | 0.200155 |
| 4.42615 | 0.201597 |
| 4.47713 | 0.202772 |
| 4.52808 | 0.203726 |
| 4.5788 | 0.205969 |
| 4.62953 | 0.206999 |
| 4.68053 | 0.207731 |
| 4.7315 | 0.209387 |
| 4.78248 | 0.21076 |
| 4.83347 | 0.211775 |
| 4.88447 | 0.212858 |
| 4.9352 | 0.213789 |
| 4.98618 | 0.215712 |
| 5.0369 | 0.217054 |
| 5.08788 | 0.216871 |
| 5.13887 | 0.217619 |
| 5.18962 | 0.219664 |
| 5.24032 | 0.219938 |
| 5.29105 | 0.22077 |
| 5.34203 | 0.221075 |
| 5.393 | 0.221853 |
| 5.44373 | 0.223143 |
| 5.49447 | 0.224272 |
| 5.54545 | 0.223631 |
| 5.59565 | 0.224699 |
| 5.64717 | 0.225202 |
| 5.69788 | 0.225332 |
| 5.74888 | 0.225874 |
| 5.7996 | 0.226835 |
| 5.8503 | 0.226362 |
| 5.90103 | 0.226583 |
| 5.95175 | 0.227468 |
| 6.00247 | 0.228361 |
| 6.0532 | 0.227476 |
| 6.1042 | 0.227331 |
| 6.15492 | 0.228376 |
| 6.2059 | 0.228697 |
| 6.25662 | 0.228323 |
| 6.30762 | 0.228216 |
| 6.35858 | 0.228109 |
| 6.4093 | 0.22843 |
| 6.46002 | 0.228399 |
| 6.511 | 0.229818 |
| 6.56172 | 0.230528 |
| 6.61272 | 0.229093 |
| 6.66343 | 0.229345 |
| 6.71415 | 0.228681 |
| 6.76488 | 0.22965 |
| 6.81587 | 0.230612 |
| 6.8666 | 0.231344 |
| 6.91732 | 0.231245 |
| 6.96802 | 0.230604 |
| 7.019 | 0.228872 |
| 7.06997 | 0.230352 |
| 7.12072 | 0.229757 |
| 7.17143 | 0.230764 |
| 7.22215 | 0.230078 |
| 7.27288 | 0.231153 |
| 7.3236 | 0.229589 |
| 7.37432 | 0.230955 |
| 7.4253 | 0.230879 |
| 7.47628 | 0.230085 |
| 7.52727 | 0.230604 |
| 7.57823 | 0.229132 |
| 7.62922 | 0.229406 |
| 7.68018 | 0.230642 |
| 7.73117 | 0.230772 |
| 7.78188 | 0.23078 |
| 7.83287 | 0.229597 |
| 7.8836 | 0.230947 |
| 7.93433 | 0.230925 |
| 7.98532 | 0.23097 |
| 8.03602 | 0.230322 |
| 8.087 | 0.230528 |
| 8.13798 | 0.23004 |
| 8.18895 | 0.230207 |
| 8.23993 | 0.22991 |
| 8.29065 | 0.23007 |
| 8.3419 | 0.23049 |
| 8.39288 | 0.229307 |
| 8.4436 | 0.230017 |
| 8.49458 | 0.229605 |
| 8.54557 | 0.229322 |
| 8.59655 | 0.229345 |
| 8.64753 | 0.229635 |
| 8.6985 | 0.229536 |
| 8.74948 | 0.229414 |
| 8.80047 | 0.229696 |
| 8.85147 | 0.229437 |
| 8.90218 | 0.229551 |
| 8.95317 | 0.229154 |
| 9.0044 | 0.229528 |
| 9.05513 | 0.23004 |
| 9.10583 | 0.229673 |
| 9.15655 | 0.230017 |
| 9.20753 | 0.228842 |
| 9.25827 | 0.22862 |
| 9.30898 | 0.229193 |
| 9.35972 | 0.228697 |
| 9.41068 | 0.228582 |
| 9.46167 | 0.229078 |
| 9.51267 | 0.22901 |
| 9.56365 | 0.228865 |
| 9.61438 | 0.228552 |
| 9.6651 | 0.228529 |
| 9.7161 | 0.228025 |
| 9.76735 | 0.227812 |
| 9.81808 | 0.229101 |
| 9.86908 | 0.228689 |
| 9.92007 | 0.228018 |
| 9.97078 | 0.228712 |
| 10.0215 | 0.228712 |
| 10.0722 | 0.228842 |
| 10.1232 | 0.228086 |

| Time [min] | 25 nM |
| --- | --- |
| 0 | -0.00025695 |
| 0.050716 | 0.00069605 |
| 0.101717 | 0.00046805 |
| 0.15245 | -0.00046295 |
| 0.20345 | 0.00033005 |
| 0.254166 | 0.00024605 |
| 0.304883 | -0.00070695 |
| 0.355617 | 0.00036105 |
| 0.406066 | -0.00134095 |
| 0.4576 | -0.00121795 |
| 0.5086 | 0.00036105 |
| 0.559333 | 0.00018505 |
| 0.610316 | 0.00044505 |
| 0.661016 | 0.00026205 |
| 0.712033 | 0.00033005 |
| 0.762766 | 0.00039905 |
| 0.813766 | -0.00060795 |
| 0.864516 | 0.00030705 |
| 0.915516 | -0.00134095 |
| 0.966233 | 0.00033805 |
| 1.01697 | -0.00134095 |
| 1.0677 | -0.00051695 |
| 1.11843 | 0.00099405 |
| 1.16918 | 0.00062005 |
| 1.21992 | -9.695e-05 |
| 1.27067 | -0.00024995 |
| 1.3214 | 0.00054405 |
| 1.3724 | 0.00342805 |
| 1.42312 | 0.00642605 |
| 1.47387 | 0.00899705 |
| 1.5246 | 0.0131021 |
| 1.57533 | 0.0139791 |
| 1.62608 | 0.0167641 |
| 1.67678 | 0.0196711 |
| 1.72725 | 0.0222271 |
| 1.7785 | 0.0230431 |
| 1.82948 | 0.0249881 |
| 1.88048 | 0.0275601 |
| 1.93147 | 0.0289941 |
| 1.98218 | 0.0325111 |
| 2.0329 | 0.0342891 |
| 2.0839 | 0.0354871 |
| 2.13488 | 0.0376001 |
| 2.18587 | 0.0390041 |
| 2.23687 | 0.0410411 |
| 2.28785 | 0.0438641 |
| 2.33885 | 0.0446951 |
| 2.38982 | 0.0464501 |
| 2.44082 | 0.0486401 |
| 2.4918 | 0.0505161 |
| 2.54283 | 0.0512261 |
| 2.59383 | 0.0524311 |
| 2.64483 | 0.0552471 |
| 2.69582 | 0.0563761 |
| 2.74685 | 0.0584281 |
| 2.79753 | 0.0597401 |
| 2.84797 | 0.0610911 |
| 2.89925 | 0.0629291 |
| 2.95002 | 0.0647911 |
| 3.001 | 0.0667901 |
| 3.05173 | 0.0669881 |
| 3.10245 | 0.0703221 |
| 3.15345 | 0.0709561 |
| 3.20445 | 0.0724131 |
| 3.25543 | 0.0746021 |
| 3.30615 | 0.0750141 |
| 3.3572 | 0.0773571 |
| 3.40797 | 0.0788751 |
| 3.45842 | 0.0794621 |
| 3.5094 | 0.0817051 |
| 3.5607 | 0.0835291 |
| 3.61167 | 0.0839261 |
| 3.66265 | 0.0855581 |
| 3.71365 | 0.0869701 |
| 3.76437 | 0.0882361 |
| 3.81508 | 0.0897851 |
| 3.86555 | 0.0928521 |
| 3.91653 | 0.0922641 |
| 3.9678 | 0.0935921 |
| 4.0188 | 0.0951101 |
| 4.06952 | 0.0971851 |
| 4.1205 | 0.0983601 |
| 4.17148 | 0.0991161 |
| 4.22247 | 0.101496 |
| 4.27345 | 0.103052 |
| 4.32445 | 0.10406 |
| 4.37545 | 0.105151 |
| 4.42643 | 0.106898 |
| 4.47715 | 0.107783 |
| 4.52788 | 0.109591 |
| 4.57865 | 0.109957 |
| 4.62962 | 0.111971 |
| 4.68035 | 0.111086 |
| 4.73132 | 0.111773 |
| 4.78205 | 0.113581 |
| 4.83303 | 0.11674 |
| 4.88378 | 0.117968 |
| 4.93503 | 0.119387 |
| 4.98552 | 0.117747 |
| 5.0365 | 0.12163 |
| 5.08725 | 0.122011 |
| 5.13797 | 0.1215 |
| 5.18895 | 0.123354 |
| 5.23968 | 0.123827 |
| 5.2904 | 0.127375 |
| 5.34138 | 0.125483 |
| 5.39238 | 0.129412 |
| 5.4431 | 0.130297 |
| 5.49412 | 0.129061 |
| 5.54485 | 0.131937 |
| 5.59557 | 0.130228 |
| 5.64628 | 0.130671 |
| 5.69728 | 0.133135 |
| 5.74827 | 0.134425 |
| 5.79927 | 0.135218 |
| 5.85023 | 0.134692 |
| 5.90123 | 0.13341 |
| 5.95222 | 0.136195 |
| 6.00293 | 0.134814 |
| 6.05367 | 0.135485 |
| 6.10465 | 0.136111 |
| 6.15565 | 0.139315 |
| 6.20638 | 0.137354 |
| 6.25737 | 0.138514 |
| 6.3081 | 0.139208 |
| 6.35908 | 0.139757 |
| 6.40953 | 0.140848 |
| 6.46107 | 0.14332 |
| 6.51178 | 0.142557 |
| 6.56275 | 0.142802 |
| 6.61348 | 0.143244 |
| 6.66422 | 0.145884 |
| 6.71497 | 0.143977 |
| 6.76593 | 0.144251 |
| 6.81668 | 0.147509 |
| 6.8674 | 0.145281 |
| 6.91813 | 0.146174 |
| 6.9691 | 0.146052 |
| 7.02012 | 0.147364 |
| 7.07108 | 0.147692 |
| 7.12208 | 0.147921 |
| 7.17282 | 0.148012 |
| 7.22353 | 0.148653 |
| 7.27427 | 0.148348 |
| 7.32498 | 0.148615 |
| 7.3757 | 0.149462 |
| 7.42642 | 0.149515 |
| 7.4774 | 0.150202 |
| 7.52813 | 0.150378 |
| 7.57885 | 0.15082 |
| 7.62983 | 0.150973 |
| 7.68082 | 0.15127 |
| 7.73183 | 0.151491 |
| 7.78253 | 0.151736 |
| 7.8333 | 0.15214 |
| 7.88402 | 0.152293 |
| 7.93475 | 0.152544 |
| 7.98572 | 0.155474 |
| 8.03645 | 0.153742 |
| 8.08717 | 0.153765 |
| 8.13792 | 0.155924 |
| 8.18865 | 0.154291 |
| 8.23938 | 0.154642 |
| 8.2901 | 0.155909 |
| 8.34112 | 0.15478 |
| 8.39183 | 0.155093 |
| 8.44283 | 0.155588 |
| 8.4938 | 0.157526 |
| 8.5448 | 0.157641 |
| 8.59552 | 0.156054 |
| 8.64623 | 0.156069 |
| 8.69722 | 0.156489 |
| 8.74822 | 0.158434 |
| 8.79892 | 0.156908 |
| 8.84965 | 0.156924 |
| 8.90063 | 0.158915 |
| 8.95137 | 0.159289 |
| 9.00205 | 0.157412 |
| 9.05252 | 0.159884 |
| 9.10403 | 0.15774 |
| 9.15503 | 0.157824 |
| 9.20575 | 0.158144 |
| 9.25648 | 0.158915 |
| 9.30722 | 0.15922 |
| 9.35795 | 0.158732 |
| 9.40893 | 0.161288 |
| 9.45968 | 0.159533 |
| 9.51042 | 0.159724 |
| 9.56115 | 0.162234 |
| 9.6119 | 0.162348 |
| 9.66288 | 0.1626 |
| 9.7136 | 0.162402 |
| 9.76458 | 0.161646 |
| 9.81557 | 0.161043 |
| 9.86655 | 0.161509 |
| 9.9173 | 0.162028 |
| 9.96802 | 0.162348 |
| 10.019 | 0.162234 |
| 10.07 | 0.162493 |
| 10.121 | 0.162928 |

| Time [min] | 25 nM |
| --- | --- |
| -1.01733 | -0.000349952 |
| -0.966617 | 0.000344048 |
| -0.915901 | -0.000112952 |
| -0.864933 | 0.000733048 |
| -0.813949 | 0.000299048 |
| -0.762968 | -0.000227952 |
| -0.711967 | -0.000341952 |
| -0.661233 | -0.000349952 |
| -0.61025 | 0.000184048 |
| -0.559534 | 0.000291048 |
| -0.508533 | -0.000593952 |
| -0.45755 | 0.000146048 |
| -0.406583 | -0.000471952 |
| -0.355833 | 9.30476e-05 |
| -0.305101 | -0.000555952 |
| -0.2541 | 2.40476e-05 |
| -0.203117 | -0.00123495 |
| -0.152416 | 0.000215048 |
| -0.101699 | 0.000573048 |
| -0.050701 | 0.000283048 |
| 0 | 0.00105405 |
| 0.050999 | -0.00108995 |
| 0.101733 | 0.000169048 |
| 0.152701 | -0.000242952 |
| 0.2037 | -0.000501952 |
| 0.254666 | -0.000761952 |
| 0.305384 | 9.04762e-06 |
| 0.35635 | -0.00147095 |
| 0.407084 | -5.99524e-05 |
| 0.458067 | -0.00121995 |
| 0.509066 | -0.000143952 |
| 0.56005 | 0.000512048 |
| 0.610783 | -0.000822952 |
| 0.661517 | -0.00134195 |
| 0.712517 | 0.000322048 |
| 0.763484 | -7.49524e-05 |
| 0.814234 | -0.000151952 |
| 0.864933 | -0.000662952 |
| 0.915667 | 0.000100048 |
| 0.966401 | -0.000776952 |
| 1.01738 | -0.000410952 |
| 1.06837 | -0.000974952 |
| 1.11937 | -0.000501952 |
| 1.17033 | -0.000425952 |
| 1.22133 | -0.000608952 |
| 1.27205 | 0.00205305 |
| 1.32305 | 0.00496805 |
| 1.37403 | 0.00911805 |
| 1.42477 | 0.011842 |
| 1.47547 | 0.015802 |
| 1.5262 | 0.018144 |
| 1.57692 | 0.022218 |
| 1.6279 | 0.025193 |
| 1.67888 | 0.027757 |
| 1.72988 | 0.031831 |
| 1.7806 | 0.035241 |
| 1.83133 | 0.037858 |
| 1.88205 | 0.04075 |
| 1.93307 | 0.044122 |
| 1.98402 | 0.047525 |
| 2.03502 | 0.050477 |
| 2.08573 | 0.05288 |
| 2.1367 | 0.055879 |
| 2.1874 | 0.05848 |
| 2.2384 | 0.061982 |
| 2.2891 | 0.064752 |
| 2.3401 | 0.067872 |
| 2.39108 | 0.070916 |
| 2.4418 | 0.073236 |
| 2.49278 | 0.075791 |
| 2.5435 | 0.078851 |
| 2.59422 | 0.077569 |
| 2.64493 | 0.084237 |
| 2.6959 | 0.086862 |
| 2.74687 | 0.089906 |
| 2.79758 | 0.092385 |
| 2.84857 | 0.09382 |
| 2.89957 | 0.09337 |
| 2.9503 | 0.095536 |
| 3.00103 | 0.098397 |
| 3.05175 | 0.104447 |
| 3.10273 | 0.103288 |
| 3.15372 | 0.105577 |
| 3.20442 | 0.110688 |
| 3.25542 | 0.110482 |
| 3.30613 | 0.114816 |
| 3.35713 | 0.115274 |
| 3.40812 | 0.11786 |
| 3.45883 | 0.120179 |
| 3.50982 | 0.122567 |
| 3.56055 | 0.124452 |
| 3.61127 | 0.126886 |
| 3.66198 | 0.129655 |
| 3.71243 | 0.131913 |
| 3.7637 | 0.133302 |
| 3.81468 | 0.135949 |
| 3.8654 | 0.137803 |
| 3.91612 | 0.139535 |
| 3.96683 | 0.141786 |
| 4.01755 | 0.143685 |
| 4.06852 | 0.145753 |
| 4.11925 | 0.147363 |
| 4.17023 | 0.150048 |
| 4.22098 | 0.152543 |
| 4.272 | 0.15426 |
| 4.32298 | 0.156213 |
| 4.37373 | 0.158876 |
| 4.42445 | 0.159036 |
| 4.47543 | 0.160722 |
| 4.52642 | 0.163034 |
| 4.57742 | 0.164117 |
| 4.6284 | 0.166078 |
| 4.67938 | 0.167848 |
| 4.73035 | 0.170312 |
| 4.78135 | 0.171479 |
| 4.83233 | 0.173776 |
| 4.88333 | 0.174295 |
| 4.9343 | 0.176339 |
| 4.9853 | 0.177682 |
| 5.03628 | 0.179467 |
| 5.08727 | 0.181405 |
| 5.13823 | 0.181832 |
| 5.18922 | 0.183007 |
| 5.2402 | 0.184671 |
| 5.2912 | 0.185838 |
| 5.3419 | 0.187844 |
| 5.39288 | 0.189736 |
| 5.44362 | 0.189843 |
| 5.49462 | 0.192063 |
| 5.54533 | 0.19201 |
| 5.59607 | 0.193338 |
| 5.64705 | 0.194566 |
| 5.69803 | 0.194101 |
| 5.74877 | 0.194421 |
| 5.79978 | 0.194963 |
| 5.85077 | 0.197328 |
| 5.9015 | 0.197312 |
| 5.95248 | 0.198831 |
| 6.0032 | 0.198381 |
| 6.05418 | 0.200196 |
| 6.1049 | 0.199792 |
| 6.15587 | 0.200494 |
| 6.20687 | 0.201005 |
| 6.2576 | 0.200181 |
| 6.30858 | 0.201852 |
| 6.35957 | 0.201478 |
| 6.41055 | 0.20215 |
| 6.46155 | 0.202699 |
| 6.51228 | 0.203935 |
| 6.563 | 0.203736 |
| 6.614 | 0.203569 |
| 6.66497 | 0.205316 |
| 6.71597 | 0.204873 |
| 6.76668 | 0.20617 |
| 6.8174 | 0.206582 |
| 6.86838 | 0.20662 |
| 6.91937 | 0.206147 |
| 6.97008 | 0.205293 |
| 7.02081 | 0.206422 |
| 7.07178 | 0.205819 |
| 7.12251 | 0.206849 |
| 7.17322 | 0.206506 |
| 7.2242 | 0.207322 |
| 7.27545 | 0.208367 |
| 7.32645 | 0.207002 |
| 7.37743 | 0.207612 |
| 7.42815 | 0.207299 |
| 7.47885 | 0.208161 |
| 7.52983 | 0.207605 |
| 7.58083 | 0.208764 |
| 7.63155 | 0.208642 |
| 7.68253 | 0.209184 |
| 7.73353 | 0.209138 |
| 7.78428 | 0.209619 |
| 7.83525 | 0.209726 |
| 7.88623 | 0.209855 |
| 7.93695 | 0.210122 |
| 7.98767 | 0.210458 |
| 8.0384 | 0.210435 |
| 8.08913 | 0.21074 |
| 8.13985 | 0.21074 |
| 8.19083 | 0.210679 |
| 8.24182 | 0.210343 |
| 8.29278 | 0.210984 |
| 8.34378 | 0.210183 |
| 8.39477 | 0.211518 |
| 8.44548 | 0.211076 |
| 8.49647 | 0.210611 |
| 8.5472 | 0.209954 |
| 8.59819 | 0.21145 |
| 8.64915 | 0.211434 |
| 8.69989 | 0.211984 |
| 8.75062 | 0.211724 |
| 8.80159 | 0.210694 |
| 8.85258 | 0.211846 |
| 8.90328 | 0.212114 |
| 8.954 | 0.211251 |
| 9.00472 | 0.211671 |
| 9.05572 | 0.212266 |
| 9.10643 | 0.21148 |

| Time [min] | 25 nM |
| --- | --- |
| 0 | -0.00081 |
| 0.050983 | -0.000505 |
| 0.101967 | 0.000182 |
| 0.152683 | 0.000174 |
| 0.203684 | 0.000426 |
| 0.254667 | 0.000335 |
| 0.305384 | -1e-06 |
| 0.356367 | 0.000434 |
| 0.4071 | -0.000383 |
| 0.4581 | 0.000502 |
| 0.509084 | -0.000627 |
| 0.560067 | -0.000817 |
| 0.6108 | -0.001138 |
| 0.661783 | -0.00023 |
| 0.712517 | -0.000856 |
| 0.762684 | -0.0001 |
| 0.8142 | -0.000184 |
| 0.865167 | -0.000405 |
| 0.9159 | 0.000357 |
| 0.966884 | 0.000487 |
| 1.01787 | -4.7e-05 |
| 1.06887 | -0.000375 |
| 1.1196 | -0.000772 |
| 1.1698 | 0.000243 |
| 1.22107 | -4.7e-05 |
| 1.27177 | 0.000197 |
| 1.32275 | 0.002089 |
| 1.37372 | 0.00376 |
| 1.42447 | 0.004943 |
| 1.47522 | 0.004523 |
| 1.52567 | 0.007064 |
| 1.57693 | 0.007636 |
| 1.6279 | 0.008841 |
| 1.67887 | 0.009955 |
| 1.7296 | 0.011214 |
| 1.78088 | 0.011329 |
| 1.83188 | 0.012664 |
| 1.8829 | 0.013892 |
| 1.93365 | 0.012908 |
| 1.98467 | 0.015517 |
| 2.0354 | 0.016272 |
| 2.08637 | 0.016883 |
| 2.13708 | 0.017997 |
| 2.18807 | 0.018958 |
| 2.23905 | 0.019332 |
| 2.29007 | 0.02082 |
| 2.34052 | 0.021804 |
| 2.39208 | 0.022361 |
| 2.44307 | 0.023192 |
| 2.4938 | 0.024268 |
| 2.5448 | 0.024787 |
| 2.5958 | 0.02603 |
| 2.64678 | 0.026656 |
| 2.69777 | 0.027655 |
| 2.74875 | 0.028289 |
| 2.79972 | 0.029021 |
| 2.8507 | 0.030272 |
| 2.90147 | 0.031096 |
| 2.95218 | 0.031257 |
| 3.00317 | 0.032836 |
| 3.05395 | 0.033805 |
| 3.1047 | 0.034575 |
| 3.15542 | 0.035743 |
| 3.20613 | 0.036177 |
| 3.25712 | 0.036933 |
| 3.30783 | 0.037177 |
| 3.35883 | 0.038359 |
| 3.40983 | 0.039084 |
| 3.46082 | 0.04045 |
| 3.51178 | 0.040824 |
| 3.56278 | 0.041503 |
| 3.6135 | 0.042296 |
| 3.6645 | 0.042853 |
| 3.71548 | 0.044371 |
| 3.76647 | 0.044982 |
| 3.81745 | 0.045844 |
| 3.86817 | 0.046447 |
| 3.91915 | 0.046965 |
| 3.96988 | 0.048285 |
| 4.02085 | 0.048911 |
| 4.07157 | 0.049476 |
| 4.12255 | 0.049887 |
| 4.17353 | 0.051024 |
| 4.22452 | 0.051879 |
| 4.27552 | 0.052184 |
| 4.32625 | 0.053557 |
| 4.37697 | 0.053428 |
| 4.42795 | 0.054259 |
| 4.47893 | 0.055167 |
| 4.52965 | 0.055663 |
| 4.58037 | 0.056899 |
| 4.63082 | 0.057166 |
| 4.68237 | 0.057814 |
| 4.73307 | 0.058783 |
| 4.78432 | 0.058989 |
| 4.83528 | 0.059638 |
| 4.88632 | 0.060347 |
| 4.9373 | 0.060782 |
| 4.98828 | 0.060988 |
| 5.03928 | 0.062682 |
| 5.09028 | 0.063071 |
| 5.14098 | 0.063956 |
| 5.19198 | 0.064101 |
| 5.24242 | 0.064559 |
| 5.29367 | 0.064811 |
| 5.34438 | 0.065718 |
| 5.39512 | 0.066298 |
| 5.44582 | 0.06668 |
| 5.49682 | 0.067954 |
| 5.54753 | 0.068953 |
| 5.59883 | 0.068633 |
| 5.64955 | 0.070044 |
| 5.7003 | 0.070021 |
| 5.75102 | 0.070487 |
| 5.80175 | 0.07157 |
| 5.8522 | 0.071479 |
| 5.90347 | 0.071914 |
| 5.95445 | 0.072593 |
| 6.00533 | 0.072677 |
| 6.05632 | 0.073813 |
| 6.10732 | 0.074859 |
| 6.1583 | 0.073745 |
| 6.20927 | 0.075744 |
| 6.26002 | 0.076102 |
| 6.31128 | 0.076072 |
| 6.36227 | 0.076102 |
| 6.41328 | 0.07717 |
| 6.46427 | 0.07772 |
| 6.51525 | 0.078017 |
| 6.56623 | 0.077956 |
| 6.61725 | 0.079032 |
| 6.66822 | 0.079223 |
| 6.71922 | 0.078887 |
| 6.76993 | 0.079787 |
| 6.82065 | 0.080054 |
| 6.87135 | 0.080359 |
| 6.92262 | 0.08055 |
| 6.97333 | 0.08023 |
| 7.02407 | 0.081351 |
| 7.07503 | 0.081801 |
| 7.12603 | 0.081313 |
| 7.17727 | 0.081397 |
| 7.22828 | 0.082244 |
| 7.27925 | 0.082541 |
| 7.33 | 0.082862 |
| 7.38098 | 0.082434 |
| 7.43172 | 0.08335 |
| 7.48323 | 0.083388 |
| 7.53422 | 0.083083 |
| 7.5847 | 0.081664 |
| 7.63648 | 0.083403 |
| 7.68722 | 0.084205 |
| 7.73822 | 0.083953 |
| 7.78922 | 0.08377 |
| 7.83993 | 0.083686 |
| 7.8909 | 0.084227 |
| 7.94163 | 0.083907 |
| 7.99237 | 0.083907 |
| 8.04308 | 0.082778 |
| 8.09332 | 0.083915 |
| 8.14483 | 0.084052 |
| 8.19562 | 0.0838 |
| 8.2466 | 0.083945 |
| 8.29757 | 0.083327 |
| 8.34853 | 0.084182 |
| 8.39927 | 0.084037 |
| 8.45025 | 0.083228 |
| 8.50122 | 0.084487 |
| 8.5522 | 0.085067 |
| 8.60318 | 0.083663 |
| 8.65418 | 0.083495 |
| 8.70488 | 0.08425 |
| 8.75562 | 0.084494 |
| 8.8066 | 0.085486 |
| 8.85758 | 0.085135 |
| 8.90842 | 0.085036 |
| 8.95865 | 0.085364 |
| 9.01023 | 0.084876 |
| 9.0613 | 0.084853 |
| 9.11202 | 0.085181 |
| 9.16302 | 0.085624 |
| 9.21373 | 0.085517 |
| 9.26473 | 0.084205 |
| 9.31552 | 0.0848 |
| 9.36625 | 0.084243 |
| 9.41725 | 0.085715 |
| 9.46798 | 0.08583 |
| 9.51895 | 0.083884 |
| 9.56995 | 0.08583 |
| 9.62092 | 0.085746 |
| 9.67192 | 0.084395 |
| 9.72288 | 0.085936 |
| 9.77362 | 0.086364 |
| 9.82435 | 0.085631 |
| 9.87508 | 0.08628 |
| 9.92555 | 0.08496 |
| 9.97657 | 0.086135 |
| 10.0273 | 0.086264 |
| 10.0785 | 0.086089 |
| 10.1295 | 0.085845 |

| Time [min] | 10 nM |
| --- | --- |
| 0 | 0.0008228 |
| 0.050983 | 0.0001898 |
| 0.101734 | -0.0006722 |
| 0.15245 | 0.0009598 |
| 0.203417 | -6.22e-05 |
| 0.254167 | -0.0005962 |
| 0.305151 | 0.0003418 |
| 0.355883 | -0.0007262 |
| 0.406616 | -0.0006042 |
| 0.4576 | -0.0007792 |
| 0.508566 | -0.0011152 |
| 0.559549 | -0.0006652 |
| 0.610533 | -0.0007182 |
| 0.661783 | -0.0003982 |
| 0.712801 | 0.0002738 |
| 0.763782 | 0.0003878 |
| 0.814484 | 0.0004258 |
| 0.865484 | -5.42e-05 |
| 0.91645 | -0.0003292 |
| 0.9674 | -0.0004742 |
| 1.01838 | -0.0007492 |
| 1.06938 | -0.0006722 |
| 1.12037 | -0.0008172 |
| 1.17135 | -0.0005812 |
| 1.22208 | 0.0004188 |
| 1.27282 | -0.0001462 |
| 1.32352 | 0.0007618 |
| 1.37427 | 0.0019598 |
| 1.42528 | 0.0029588 |
| 1.476 | 0.0044088 |
| 1.52725 | 0.0053318 |
| 1.57798 | 0.0063768 |
| 1.62868 | 0.0075978 |
| 1.67942 | 0.0083458 |
| 1.7304 | 0.0096958 |
| 1.7814 | 0.0102298 |
| 1.8321 | 0.0117178 |
| 1.88283 | 0.0128088 |
| 1.93383 | 0.0135178 |
| 1.9848 | 0.0148308 |
| 2.03578 | 0.0155098 |
| 2.08678 | 0.0168748 |
| 2.13777 | 0.0172108 |
| 2.18875 | 0.0186378 |
| 2.23948 | 0.0198428 |
| 2.2905 | 0.0208808 |
| 2.3412 | 0.0218188 |
| 2.39192 | 0.0225358 |
| 2.44265 | 0.0236348 |
| 2.4934 | 0.0244058 |
| 2.5441 | 0.0253968 |
| 2.59508 | 0.0265948 |
| 2.64582 | 0.0274418 |
| 2.69653 | 0.0284638 |
| 2.74752 | 0.0288228 |
| 2.79825 | 0.0304708 |
| 2.84897 | 0.0313178 |
| 2.89968 | 0.0324388 |
| 2.95013 | 0.0329048 |
| 3.00138 | 0.0338348 |
| 3.05237 | 0.0350178 |
| 3.10337 | 0.0359788 |
| 3.15408 | 0.0366128 |
| 3.20508 | 0.0374668 |
| 3.2558 | 0.0385578 |
| 3.30678 | 0.0388778 |
| 3.35752 | 0.0397018 |
| 3.40823 | 0.0408538 |
| 3.45922 | 0.0414188 |
| 3.50995 | 0.0423568 |
| 3.56068 | 0.0434178 |
| 3.61167 | 0.0440358 |
| 3.66265 | 0.0453708 |
| 3.71338 | 0.0460728 |
| 3.76435 | 0.0465768 |
| 3.8151 | 0.0475608 |
| 3.86582 | 0.0482318 |
| 3.9168 | 0.0491168 |
| 3.96778 | 0.0493988 |
| 4.0185 | 0.0507728 |
| 4.06922 | 0.0507728 |
| 4.12022 | 0.0517028 |
| 4.1712 | 0.0522598 |
| 4.22218 | 0.0536258 |
| 4.27292 | 0.0541068 |
| 4.32363 | 0.0546028 |
| 4.37433 | 0.0556098 |
| 4.42532 | 0.0562578 |
| 4.4763 | 0.0567008 |
| 4.52728 | 0.0576618 |
| 4.57802 | 0.0586388 |
| 4.62872 | 0.0591568 |
| 4.67972 | 0.0596298 |
| 4.7307 | 0.0602868 |
| 4.78168 | 0.0611938 |
| 4.83267 | 0.0617968 |
| 4.88363 | 0.0629258 |
| 4.93462 | 0.0630408 |
| 4.98558 | 0.0630478 |
| 5.03658 | 0.0639718 |
| 5.0873 | 0.0644668 |
| 5.13828 | 0.0652688 |
| 5.18925 | 0.0664738 |
| 5.24023 | 0.0660468 |
| 5.29123 | 0.0674048 |
| 5.3422 | 0.0683278 |
| 5.39317 | 0.0679618 |
| 5.4442 | 0.0690828 |
| 5.49515 | 0.0694108 |
| 5.54613 | 0.0697468 |
| 5.59687 | 0.0713868 |
| 5.64758 | 0.0709068 |
| 5.69857 | 0.0722948 |
| 5.74955 | 0.0720508 |
| 5.80055 | 0.0721958 |
| 5.85153 | 0.0727828 |
| 5.90252 | 0.0740648 |
| 5.9535 | 0.0743628 |
| 6.00422 | 0.0746368 |
| 6.05495 | 0.0747438 |
| 6.10592 | 0.0750648 |
| 6.15692 | 0.0752548 |
| 6.20763 | 0.0758728 |
| 6.25835 | 0.0768188 |
| 6.30907 | 0.0766358 |
| 6.36007 | 0.0767358 |
| 6.41105 | 0.0770408 |
| 6.46207 | 0.0771628 |
| 6.51302 | 0.0777578 |
| 6.564 | 0.0778718 |
| 6.61473 | 0.0785968 |
| 6.66547 | 0.0785358 |
| 6.71645 | 0.0779868 |
| 6.76743 | 0.0785208 |
| 6.81817 | 0.0789558 |
| 6.86942 | 0.0794968 |
| 6.9204 | 0.0796498 |
| 6.97138 | 0.0799698 |
| 7.0221 | 0.0798178 |
| 7.07283 | 0.0794358 |
| 7.12355 | 0.0802298 |
| 7.17427 | 0.0799858 |
| 7.22523 | 0.0803668 |
| 7.27622 | 0.0803058 |
| 7.32722 | 0.0805578 |
| 7.3782 | 0.0804428 |
| 7.42918 | 0.0811448 |
| 7.47995 | 0.0810998 |
| 7.5309 | 0.0821518 |
| 7.58165 | 0.0822898 |
| 7.63237 | 0.0815648 |
| 7.68337 | 0.0811298 |
| 7.73408 | 0.0818318 |
| 7.7848 | 0.0821448 |
| 7.83552 | 0.0818778 |
| 7.8865 | 0.0816718 |
| 7.93723 | 0.0824578 |
| 7.98823 | 0.0821448 |
| 8.03893 | 0.0826478 |
| 8.08967 | 0.0826708 |
| 8.14065 | 0.0821518 |
| 8.19165 | 0.0829308 |
| 8.24263 | 0.0830978 |
| 8.29337 | 0.0830068 |
| 8.34433 | 0.0822588 |
| 8.39532 | 0.0827398 |
| 8.44603 | 0.0824268 |
| 8.49702 | 0.0836938 |
| 8.548 | 0.0833198 |
| 8.59898 | 0.0828238 |
| 8.64942 | 0.0828538 |
| 8.70095 | 0.0824958 |
| 8.75192 | 0.0830828 |
| 8.80292 | 0.0830528 |
| 8.85363 | 0.0833578 |
| 8.90435 | 0.0841818 |
| 8.95535 | 0.0837318 |
| 9.00607 | 0.0841968 |
| 9.05677 | 0.0830068 |
| 9.1075 | 0.0841438 |
| 9.15822 | 0.0836018 |
| 9.2092 | 0.0836708 |
| 9.25992 | 0.0843418 |
| 9.31091 | 0.0838308 |
| 9.36188 | 0.0843118 |
| 9.41287 | 0.0842278 |
| 9.46358 | 0.0842958 |
| 9.51432 | 0.0835638 |
| 9.56555 | 0.0839148 |
| 9.61623 | 0.0843498 |
| 9.66723 | 0.0838918 |
| 9.71822 | 0.0842428 |
| 9.76893 | 0.0847388 |
| 9.81992 | 0.0839828 |
| 9.87115 | 0.0844558 |
| 9.92188 | 0.0842578 |
| 9.97285 | 0.0842278 |
| 10.0238 | 0.0833348 |
| 10.0748 | 0.0844948 |
| 10.1255 | 0.0838538 |

| Time [min] | 10 nM | 5 nM |
| --- | --- | --- |
| 0 | -0.00050105 | -0.0003638 |
| 0.050983 | -0.00105805 | -0.0006918 |
| 0.101984 | 0.00152095 | -0.0002568 |
| 0.152966 | -0.00090505 | 0.0001932 |
| 0.203967 | -0.00079105 | -0.0003638 |
| 0.25495 | 0.00110095 | 4.82e-05 |
| 0.305934 | -0.00041705 | -0.0003568 |
| 0.356666 | -0.00124105 | -0.0004938 |
| 0.407667 | -0.00095105 | -0.0003258 |
| 0.4584 | -0.00114905 | -0.0002188 |
| 0.5094 | -0.00105805 | -0.0001958 |
| 0.560384 | 0.00058995 | -0.0005318 |
| 0.611133 | -0.00126405 | -0.0006998 |
| 0.66185 | -0.00118805 | -0.0006918 |
| 0.712583 | 0.00084195 | -0.0004938 |
| 0.763317 | -0.00114905 | -0.0008298 |
| 0.81395 | -0.00049305 | 1.72e-05 |
| 0.8652 | -0.00142405 | -0.0009208 |
| 0.915933 | 0.00010195 | 0.0002842 |
| 0.9669 | 0.00086495 | 3.32e-05 |
| 1.0179 | -0.00184405 | -0.0002568 |
| 1.06888 | 0.00067395 | 0.0005902 |
| 1.12015 | -0.00132505 | 0.0014902 |
| 1.17113 | -0.00166805 | 0.0005212 |
| 1.22187 | -0.00161505 | 0.0005592 |
| 1.27288 | -0.00102005 | 0.0012612 |
| 1.32388 | 0.00142195 | 0.0020922 |
| 1.37458 | 0.00082695 | 0.0019552 |
| 1.42533 | 0.00173495 | 0.0029242 |
| 1.47605 | 0.00293995 | 0.0049842 |
| 1.52677 | 0.00396995 | 0.0042592 |
| 1.57775 | 0.00316095 | 0.0041452 |
| 1.62875 | 0.00316095 | 0.0049232 |
| 1.67972 | 0.00551095 | 0.0047632 |
| 1.73048 | 0.00454995 | 0.0054192 |
| 1.78122 | 0.00430595 | 0.0057702 |
| 1.8322 | 0.00726595 | 0.0065862 |
| 1.88318 | 0.00628895 | 0.0065942 |
| 1.93417 | 0.00644895 | 0.0072192 |
| 1.98515 | 0.00692295 | 0.0075632 |
| 2.0359 | 0.00722795 | 0.0077762 |
| 2.08688 | 0.00721995 | 0.0080132 |
| 2.13788 | 0.00781495 | 0.0085622 |
| 2.1886 | 0.00844095 | 0.0087612 |
| 2.23933 | 0.00853195 | 0.0090122 |
| 2.29005 | 0.00996695 | 0.0100042 |
| 2.34078 | 0.00981395 | 0.0097982 |
| 2.3915 | 0.010333 | 0.0100272 |
| 2.44247 | 0.011493 | 0.0097602 |
| 2.49345 | 0.011386 | 0.0116752 |
| 2.54418 | 0.011401 | 0.0116832 |
| 2.5949 | 0.012202 | 0.0116602 |
| 2.64588 | 0.012202 | 0.0122622 |
| 2.69688 | 0.012599 | 0.0118432 |
| 2.74738 | 0.012675 | 0.0124302 |
| 2.79893 | 0.014003 | 0.0127432 |
| 2.84992 | 0.013873 | 0.0130792 |
| 2.90092 | 0.014231 | 0.0135062 |
| 2.95193 | 0.015246 | 0.0134912 |
| 3.0029 | 0.01475 | 0.0142542 |
| 3.05335 | 0.015216 | 0.0136972 |
| 3.10487 | 0.015719 | 0.0144442 |
| 3.1556 | 0.015643 | 0.0144142 |
| 3.20658 | 0.016406 | 0.0154592 |
| 3.25783 | 0.016314 | 0.0156962 |
| 3.30882 | 0.016772 | 0.0151392 |
| 3.35957 | 0.016894 | 0.0156732 |
| 3.41057 | 0.018878 | 0.0160472 |
| 3.46103 | 0.019648 | 0.0167942 |
| 3.51228 | 0.019748 | 0.0165042 |
| 3.56302 | 0.019046 | 0.0175572 |
| 3.614 | 0.019069 | 0.0171302 |
| 3.66472 | 0.019839 | 0.0178172 |
| 3.71542 | 0.021426 | 0.0182972 |
| 3.7656 | 0.020747 | 0.0186642 |
| 3.81685 | 0.021022 | 0.0183972 |
| 3.86787 | 0.021632 | 0.0191592 |
| 3.91885 | 0.020564 | 0.0188542 |
| 3.96985 | 0.021693 | 0.0199152 |
| 4.02083 | 0.021777 | 0.0201742 |
| 4.07185 | 0.022822 | 0.0204952 |
| 4.12282 | 0.022326 | 0.0208532 |
| 4.1738 | 0.023242 | 0.0213112 |
| 4.22478 | 0.022479 | 0.0214562 |
| 4.27578 | 0.023944 | 0.0219982 |
| 4.32652 | 0.023837 | 0.0220742 |
| 4.37753 | 0.02463 | 0.0217762 |
| 4.42825 | 0.023806 | 0.0227682 |
| 4.47925 | 0.026973 | 0.0234552 |
| 4.53023 | 0.024096 | 0.0227452 |
| 4.5812 | 0.024806 | 0.0228062 |
| 4.63195 | 0.025607 | 0.0242332 |
| 4.68268 | 0.02592 | 0.0241412 |
| 4.73365 | 0.026507 | 0.0250112 |
| 4.78465 | 0.02695 | 0.0246602 |
| 4.83538 | 0.027232 | 0.0253092 |
| 4.88612 | 0.02756 | 0.0255152 |
| 4.9371 | 0.027827 | 0.0257972 |
| 4.98785 | 0.027606 | 0.0262012 |
| 5.03858 | 0.028201 | 0.0268272 |
| 5.08957 | 0.028361 | 0.0264232 |
| 5.14057 | 0.029567 | 0.0281012 |
| 5.19158 | 0.030024 | 0.0274222 |
| 5.24228 | 0.029734 | 0.0286202 |
| 5.29327 | 0.03078 | 0.0283832 |
| 5.34427 | 0.031031 | 0.0288182 |
| 5.395 | 0.030803 | 0.0281702 |
| 5.44572 | 0.030978 | 0.0291922 |
| 5.49643 | 0.031916 | 0.0288872 |
| 5.54742 | 0.031817 | 0.0288412 |
| 5.59842 | 0.031588 | 0.0296272 |
| 5.64942 | 0.032611 | 0.0295132 |
| 5.70015 | 0.031642 | 0.0295202 |
| 5.75087 | 0.032717 | 0.0297872 |
| 5.8016 | 0.032527 | 0.0299022 |
| 5.85233 | 0.033107 | 0.0300622 |
| 5.90307 | 0.033763 | 0.0299472 |
| 5.9538 | 0.033381 | 0.0302682 |
| 6.00478 | 0.033488 | 0.0303372 |
| 6.0555 | 0.033816 | 0.0302602 |
| 6.10622 | 0.03374 | 0.0303522 |
| 6.15695 | 0.0338009 | 0.0301762 |
| 6.2077 | 0.033839 | 0.0309772 |
| 6.25867 | 0.034289 | 0.0310002 |
| 6.30942 | 0.034068 | 0.0306952 |
| 6.36038 | 0.034953 | 0.0315042 |
| 6.41112 | 0.0348 | 0.0309542 |
| 6.4621 | 0.035174 | 0.0310382 |
| 6.51308 | 0.034755 | 0.0308022 |
| 6.5638 | 0.034785 | 0.0315952 |
| 6.61457 | 0.03638 | 0.0308322 |
| 6.66555 | 0.034495 | 0.0317402 |
| 6.71655 | 0.036082 | 0.0312672 |
| 6.76752 | 0.035243 | 0.0318782 |
| 6.81825 | 0.035159 | 0.0315572 |
| 6.86923 | 0.035006 | 0.0321222 |
| 6.92023 | 0.035632 | 0.0317402 |
| 6.97122 | 0.035632 | 0.0318932 |
| 7.02193 | 0.034869 | 0.0318932 |
| 7.07267 | 0.03535 | 0.0326022 |
| 7.1234 | 0.036532 | 0.0323662 |
| 7.17415 | 0.036174 | 0.0324802 |
| 7.22542 | 0.036135 | 0.0321222 |
| 7.27615 | 0.035388 | 0.0327472 |
| 7.32713 | 0.036853 | 0.0323812 |
| 7.37812 | 0.037143 | 0.0324042 |
| 7.42915 | 0.036364 | 0.0323202 |
| 7.48013 | 0.036715 | 0.0327172 |
| 7.53115 | 0.036608 | 0.0324422 |
| 7.58215 | 0.038966 | 0.0325872 |
| 7.63288 | 0.03667 | 0.0331062 |
| 7.68333 | 0.0371499 | 0.0327702 |
| 7.73487 | 0.037516 | 0.0330912 |
| 7.78562 | 0.037898 | 0.0328542 |
| 7.83632 | 0.039393 | 0.0329382 |
| 7.88733 | 0.036944 | 0.0334042 |
| 7.93835 | 0.039904 | 0.0331292 |
| 7.9891 | 0.037082 | 0.0333732 |
| 8.03962 | 0.038737 | 0.0332512 |
| 8.09113 | 0.038035 | 0.0330602 |
| 8.14212 | 0.040004 | 0.0327552 |
| 8.19283 | 0.037722 | 0.0335712 |
| 8.24383 | 0.037539 | 0.0335102 |
| 8.29482 | 0.037982 | 0.0332892 |
| 8.34582 | 0.03815 | 0.0329762 |
| 8.39653 | 0.038302 | 0.0325572 |
| 8.44753 | 0.037852 | 0.0327862 |
| 8.49853 | 0.038005 | 0.0340982 |
| 8.54952 | 0.037761 | 0.0338462 |
| 8.60053 | 0.038302 | 0.0329232 |
| 8.65127 | 0.038035 | 0.0337242 |
| 8.70202 | 0.039332 | 0.0333812 |
| 8.75302 | 0.038371 | 0.0335792 |
| 8.80377 | 0.038447 | 0.0330832 |
| 8.85475 | 0.03789 | 0.0338772 |
| 8.9057 | 0.038096 | 0.0338612 |
| 8.95643 | 0.038394 | 0.0331442 |
| 9.00715 | 0.038012 | 0.0332812 |
| 9.05815 | 0.038295 | 0.0332202 |
| 9.10913 | 0.038417 | 0.0334492 |
| 9.16013 | 0.038173 | 0.0333652 |
| 9.21085 | 0.038882 | 0.0340832 |
| 9.26168 | 0.038066 | 0.0333042 |
| 9.31267 | 0.038195 | 0.0338842 |
| 9.36417 | 0.038287 | 0.0339992 |
| 9.41465 | 0.038424 | 0.0334112 |
| 9.46563 | 0.040736 | 0.0341662 |
| 9.5169 | 0.038333 | 0.0329842 |
| 9.56792 | 0.039378 | 0.0344032 |
| 9.61863 | 0.038028 | 0.0336632 |
| 9.66937 | 0.039157 | 0.0338922 |
| 9.72012 | 0.037547 | 0.0338382 |
| 9.77112 | 0.040767 | 0.0331752 |
| 9.8221 | 0.03876 | 0.0339072 |
| 9.87308 | 0.038432 | 0.0333582 |
| 9.92408 | 0.038173 | 0.0334262 |
| 9.97507 | 0.038935 | 0.0336172 |
| 10.0261 | 0.038127 | 0.0331752 |
| 10.0771 | 0.037905 | 0.0338082 |
| 10.128 | 0.038127 | 0.0335942 |

**Figure S6.** Raw SPR sensorgrams for TcdA injections (t = 1.2 to 9.5 min) over SA1 functionalized gold sensors. SPR response was normalized with sensor and channel dependent baselines, established for > 5 min prior to injection.

5. **Modeling TcdA Enzyme Kinetics with Competitive Inhibition by SA1**

Based on the design and MD simulations of SA1 in the UDP-Glucose binding pocket of the TcdA GTD, SA1 likely acts as a competitive inhibitor to UDP-Glucose binding, and thus glucosyltransferase activity. Binding studies performed in the absence of UDP-Glucose reproducibly demonstrated TcdA:SA1 binding. Based on MD simulations of SA1:TcdA GTD and the crystal structure (PDB 3SRZ) of UDP-Glucose:TcdA GTD showing that SA1 and UDP-Glucose independently occupy the same binding pocket, it is unlikely that a stable TcdA•SA1•UDP-Glucose complex forms, making non-competitive or uncompetitive inhibition models less likely. The competitive inhibition model is described in Equation S5.

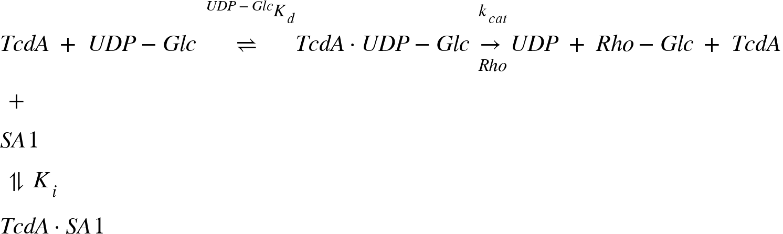
(S5) where ^UDP-Glc^K_D_ is the dissociation constant for TcdA:UDP-Glucose binding, k_cat_ [1/time] is the glucosylation rate constant, K_i_ [mol/length^3^] is the inhibition constant for TcdA:SA1. Under these conditions, the inhibition constant, K_i_, is the dissociation constant for TcdA:SA1, K_D_. The modified Michaelis-Menten equation for competitive inhibition is as follows,

$$\begin{aligned} v=\frac{k_{cat}\left[ TcdA \right]\left[ UDP-Glc \right]}{K_{M}\left( 1+\frac{\left[ SA1 \right]}{K_{i}} \right)+\left[ UDP-Glc \right]}\#\left( S6 \right) \end{aligned}$$

where v is the reaction rate [mol/length^3^/time] and K_M_ [mol/length^3^] is the Michaelis constant. Without inhibition, the Michaelis-Menten equation simplifies to Equation S7.

$$\begin{aligned} v=\frac{k_{cat}\left[ TcdA \right]\left[ UDP-Glc \right]}{K_{M}+\left[ UDP-Glc \right]}\#\left( S7 \right) \end{aligned}$$

**Table S4.** Kinetic parameters for glucosylation of Rho family proteins by TcdA.

| **Kinetic Parameter** | **Measured Value** | **Reference** |
| --- | --- | --- |
| ^UDP-Glc^K_D_ | 45 μM | ^17^ |
| k_cat_ | 0.0020 1/s | ^18^ |
| K_M_ | 4.5 μM | ^18^ |
| V_max_ | 0.00020 μM/s | ^18^ |
| K_i_ (^SA1^K_D_) | 56.1 ± 29.8 nM | This study |

**Table S5.** Physiological concentrations of key species.

| **Physiological Condition** | **Value** | **Reference** |
| --- | --- | --- |
| [UDP-Glucose] | 92 μM | ^19^ |
| [TcdA] | 30 pM | ^20^ |

Using parameter values from Table S4 and physiological conditions from Table S5, Equations S6 and S7 were solved for concentrations of SA1 from 0 to 1 mM. The theoretical IC_50_ of SA1 is 1.2 ± 0.6 μM. The dissociation constant for SA1:TcdA, ^SA1^K_D_, is approximately 1000-fold smaller than for UDP-Glucose:TcdA, ^UDP-Glc^K_D_, and the association constant, ^SA1^k_a_, is also quite small, favoring fast binding. However, without more detailed measurements of UDP-Glucose:TcdA binding (^UDP-Glc^k_a_) it is not possible to directly describe how SA1 and UDP-Glucose compete for the active site of TcdA GTD. The true IC_50_ in a physiologically relevant context likely lies between the theoretical IC_50_ of SA1 (~1 μM) and the concentration used in the TEER assays (1 mM). The IC_50_ in physiologically relevant, and thus more complex conditions will not only capture the competitive inhibition kinetics in the local environment (cell cytosol), but also SA1’s transport to the site of action and its degradation. Future work developing a dose-response curve for SA1 in the TEER assay will better inform the therapeutic dosage.

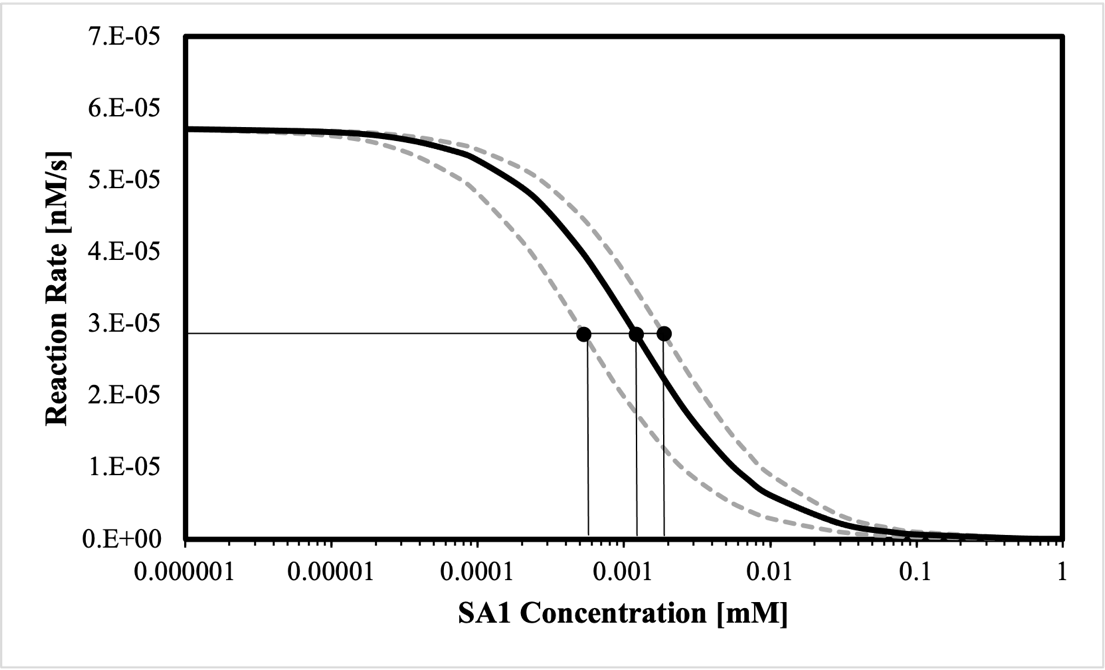

**Figure S7.** TcdA inhibition by SA1. 95% confidence interval for ^SA1^K_D_ shown as grey dashed lines. Theoretical SA1 IC_50_ = 1.2 ± 0.6 μM.

**References**

1.	X. Xiao, C. K. Hall, P. F. Agris, The design of a peptide sequence to inhibit HIV replication: A search algorithm combining Monte Carlo and self-consistent mean field techniques. *J. Biomol. Struct. and Dyn*. **32**, 1523-1536 (2014).

2.	X. Xiao, P. F. Agris, C. K. Hall. Introducing folding stability into the score function for computational design of RNA-binding peptides boosts the probability of success. *Proteins: Struct. Funct. Genet*. **84**, 700-711 (2016).

3.	X. Xiao, M. E. Hung, J. N. Leonard, C. K. Hall, Adding energy minimization strategy to peptide-design algorithm enables better search for RNA-binding peptides: Redesigned λ N peptide binds boxB RNA. *J. Comp. Chem.* **37**, 2423-2435 (2016).

4.	X. Xiao, Y. Wang, J. N. Leonard, C. K. Hall. Extended Concerted Rotation Technique Enhances the Sampling Efficiency of the Computational Peptide-Design Algorithm. *J. Chem. Theory Comput*. **13**, 5709-5720 (2017).

5.	X. Xiao, B. Zhao, P. F. Agris, C. K. Hall. Simulation study of the ability of a computationally-designed peptide to recognize target tRNALys3 and other decoy tRNAs. *Protein Sci*. **25**, 2243-2255 (2016).

6.	X. Xiao, P. F. Agris, C. K. Hall, Designing peptide sequences in flexible chain conformations to bind RNA: A search algorithm combining Monte Carlo, self-consistent mean field, and concerted rotation techniques. *J. Chem. Theory Comput*. **11**, 740-752 (2015).

7.	K. Day, *et al*., Discovery and Evaluation of Peptide Ligands for Selective Adsorption and Release of Cas9 Nuclease on Solid Substrates. *Bioconjugate Chem*. **30**, 3057-3068 (2019).

8.	S Saberi-Bosari, *et al*., Affordable Microfluidic Bead-Sorting Platform for Automated Selection of Porous Particles Functionalized with Bioactive Compounds. *Scientific Reports*. **9**, 7210 (2019).

9.	M. T. Ok, *et al*., A Leaky Colon Model Reveals Uncoupled Apical/Basal Cytotoxicity in Early Clostridioides difficile Toxin Exposure. *bioRxiv* 2022. Available at <https://doi.org/10.1101/2022.10.13.511617> (accessed 2 November 2022).

10.	X. Xiao, *et al*. In Silico Identification and Experimental Validation of Peptide-Based Inhibitors Targeting Clostridium difficile Toxin A. *ACS Chem. Biol*. **17**, 118-128 (2022).

11.	S. J. Abdeen, R. J. Swett, A. L. Feig, Peptide inhibitors targeting clostridium difficile toxins A and B. *ACS Chem. Biol*. **5**, 1097-1103 (2010).

12.	N. Islam, F. Shen, P. V. Gurgel, O. J. Rojas, R. G. Carbonell, Dynamic and equilibrium performance of sensors based on short peptide ligands for affinity adsorption of human IgG using surface plasmon resonance. *Biosens. Bioelectron*. **58**, 380-387 (2014).

13.	V. Humblot, *et al*., The antibacterial activity of Magainin I immobilized onto mixed thiols Self-Assembled Monolayers. *Biomaterials*. **30**, 3503-3512 (2009).

14.	N. Islam, P. V. Gurgel, O. J. Rojas, R. G. Carbonell. Use of a Branched Linker for Enhanced Biosensing Properties in IgG Detection from Mixed Chinese Hamster Ovary Cell Cultures. *Bioconjugate Chem*. **30**, 815-825 (2019).

15.	F. Cheng, L. J. Gamble, D. G. Castner. XPS, TOF-SIMS, NEXAFS, and SPR characterization of nitrilotriacetic acid-terminated self-assembled monolayers for controllable immobilization of proteins. *Anal. Chem*. **80**, 2564-2573 (2008).

16.	S. Aoyagi, *et al*., Evaluation of Time-of-Flight Secondary Ion Mass Spectrometry Spectra of Peptides by Random Forest with Amino Acid Labels: Results from a Versailles Project on Advanced Materials and Standards Interlaboratory Study. *Anal. Chem*. **93**, 4191-4197 (2021).

17.	S. Bhattacharyya, A. Kerzmann, A. L. Feig, Fluorescent analogs of UDP-glucose and their use in characterizing substrate binding by toxin A from Clostridium difficile. *Eur. J. Biochem*. **269**, 3425-3432 (2002).

18.	J. W Loughney, *et al*., Development of a non-radiolabeled glucosyltransferase activity assay for C. difficile toxin A and B using ultra performance liquid chromatography. *J. Chromatogr. A*. **1498**, 169-175 (2017).

19.	M. R. Laughlin, W. A. Petit, J. M. Dizon, R. G. Shulman, E. J. Barrett. NMR measurements of in vivo myocardial glycogen metabolism. *J. Biol. Chem*. **263**, 2285-2291 (1988).

20.	L. Song, *et al*., Development and Validation of Digital Enzyme-Linked Immunosorbent Assays for Ultrasensitive Detection and Quantification of Clostridium difficile Toxins in Stool. *J. Clin. Microbiol*. **53**, 3204-3212 (2015).
